# Default Mode Network Connectivity Is Associated with Capture by a Distractor Dimension Carrying Learned Spatial Regularity

**DOI:** 10.64898/2026.04.22.720249

**Authors:** Siyi Chen, Hermann J. Müller, Zhuanghua Shi

**Affiliations:** Neuro-Cognitive Psychology, Department of Psychology, LMU Munich, Leopoldstr. 13, 80802 Munich, Germany; NeuroImaging Core Unit Munich (NICUM), Department of Psychiatry and Psychotherapy, LMU Munich, Nußbaumstraße 7, 80336 Munich, Germany

**Keywords:** attentional capture, default mode network, statistical learning, functional connectivity, individual differences

## Abstract

Salient distractors interfere with visual search, but observers can learn to filter out distractors that occur frequently and in predictable locations. We examined whether within-default-mode-network (DMN) task-run connectivity was associated with distractor capture and its location-dependent modulation in task-fMRI data from two groups in which either orientation or color distractors carried a spatial-probability bias. Higher within-DMN connectivity was associated with greater distractor capture by the biased distractor dimension, which was also the most frequent distractor type, whereas capture by the unbiased dimension was essentially unrelated to connectivity. A hierarchical trial-level model also showed stronger DMN– distractor-capture coupling for the biased than for the unbiased dimension when both were evaluated under the same frequent-/rare-region weighting. DMN connectivity was also associated with spatial-probability learning, indexed by the rare-minus-frequent reaction-time difference on biased-dimension trials, but this association was closely tied to overall capture. When the physical distractor dimension was held constant, orientation showed stronger DMN–distractor-capture coupling when it carried the spatial bias; in contrast, the corresponding color comparison remained uncertain. The broader network analyses provided converging support for this DMN–distractor-capture coupling pattern. Higher DMN connectivity was thus not associated with poorer performance in general, but with less effective filtering of the distractor that the task set was tuned to suppress. This pattern is consistent with a DMN state that interferes with implementing the optimal task set.

## Introduction

Salient but irrelevant stimuli often draw attention away from the current task, slowing responses to task-relevant targets. This phenomenon, of *attentional capture*, illustrates a central limit of cognitive control [1, 2, 3, 4]. Yet capture is not uniform across observers. Some individuals remain strongly affected by salient distractors, whereas others show greater resistance to interference, consistent with individual differences in attentional control [5, 6]. Understanding the neural correlates of these individual differences is important because distractor vulnerability is not simply a property of the stimulus; it may also depend on how effectively observers implement a task set that keeps known distractors from competing with the target, and this implementation may vary with the state of large-scale brain networks.

One source of such prior experience is statistical learning. Work on contextual cueing first showed that implicitly learned spatial context can guide attention during visual search [7, 8]. Related distractor-location learning studies show that, when distractors appear more often in a particular location or region, observers can (typically implicitly) learn this spatial regularity and reduce capture from that high-probability region [9, 10, 11, 12, 13]. How effectively a distractor can be filtered also depends on its relation to the target. According to the dimension-weighting account, distractors defined in a different dimension from the target can be down-weighted at the level of the whole dimension, whereas distractors defined in the target dimension cannot be filtered in this way without also impairing target selection [2, 13, 14, 15]. Dimension-based weighting and distractor-location learning thus jointly define the optimal task set for a given search environment. This probability-driven modulation of distractor suppression provides a useful model for how attentional control is shaped by selection history. However, observers differ in the degree to which they benefit from such regularities. Some show clear reductions in capture by distractors at high-vs. low probability locations, whereas others show little effect of distractor-location learning. These differences raise an individual-differences question: is variability in learned distractor vulnerability related only to local modulation of attentional selection, or is it also associated with broader differences in brain-network organization?

The default-mode network (DMN) is a plausible candidate for such an individual-difference marker. The DMN was initially characterized as a set of midline, medial temporal, and lateral parietal regions that reduce activity during externally focused tasks [16], and subsequent work linked it to internally oriented cognition, self-generated thought, contextual memory, and mind wandering [17, 18, 19]. DMN engagement during task performance has been associated with slower responses, increased errors, and attentional lapses [20, 21]. Prior work also suggests that attentional performance depends partly on how strongly the DMN is functionally separated from task-positive attention networks: greater DMN–attention-network segregation has been associated with lower reaction-time (RT) variability [22]. These findings suggest that a strongly engaged or internally coupled DMN may interfere with maintaining the externally oriented task set that efficient search requires.

Connectivity-based prediction studies further highlight the relevance of DMN organization for attentional performance. Rosenberg and colleagues [23], for example, showed that whole-brain functional connectivity predicts individual differences in sustained attention, with DMN-related connections forming an important part of the predictive network. This finding fits with broader evidence that attention depends on coordinated interactions among externally oriented task-positive systems, reorienting/salience systems, and the default-mode system [22, 24].

However, most of this work has focused on sustained-attention contexts, where poor performance is typically interpreted as a lapse of task focus or a failure to maintain goal-directed control over time. Attentional capture poses a related but more specific question. Capture is a brief, stimulus-triggered event, reflecting momentary competition between target and distractor information. How strongly a distractor competes, however, depends on the task set in place when it appears: down-weighting of the distractor dimension and suppression of the region where distractors often appear are both adjusted over many trials to the statistics of the task environment [9, 13]. A sustained network state could therefore influence capture by affecting how well this task set is implemented. It remains unclear whether DMN connectivity, previously linked mainly to sustained attention and internally directed cognition, also relates to individual vulnerability to salient distractors during visual selection, and in particular to capture by the distractors that the task set is tuned to filter out.

We tested this question by reanalyzing an existing task-fMRI dataset originally designed to examine statistical learning of distractor suppression in visual search [25]. Two groups of participants searched for an orientation-defined target (of a relatively low salience) while salient distractors appeared with unequal spatial probabilities. In the same-dimension spatial-bias (SS) group, the spatial regularity was assigned to orientation distractors that shared the task-critical (i.e., target-defining) dimension; in the different-dimension spatial-bias (DS) group, it was assigned to color-defined distractors (with the orientation and color distractors matched for saliency). Thus, the physical distractor dimension carrying the spatial bias was counterbalanced across groups, allowing us to examine whether DMN connectivity followed assigned bias status or physical distractor dimension. The dimension that carried the spatial bias was also presented more frequently within each group, making it the main distractor the task set had to filter; within-group spatial regularity and dimensional exposure were therefore not manipulated independently.

Our primary question was whether mean within-DMN task-run connectivity was associated with distractor capture and its location-dependent modulation in task-fMRI data from the two groups in which either orientation or color distractors carried a spatial-probability bias. We first addressed this question using participant-level capture scores. We then tested the same question at the trial level using a hierarchical model that distinguished distractor-absent, biased-frequent, biased-rare, and unbiased trials. This model estimated how distractor-related RT costs varied with DMN connectivity, while accounting for repeated observations within participants and without requiring equal trial counts across conditions. Biased and unbiased coupling were then compared under a common 50/50 frequent/rare-region weighting, standardizing the spatial-region mixture. We additionally asked whether the observed coupling followed the dimension assigned the spatial bias or the physical distractor dimension. To address this, we examined the coupling between DMN connectivity and attentional capture by orientation– and color-defined distractors within each group and compared the SS and DS groups while holding physical distractor dimension constant. Spatial-probability learning was examined as the spatial gradient of the same capture effect, defined as the RT difference between distractors in the rare minus the frequent region (rare-minus-frequent RT difference) on biased-dimension trials, which equals the difference in capture between the two regions. A corresponding hierarchical model tested whether this location-dependent effect covaried with DMN connectivity and whether that coupling changed across scanner runs. Finally, we examined whether the DMN associations were specific to this network or reflected broader patterns of functional connectivity by comparing within-and between-network connectivity associations. These broader network analyses were exploratory and provided context for interpreting the focal DMN findings.

## Methods

### Participants

The source dataset included 35 healthy adults recruited from the local community. All reported normal or corrected-to-normal vision and no history of neurological or psychiatric disorders, gave written informed consent, and participated under a protocol approved by the local ethics committee in accordance with the Declaration of Helsinki. Participants were randomly assigned to the same-dimension spatial-bias (SS) group or the different-dimension spatial-bias (DS) group.

The primary connectivity and brain–behavior analyses used data from 33 participants. Two recruited participants were excluded from these analyses: one because preprocessed blood-oxygen-level-dependent (BOLD) data were incomplete, and one because of excessive motion artifacts. Participants were excluded if they had both mean framewise displacement (FD) greater than 0.30 mm and more than 10% of task-run volumes with FD greater than 0.50 mm. One participant met both criteria and was excluded (mean FD = 0.31 mm; 14.7% of volumes with FD > 0.50 mm).

The final connectivity sample included 16 participants in the same-dimension spatial-bias group and 17 in the different-dimension spatial-bias group. Across the two groups, participants’ ages ranged between 22–41 years (M = 28.12, SD = 4.10; 21 females, 12 males); and the two groups did not differ reliably in age (p = .939) or sex distribution (p = .895).

### Task and Design

Participants performed a visual-search task in which they reported whether the notch of the uniquely tilted (i.e., orientation-defined) target was located at the top or the bottom by pressing the corresponding left-or right-hand response key with their respective index finger. The mapping between notch position (top/bottom) and response hand (left/right) was counterbalanced across participants.

Each trial began with a jittered fixation interval of 0.5 or 5.5 s (with equal probability), followed by a 1-s search display. Responses were accepted until 1.9 s after display onset, and the inter-trial interval varied randomly between 3 and 4 s. Displays contained 26 blue-cyan bar-like items (programmed RGB: 0, 140, 209) arranged on three concentric rings around the fixation point (Figure 1). The concentric-ring arrangement followed previous spatial distractor-learning paradigms [9, 13]. Target and distractor positions were restricted to the middle ring to maintain comparable local stimulus configurations across locations, thereby minimizing differences in bottom-up salience arising from variation in feature contrast with surrounding items [26]. The target was the only item tilted 12° from vertical. The same-dimension singleton distractor was a bar-like tilted 45° from vertical (i.e., it was more salient than the 12°-tilted target), whereas the different-dimension singleton distractor was an upright red item (matched in saliency to the orientation distractor). **Figure 1** illustrates the three display types used in the task.

**Figure 1.**
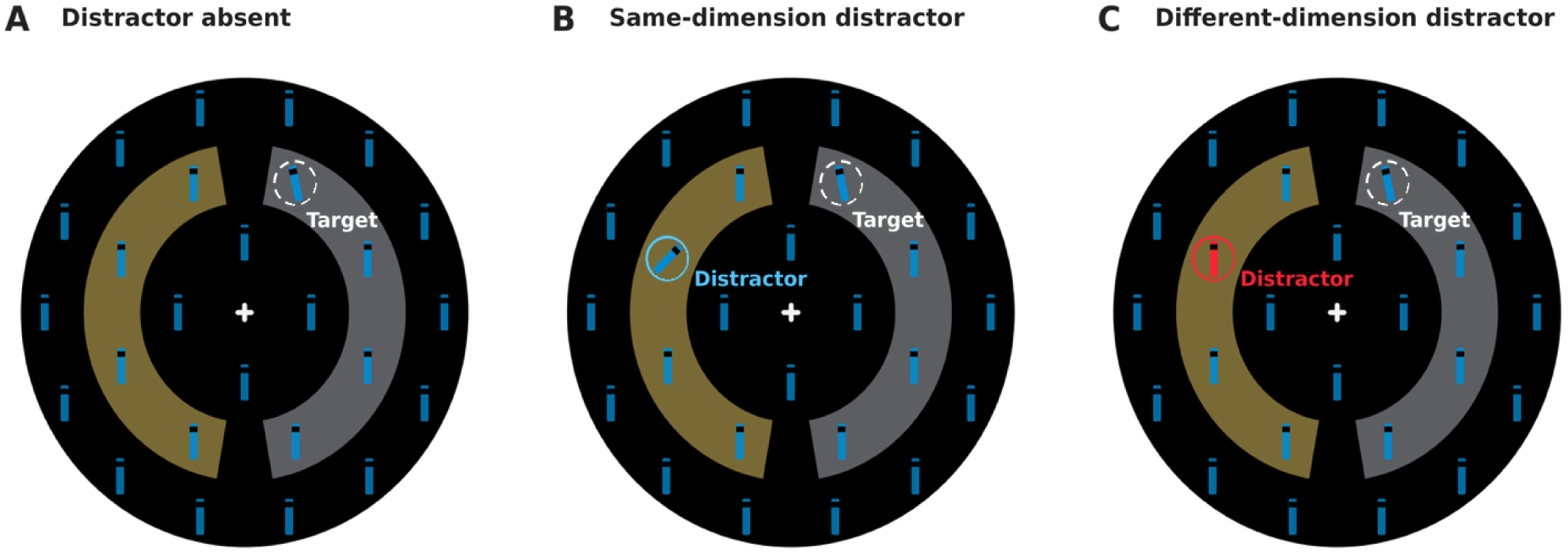
Visual-search displays and spatial-probability manipulation. Participants searched among 26 bar-like (‘i’-type) items arranged on three concentric rings for the unique target tilted 12° from vertical and reported whether its notch (the ‘i’-dot) was positioned at the top or the bottom. Non-target items and the target were blue-cyan (programmed RGB: 0, 140, 209). (**A**) Distractor-absent display. (**B**) Same-dimension distractor display, with a blue-cyan orientation singleton tilted 45°. (**C**) Different-dimension distractor display, with an upright red color singleton. Both groups received all three display types; orientation was the spatially biased distractor dimension in the SS group, and color in the DS group. Target and distractor positions were restricted to the eight middle-ring locations (to equalize feature contrast to adjacent items on the same ring and the inner and outer rings [26]), divided into two four-position lateral regions. Each 184-trial scanner run contained 32 distractor-absent trials (17%), 112 biased-dimension trials (61%) and 40 unbiased-dimension trials (22%); a distractor was therefore present on 152 trials. Within the biased dimension, 92 of 112 distractors occurred in the frequent region (82%) and 20 in the opposite rare region (18%). Unbiased-dimension distractors occurred equally in the two regions (20/40 per region). Yellow and gray sectors show one example of frequent/rare region assignment; the frequent side was counterbalanced across participants. Highlight circles, labels and shaded sectors are explanatory annotations, not participantvisible display elements. The panels are illustrative search displays, not screenshots or a trial-timing diagram.

The task followed the distractor-location probability-learning approach used in previous studies [9, 10, 11, 12], in which distractors occur more frequently in a particular location or region. Specifically, adopting the design of a recent study examining for differential carry-over effects in distractor suppression learning between distractors defined in the same vs. a different dimension to the target [13], one distractor dimension was assigned a participant-specific high-probability region on either the left or right side of the middle ring, while the other dimension had no spatial-probability bias. Target and distractor positions were restricted to the middle ring to maintain comparable local stimulus arrangements across locations (**Figure 1**). Distractors were absent on 17% of trials. Across all trials, distractors defined in the spatially biased (distractor) dimension appeared on 61% of trials; within these trials, the distractor appeared in the frequent region in approximately 82% of trials and in the opposite, rare region in 18% of trials. The other distractor dimension was spatially unbiased and appeared on 22% of trials, with locations distributed across the same middle-ring positions but without a high-/low-probability spatial bias. Which (left/right) region was frequent vs. rare to contain a spatially biased distractor was counterbalanced across participants. In the SS (same-dimension spatial-bias) group, the bias applied to orientation distractors; in the DS (different-dimension spatial-bias) group, it applied to color distractors. Participants completed six training blocks outside the scanner, comprising 276 trials, followed by two test runs inside the scanner, comprising 368 trials in total (184 trials per run). Each scanner run contained 32 distractor-absent trials, 40 unbiased-dimension trials (20 per side), and 112 biased-dimension trials, with the distractor occurring in 92 trials in the frequent region and in 20 trials in the rare region. Given this, assignment to the spatially biased dimension was accompanied by overall greater exposure to that dimension (112 vs. 40 trials). The physical distractor dimension carrying the spatial bias was counterbalanced across groups, so orientation and color had comparable exposure and spatial-regularity roles when considered across the two groups.

### Behavioral models and reliability

Statistical significance was evaluated using the multiplicity corrections specified for each analysis. Holm’s step-down procedure was used to control the family-wise error rate at α = .05 within the analysis-specific families defined in the Supplementary Methods and corresponding tables. Both unadjusted and Holm-adjusted p values are reported where applicable. Confidence intervals are pointwise 95% intervals unless otherwise stated.

### Participant-level attentional capture

RT analyses included correct trials with RTs between 200 and 3,000 ms. Condition-specific mean RTs, error rates and retained trial counts are reported in **Table 1**. Distractor capture was defined as the participant-level mean RT on distractor-present trials minus the mean RT on distractor-absent trials. Capture was calculated separately for the biased and unbiased distractor dimensions and across all distractor-present trials combined; the combined score weighted the two dimensions by each participant’s retained trial counts. Positive values indicate distractor-related slowing. Exact outcome definitions are given in **Supplementary Equation S1**.

**Table 1.**
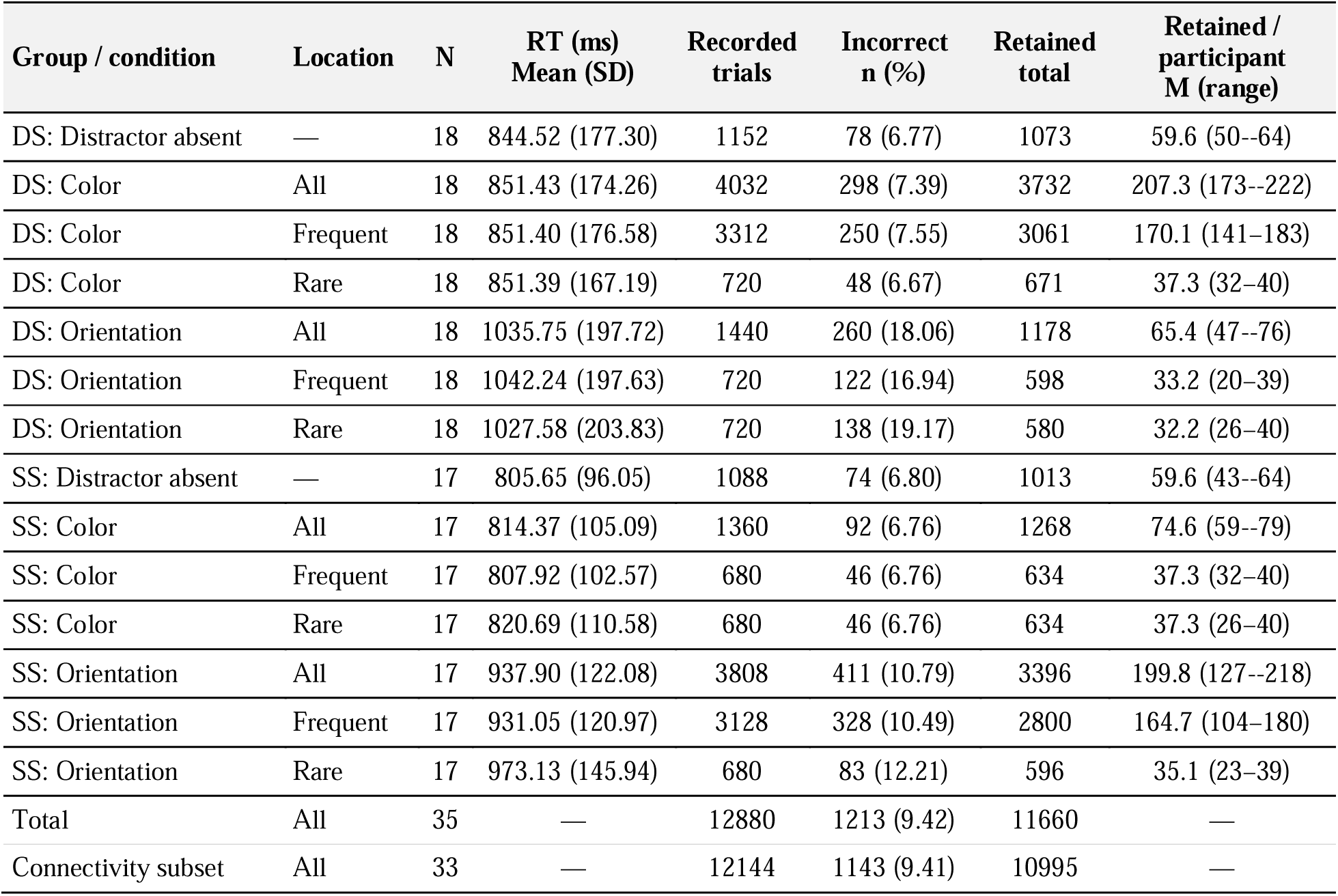
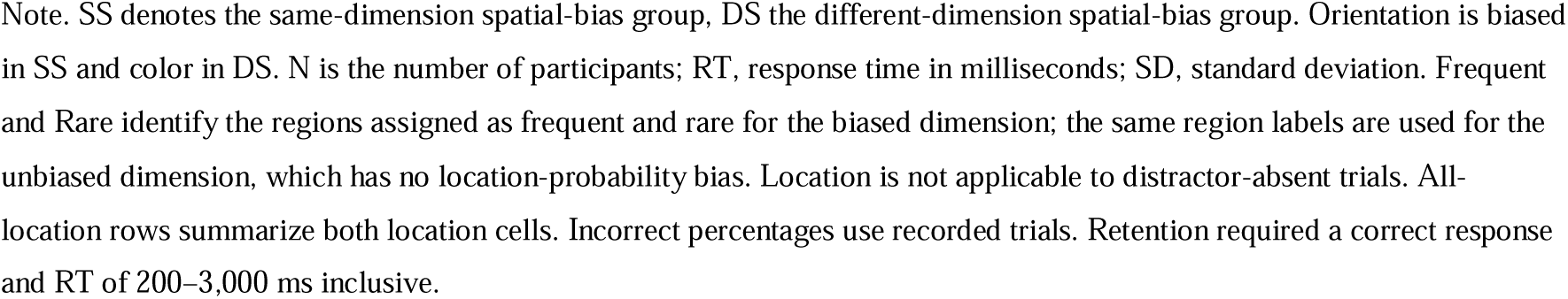
Response times, error rates and trial counts by distractor dimension and location.

| Group / condition | Location | N | RT (ms)<br>Mean (SD) | Recorded<br>trials | Incorrect<br>n (%) | Retained<br>total | Retained /<br>participant<br>M (range) |
| --- | --- | --- | --- | --- | --- | --- | --- |
| DS: Distractor absent | — | 18 | 844.52 (177.30) | 1152 | 78 (6.77) | 1073 | 59.6 (50--64) |
| DS: Color | All | 18 | 851.43 (174.26) | 4032 | 298 (7.39) | 3732 | 207.3 (173--222) |
| DS: Color | Frequent | 18 | 851.40 (176.58) | 3312 | 250 (7.55) | 3061 | 170.1 (141--183) |
| DS: Color | Rare | 18 | 851.39 (167.19) | 720 | 48 (6.67) | 671 | 37.3 (32--40) |
| DS: Orientation | All | 18 | 1035.75 (197.72) | 1440 | 260 (18.06) | 1178 | 65.4 (47--76) |
| DS: Orientation | Frequent | 18 | 1042.24 (197.63) | 720 | 122 (16.94) | 598 | 33.2 (20--39) |
| DS: Orientation | Rare | 18 | 1027.58 (203.83) | 720 | 138 (19.17) | 580 | 32.2 (26--40) |
| SS: Distractor absent | — | 17 | 805.65 (96.05) | 1088 | 74 (6.80) | 1013 | 59.6 (43--64) |
| SS: Color | All | 17 | 814.37 (105.09) | 1360 | 92 (6.76) | 1268 | 74.6 (59--79) |
| SS: Color | Frequent | 17 | 807.92 (102.57) | 680 | 46 (6.76) | 634 | 37.3 (32--40) |
| SS: Color | Rare | 17 | 820.69 (110.58) | 680 | 46 (6.76) | 634 | 37.3 (26--40) |
| SS: Orientation | All | 17 | 937.90 (122.08) | 3808 | 411 (10.79) | 3396 | 199.8 (127--218) |
| SS: Orientation | Frequent | 17 | 931.05 (120.97) | 3128 | 328 (10.49) | 2800 | 164.7 (104--180) |
| SS: Orientation | Rare | 17 | 973.13 (145.94) | 680 | 83 (12.21) | 596 | 35.1 (23--39) |
| Total | All | 35 | — | 12880 | 1213 (9.42) | 11660 | — |
| Connectivity subset | All | 33 | — | 12144 | 1143 (9.41) | 10995 | — |

Note. SS denotes the same-dimension spatial-bias group, DS the different-dimension spatial-bias group. Orientation is biased in SS and color in DS. N is the number of participants; RT, response time in milliseconds; SD, standard deviation. Frequent and Rare identify the regions assigned as frequent and rare for the biased dimension; the same region labels are used for the unbiased dimension, which has no location-probability bias. Location is not applicable to distractor-absent trials. All-location rows summarize both location cells. Incorrect percentages use recorded trials. Retention required a correct response and RT of 200–3,000 ms inclusive.

### Behavioral spatial learning

Spatial learning was defined as the rare-minus-frequent RT difference on biased-dimension trials (**see Supplementary Equation S1**). We tested whether RTs were slower for distractors appearing at rare than at frequent locations and whether this spatial-learning effect differed between the SS and DS groups. A Gaussian linear mixed-effects model (**Supplementary Equation S2**) was fitted to eligible biased-dimension trials across training and scanner sessions, with block as a continuous covariate, distractor location (frequent = 0, rare = 1), group (SS = 0, DS = 1), their interaction, and a participant-specific random intercept: RT ∼ Block + Location × Group + (1 | Subject). The model was estimated using restricted maximum likelihood (REML), with Satterthwaite approximations for degrees of freedom.

### Behavioral measurement reliability

We assessed the consistency of participant-level capture and spatial-learning scores using odd/even recorded trial indices and the two scanner runs. Odd/even correlations assessed split-half consistency within a session, whereas run-to-run correlations assessed stability across runs. Recorded parity was assigned before trial exclusions. Reliability was estimated in the pooled sample and separately within the SS and DS groups. Because uncentered pooled correlations can reflect both within-group individual differences and stable SS–DS mean differences, we also calculated group-centered pooled correlations. Group centering was performed separately within each half or run before pooling participants. Spearman–Brown-corrected coefficients were additionally reported for split-half analyses to estimate the reliability of the corresponding full-length score. Uncertainty was quantified using 10,000 paired participant-bootstrap resamples within the group [27], with group centering repeated in each resample. The exact score construction, centering, correlation, and Spearman–Brown equations are provided in **Supplementary Equation S3**.

### MRI acquisition and preprocessing

Functional images were acquired on a 3T Siemens Prisma scanner with a 32-channel head coil using a multiband echo-planar imaging sequence (TR = 1,000 ms, TE = 30 ms, flip angle = 45°, multiband factor = 4, 48 slices, 3 × 3 mm in-plane resolution 3-mm slice thickness with a 0.3-mm gap, field of view = 210 mm). Structural images were acquired with an MPRAGE sequence (TR = 2,500 ms, TE = 2.22 ms, TI = 1000 ms, flip angle = 8°, receiver bandwidth = 220 Hz/pixel, 0.8-mm isotropic voxels). We used fMRIPrep v23.1.4 [28] to preprocess the data, including motion correction, slice-timing correction, susceptibility-distortion correction, coregistration, normalization to MNI152NLin2009cAsym space, and resampling to the analysis space.

### Functional-connectivity pipeline

Connectivity analyses used the Schaefer-400 parcellation with Yeo-7 network assignments [29, 30]. For each run, the atlas was resampled to the corresponding fMRIPrep BOLD grid using nearest-neighbor interpolation and intersected with the run-specific brain mask. All connectivity, network, and edge-level analyses therefore used the same 399 anatomically covered parcels. Matrix columns were explicitly mapped to the corresponding native atlas labels. All 91 parcels assigned to the Yeo-7 Default network retained coverage in every run. The primary connectivity measure, termed Default91, was defined as the mean Fisher-z-transformed connectivity across the 4,095 unique within-network edges among these parcels.

Across the 66 task runs from 33 participants, each run contained 1,732 acquired volumes. After removal of non-steady-state volumes, the mean retained count was 1,730.73 per run (range 1,726–1,732). We extracted the mean preprocessed BOLD signal within each parcel at every volume. The same volume-selection mask was applied to parcel signals and nuisance regressors to preserve temporal alignment. The primary model included 24 motion regressors—the six rigid-body motion parameters, their temporal derivatives, their squared terms, and squared derivatives [31]—together with mean white-matter (WM) and cerebrospinal-fluid (CSF) signals. The global-signal, task-regression and combined variants included 27, 29 and 30 nuisance regressors, respectively; task regression added three Glover-HRF-convolved display-event regressors. Including the constant and linear trend removed during detrending, the corresponding design ranks were 28, 29, 31 and 32. Retained volumes minus these ranks yielded nominal residual dimensions averaging 1,702.73, 1,701.73, 1,699.73 and 1,698.73, respectively (ranges 1,698–1,704, 1,697–1,703, 1,695–1,701 and 1,694–1,700). Parcel signals and nuisance regressors were detrended and band-pass filtered between 0.01 and 0.10 Hz using TR = 1 s in Nilearn v0.13.1. After nuisance regression, residual parcel signals were standardized.

Within each run, Pearson correlations were calculated between all cleaned parcel time series and then Fisher-*z* transformed. Correlations were numerically bounded below ±1 where necessary to avoid infinite transformed values. The resulting connectivity matrices were averaged across runs with equal run weights, and matrix diagonals were set to zero.

### Primary DMN–behavior analyses

#### DMN connectivity and distractor capture

We correlated DMN connectivity with biased-dimension capture; unbiased and count-weighted combined capture were additional outcomes. Let denote participant (i)’s DMN mean Fisher-connectivity. For any participant-level quantity, pooled analyses removed the corresponding SS or DS group mean (group centering), 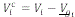 = *V_i_* – *V_gi_*, where *V_g_* is the mean for the SS or the DS group. This centering expression was applied separately to connectivity and capture. The primary pooled brain–behavior association was *r_FB_* = *cor*(*F^c^*, *B^c^*) where *B^c^*denotes group-centered biased-dimension capture. A positive value indicates that participants with higher DMN connectivity relative to their group mean tend to show greater biased-dimension capture relative to their group mean. We additionally calculated the corresponding Pearson correlations separately within the SS and DS groups. The within-group analyses were performed on the original scores. All correlations were evaluated using two-sided tests. The full notation is given in **Supplementary Equation S4.**

To describe the contributions of the two groups to the pooled biased-capture association, we partitioned the group-centered connectivity–capture cross-product into SS and DS components. Both components were normalized by the same pooled correlation denominator, so their contributions summed to the pooled correlation. Covariance shares expressed each component relative to the total cross-product. Pointwise 95% percentile intervals used 10,000 paired participant-bootstrap samples stratified by group, with centering repeated in each sample (Supplementary Equation S4).

We directly compared the DMN correlations with biased and unbiased capture using Williams’ test [32] for two dependent correlations that share the same connectivity variable. The test additionally accounts for the correlation between the two capture scores; its full specification is given in **Supplementary Equation S5**.

Partial correlations were used to assess whether the associations between DMN connectivity and behavior remained after adjusting for specified covariates. Connectivity and behavioral scores were separately regressed on the same covariates, and the resulting residuals were correlated using Pearson’s r (**Supplementary Equation S6**). For biased-dimension capture, pooled analyses controlled separately for (1) unbiased-dimension capture and group and (2) inter-trial priming and group. Inter-trial priming was defined as the mean RT on current biased-dimension distractor trials following a biased-dimension distractor minus the mean RT following either an unbiased-dimension distractor or a distractor-absent display. Within-group analyses controlled for unbiased-dimension capture only. All tests were two-sided.

### DMN connectivity and spatial location learning

The same participant-level procedure was applied to spatial learning, replacing capture with each participant’s rare-minus-frequent RT difference for the biased dimension. We report the pooled group-centered correlation and the corresponding SS and DS correlations.

### Robustness and sensitivity analyses

We assessed the robustness of the DMN connectivity associations with attentional capture and spatial learning using complementary participant-resampling, influence, preprocessing, trial-count matching, and trial-history analyses.

#### Participant-level robustness

Pointwise 95% percentile confidence intervals were estimated from 10,000 participant-bootstrap resamples stratified by group, with group means recalculated in each resample (Supplementary Equation S7). Within-group permutation tests (10,000 permutations) provided two-sided significance estimates (Equation S8). Leave-one-participant-out analyses assessed the influence of individual observations, with group centering repeated after each omission (Equation S9).

#### Connectivity-preprocessing sensitivity

Connectivity-preprocessing sensitivities retained the same atlas, filtering, and run aggregation while varying nuisance regression. These analyses included omission of nuisance regression, addition of global signal, addition of three Glover-HRF-convolved regressors for recorded display events, and addition of both task and global-signal regressors (Equation S10; Table S4A-B). To quantify nuisance-related variance, we regressed detrended, band-pass-filtered parcel time series on identically processed nuisance regressors, calculated total explained variance (R²) and the reduction in R² after removing each regressor set and refitting the model, and averaged these estimates across parcels within runs and then across all 66 runs. In DMN parcels, the primary (26-regressor) and full (30-regressor) models explained 45.48% and 56.94% of filtered time-series variance, respectively, with unique contributions of 13.71 percentage points for motion, 3.26 for WM/CSF, 10.03 for global signal and 1.62 for display-event regressors (Table S4C).

#### Conditional trial-count matching

To assess spatial-sampling imbalance, we assessed the influence of unequal trial counts by randomly subsampling biased-dimension trials to match the number of eligible unbiased-dimension trials (Equation S11). We also recalculated capture scores with equal weight assigned to frequent– and rare-region RTs and repeated the connectivity associations and biased-versus-unbiased correlation comparison (Equation S11). For spatial learning, we matched frequent– and rare-region trial counts separately within each participant and scanner run, repeating the subsampling 10,000 times and recalculating the connectivity–learning association. Because matching could change the relative contributions of the two runs, we additionally calculated an expected matched learning score from the original run-specific RT differences, weighted by the matched trial counts. This allowed us to distinguish changes due to run weighting from variability caused by random trial selection. These analyses assessed the sensitivity of the associations to unequal trial counts and spatial sampling but did not eliminate differences in measurement precision or prior exposure.

#### Trial-history adjustment

Hierarchical models assessed whether connectivity associations persisted after accounting for scanner run, chronological trial order, and prior biased– and unbiased-dimension exposures derived from the complete recorded trial sequence (Equation S12). The capture model included 10,995 eligible trials across three distractor conditions, whereas the learning model included 6,721 biased-dimension trials and retained location × connectivity × group and location × connectivity × run interactions. Both models preserved participant-specific random effects and used REML estimation with Satterthwaite inference. Model convergence and optimizer agreement were evaluated.

### Hierarchical trial-level connectivity models

#### Hierarchical trial-level capture model

Gaussian mixed-effects models assessed connectivity–RT coupling at the trial level, accounting for repeated observations within participants and unequal trial information. RT was divided by 100, DMN connectivity centered within SS and DS and scaled by the pooled participant SD, and Group coded SS = 0 and DS = 1. We fitted the models with lme4/lmerTest using REML estimation and Satterthwaite inference, and reported coupling in milliseconds per connectivity SD. The capture model retained all 10,995 eligible correct-RT scanner trials from 33 participants (SS = 16, DS = 17). Condition distinguished Absent, BiasedFrequent, BiasedRare, UnbiasedFrequent and UnbiasedRare trials, with Absent, SS and Run 1 as reference levels. On unbiased trials, frequent and rare denote the assigned spatial regions, not a spatial-probability contingency in that dimension. The model included condition-specific connectivity effects and group interactions, condition-specific run shifts, and correlated participant-specific random intercepts and slopes for the four distractor cells:

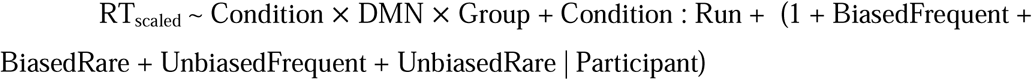

Condition × DMN interactions estimate condition-specific present-minus-absent RT– connectivity slopes. A positive slope indicates greater distractor-related slowing at higher connectivity.

We derived equal-region biased and unbiased coupling by averaging each dimension’s frequent– and rare-region slopes with equal weights. Pooled contrasts weighted SS and DS by 16/33 and 17/33, with uncertainty calculated from the fitted coefficient covariance.

To distinguish physical distractor dimension from assigned spatial bias, we reconstructed orientation– and color-specific participant capture scores and assessed their connectivity associations separately. We also compared SS and DS within each physical dimension using contrasts of the same hierarchical capture model, rather than separate subgroup fits. The full model and contrast definitions are given in **Supplementary Equation S13**.

### Hierarchical spatial-learning model

The spatial-learning model used 6,721 eligible biased-dimension scanner trials from the same 33 participants to assess DMN coupling with the rare-minus-frequent RT contrast. ‘Location’ is coded frequent = 0 and rare = 1. The binary run indicator, shown as Run in the formula, coded run-1 as 0 and run-2 as 1. The model included correlated participant-specific random intercepts and location slopes:

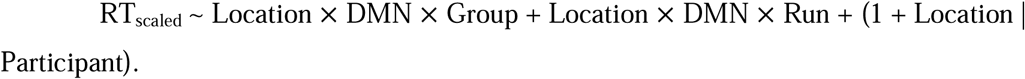

Location × DMN estimates SS DMN–learning coupling in Run 1, and Location × DMN × Group estimates its DS-minus-SS difference. Location × DMN × Run2 tests the run-2-minus-run-1 change in coupling, assumed common across SS and DS. We averaged run-specific coupling slopes with equal run weights and pooled SS and DS using weights of 16/33 and 17/33. The complete model specification and coupling contrasts are given in **Supplementary Equation S14**.

### Network-summary associations

Mean Fisher-z connectivity within and between the seven Yeo networks yielded 28 summaries, averaged across constituent edges and likewise across runs. Pooled associations centered connectivity and behavior within SS and DS. Holm-28 corrections were applied within each network-summary screen. For each outcome, six approximate Williams tests directly compared the pooled within-DMN association with the other within-network associations (Holm-6). These tests compared association strengths, rather than inferring specificity from separate significance levels.

A following sensitivity analysis assessed all 168 network-summary associations (28 network pairs × two outcomes × three samples) using 10,000 synchronized within-SS/DS permutations. We report unadjusted permutation p values, permutation p values adjusted by Holm-28, and permutation maxT-28 p values separately. Both corrections use the same 28-test outcome/sample panel. Holm-28 applies Holm’s sequential Bonferroni procedure to the 28 individual permutation p values, controlling the family-wise error rate under arbitrary dependence; maxT-28 instead compares each observed association statistic with the permutation distribution of the maximum absolute association statistic across the 28 tests; synchronized permutations retain dependence among the tests. maxT probabilities are already multiplicity adjusted and were not additionally Holm adjusted. Consequently, the two procedures can yield different adjusted p values and significance decisions (**Table 5; Table S8**). A matched exploratory extension assessed equal-region biased capture and spatial learning against the seven within-network summaries in pooled, SS and DS samples. Capture used equal frequent/rare region weights before subtracting the absent baseline, rather than the original observed mixture. Full procedures are given in **Supplementary Methods S15**.

### Signed-edge screens and enrichment

The full connectome comprised all unique, undirected connections between the 399 parcels retained from the Schaefer-400 atlas after applying the common coverage criterion. Self-connections and duplicate connections were excluded, yielding 399 × 398 / 2 = 79,401 possible edges. This complete set served as the reference universe for evaluating whether connectivity–behavior associations were disproportionately concentrated within particular networks. For each edge, we calculated the pooled, group-centered Pearson correlation between connectivity and behavioral performance, separately for distractor capture and spatial learning. Edges meeting an exploratory, uncorrected two-sided threshold of *p* < .01 were classified as positive or negative according to the direction of their association. Selected edges were counted within 28 mutually exclusive within– and between-network categories and seven overlapping network-endpoint categories. For each category, the expected number of selected edges was calculated as the total number of selected edges of the corresponding sign multiplied by that category’s proportion of all 79,401 possible edges. Observed-to-expected ratios therefore quantified whether selected associations were more or less concentrated in a network category than expected from its number of possible connections.

These ratios were descriptive; formal enrichment was assessed using 10,000 within-group behavioral-label permutations that recomputed the signed edge screens and category-surplus statistics, with Holm correction across the specified network-category families (**Supplementary Methods S15**). The edge-enrichment tables report an exploratory maxT-type adjustment implemented as maxZ, alongside unadjusted permutation p and Holm-adjusted permutation p values (**Table 6; Table S9-S10**). Pair-category families include both signs (56 tests), and network-touching families include both signs (14 tests), separately for each outcome.

### Exploratory prediction using DMN subnetwork connectivity

Connectome-based prediction evaluated distractor capture and spatial learning separately using six predefined within– and between-network DefaultA/B/C edge sets from the Yeo-17 atlas. These subdivisions complemented, rather than partitioned, the focal Yeo-7 Default91 measure. In each leave-one-participant-out fold, edges associated with the behavioral outcome (two-sided *p* < .01) were selected using only the training participants. For each participant, signed network strength was calculated as the sum of Fisher-z connectivity across positively associated edges minus the sum across negatively associated edges. An ordinary least-squares model fitted to the training participants predicted group-centered behavioral scores from network strength, and the fitted coefficients were applied to the held-out participant. Prediction performance was quantified as the Pearson correlation between observed and predicted held-out scores across participants. Significance was assessed using 10,000 within-group permutations that repeated the complete prediction procedure. Detailed equations, correction procedures, and results are provided in **Supplementary Methods S16** and the **Supplementary Results**.

### Software

MRI preprocessing was performed with fMRIPrep v23.1.4. Functional-connectivity re-extraction and processing-sensitivity analyses used Python 3.13.11 with Nilearn 0.13.1 and NiBabel 5.3.3 for image handling, parcel time-series extraction, nuisance regression, filtering, and connectivity estimation. Behavioral summaries, participant-level correlations and direct comparisons, reliability estimates, trial-count matching, bootstrap and permutation analyses, and signed-edge enrichment analyses were implemented using custom Python scripts with NumPy 2.4.2, pandas 3.0.0, SciPy 1.17.0, and statsmodels 0.14.6. Behavioral and hierarchical brain–behavior mixed-effects models, including capture, spatial learning, and trial-history sensitivity analyses, were fitted in R 4.5.2 using lme4 1.1-38 and lmerTest 3.2-0. Nested cross-validated connectome-based prediction used custom NumPy/SciPy implementations for edge selection, model fitting, and permutation inference. Figures were generated using Matplotlib 3.10.8.

## Results

### Behavioral outcomes and measurement properties

#### Spatial learning

Behavioral analyses included all 35 scanner participants (SS = 17, DS = 18). Orientation was the biased dimension in SS and the unbiased dimension in DS; color had complementary roles. The joint training-plus-scanner learning model on the biased-dimension trials (*RT ∼ Block + Location × Group + (1 | Subject)*) estimated a larger rare-minus-frequent RT effect in SS than DS (Figure 2A; DS minus SS = −35.63 ms, 95% CI [−56.42, −14.84], *t*(11402.08) = −3.36, raw *p* < .001). The mean rare-minus-frequent RT difference was positive in SS (44.93 ms, 95% CI [14.47, 75.38], *t*(16) = 3.13, raw *p* = .0065, Holm-2 *p* = .0130,= *d_z_*.76, 95% CI [.38, 1.35]) and uncertain in DS (10.27 ms, 95% CI [−6.56, 27.11], *t*(17) = 1.29, raw/Holm-2 *p* = .2153,*d_z_*= .30, 95% CI [−.16, .79]).

**Figure 2.**
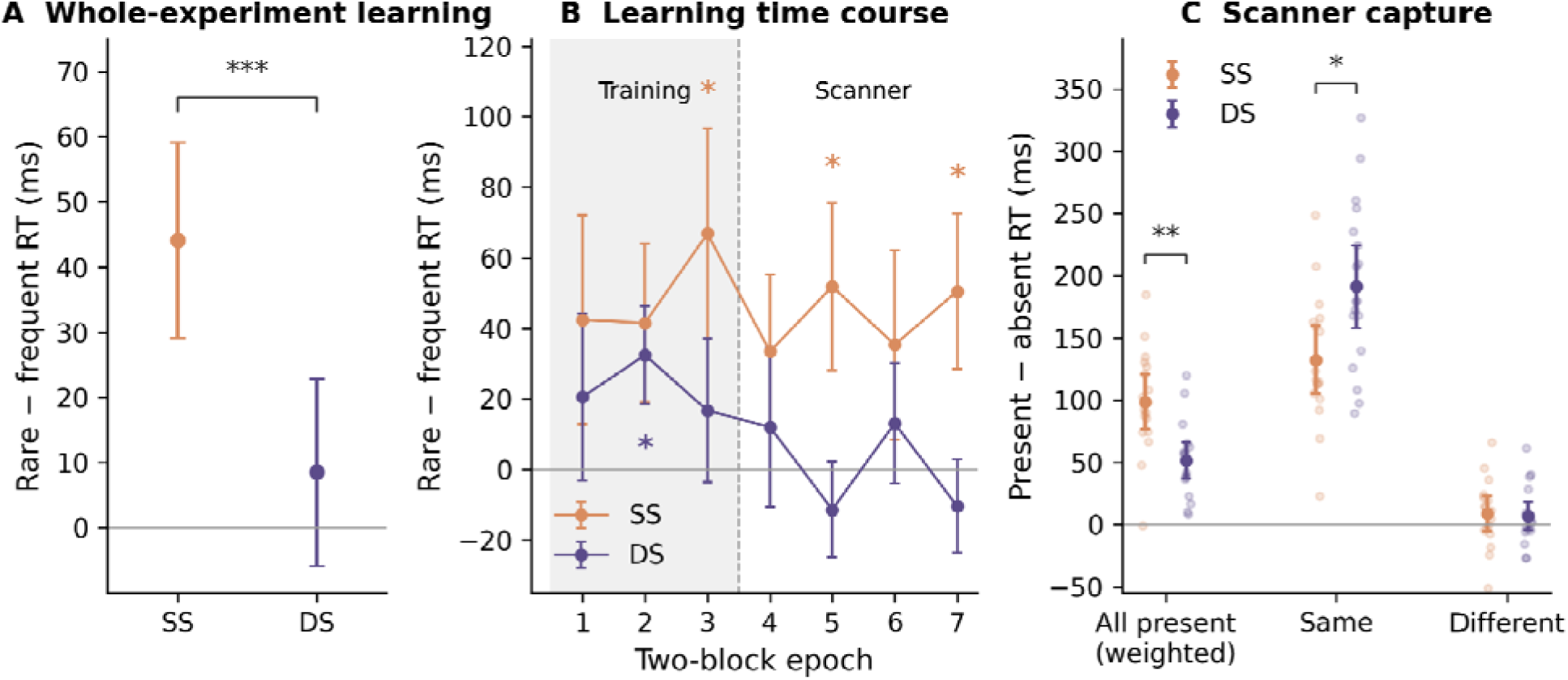
Behavioral effects. (**A**) Mixed-model estimated rare-minus-frequent RT differences across training and scanner phases (behavioral mixed model described in Methods). (**B**) Rare-minus-frequent RT differences by two-block epoch; shading denotes training, and the dashed line separates training from scanning. (**C**) Scanner capture, defined as distractor-present minus distractor-absent RT, for all distractor-present trials (weighted by each participant’s retained condition counts), same-dimension orientation distractors and different-dimension color distractors. SS denotes the orientation-biased group and DS the color-biased group. Bars show pointwise 95% confidence intervals in A/C and SEM in B; pale points in C are individual participants. The y axes in A/B show (distractor) location differences, whereas C shows capture costs. Stars denote * raw p < .05, ** p < .01, and *** p < .001.

The time course comprised seven consecutive two-block epochs: epochs 1–3 covered training and epochs 4–7 scanning (Figure 2B). SS showed nominally positive rare-minus-frequent learning in epoch 3 (66.88 ms, raw *p* = .0413, Holm-7 *p* = .2543, *d_z_* = 0.58), epoch 5 (51.88 ms, raw *p* = .0440, Holm-7 *p* = .2543, *d_z_* = 0.53), and epoch 7 (50.53 ms, raw *p* = .0363, Holm-7 *p* = .2543, *d_z_* = 0.55). DS showed a nominally positive effect in epoch 2 (32.43 ms, raw *p* = .0352, Holm-7 *p* = .2463, *d_z_*= 0.60). No individual epoch survived Holm correction across the seven epochs within its group. The remaining epochs were uncertain even before correction.

#### Distractor capture

Trial-weighted capture was positive in both groups (SS: 98.70 ms; DS: 51.35 ms; both raw and Holm-5 *p* < .001; SS *d_z_* = 2.32, DS *d_z_* = 1.72). It was smaller in DS than SS (DS minus SS = −47.35 ms, 95% CI [−72.90, −21.80], raw *p* = .0007, Holm-5 *p* = .0021, *g* = −1.27). Orientation capture was positive in both groups (SS: 132.25 ms; DS: 191.23 ms; both raw and Holm-5 *p* < .001; SS *d_z_* = 2.49, DS *d_z_* = 2.86). Color-capture estimates were uncertain in SS (8.73 ms, raw *p* = .2092, Holm-5 *p* = .4184, *d_z_* = 0.32) and DS (6.91 ms, raw *p* = .2152, Holm-5 *p* = .4304, *d_z_*= 0.30). DS showed greater orientation capture than SS (DS minus SS = 58.98 ms, 95% CI [17.52, 100.44], raw *p* = .0067, Holm-5 *p* = .0134, *g* = 0.95). The color-capture difference was uncertain (−1.81 ms, 95% CI [−19.27, 15.64], raw/Holm-5 *p* = .8335, *g* = −0.07). Full results are provided in Supplementary Table S1.

### Reliability of capture and spatial-learning scores

Reliability estimates are summarized in **Table 2**. SS orientation/biased capture showed positive odd–even consistency and run-to-run stability after correction; DS color/biased capture remained uncertain. Group-adjusted pooled capture showed positive run-to-run stability, whereas odd–even consistency was uncertain. Spatial-learning scores showed positive consistency and stability in the pooled sample and SS, with uncertain estimates in DS. The DMN connectivity predictor showed positive run-to-run stability both after group centering and without centering (**Table 2C**).

**Table 2.**
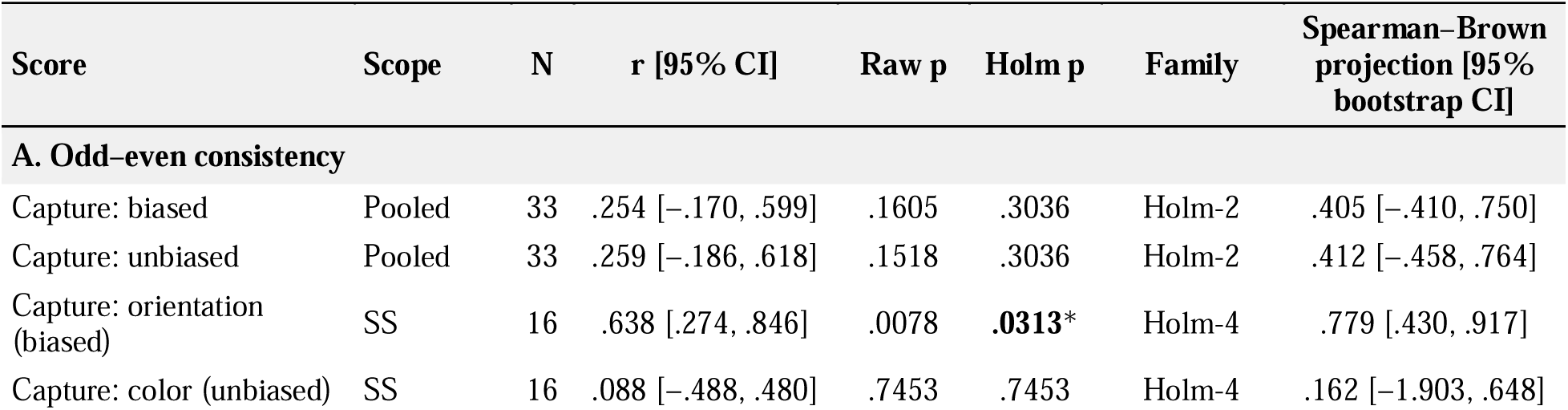
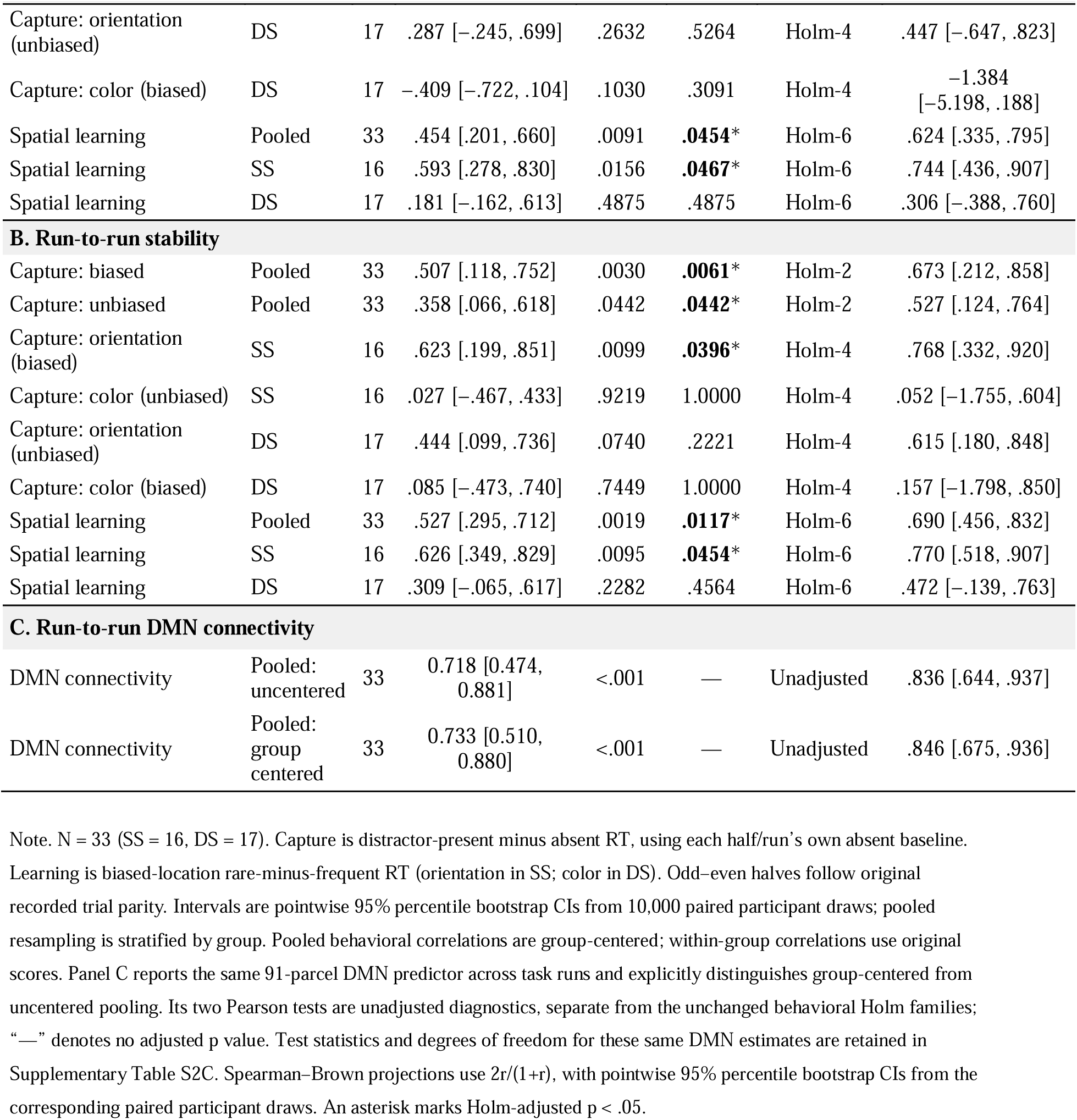
Reliability of capture and spatial-learning scores and run-to-run DMN connectivity.

Full-sample checks (N = 35) showing consistent results are summarized in **supplementary Table S2**. Uncentered pooled capture correlations were higher in the imaging sample than the corresponding group-centered estimates: odd–even r = .699/.776 and run-to-run r = .807/.819 for biased/unbiased capture, respectively (Supplementary Table S2B,D). SS–DS mean separation contributed 85.5% and 90.2% of the biased/unbiased odd– even covariance. This measures consistency of the connectivity predictor across the two task runs, not absolute stable trait reliability.

## Neural–behavioral association

### DMN connectivity and distractor capture

Higher within-DMN task-run connectivity was associated with larger biased-dimension capture in the pooled group-centered analysis (r = .550, 95% Fisher-z CI [.250, .754], focal raw p = .0011; Figure 3; Table 3A). The corresponding unbiased-dimension association was uncertain (r = .008, 95% Fisher-z CI [−.341, .356], raw/Holm-8 p = .9633). SS participants, for whom orientation was the biased dimension, contributed 80.1% of the pooled group-centered biased-capture covariance (95% bootstrap CI [52.9%, 96.3%]); DS participants, for whom color was biased, contributed 19.9% [3.7%, 47.1%]. Their respective contributions to the pooled correlation were .441 [.224, .601] and .109 [.020, .246]. The direct comparison supported a stronger DMN association with biased than unbiased capture (Δr = .542, 95% participant-bootstrap CI [.249, .842], raw p = .0052, Holm-2 p = .0100).

**Figure 3.**
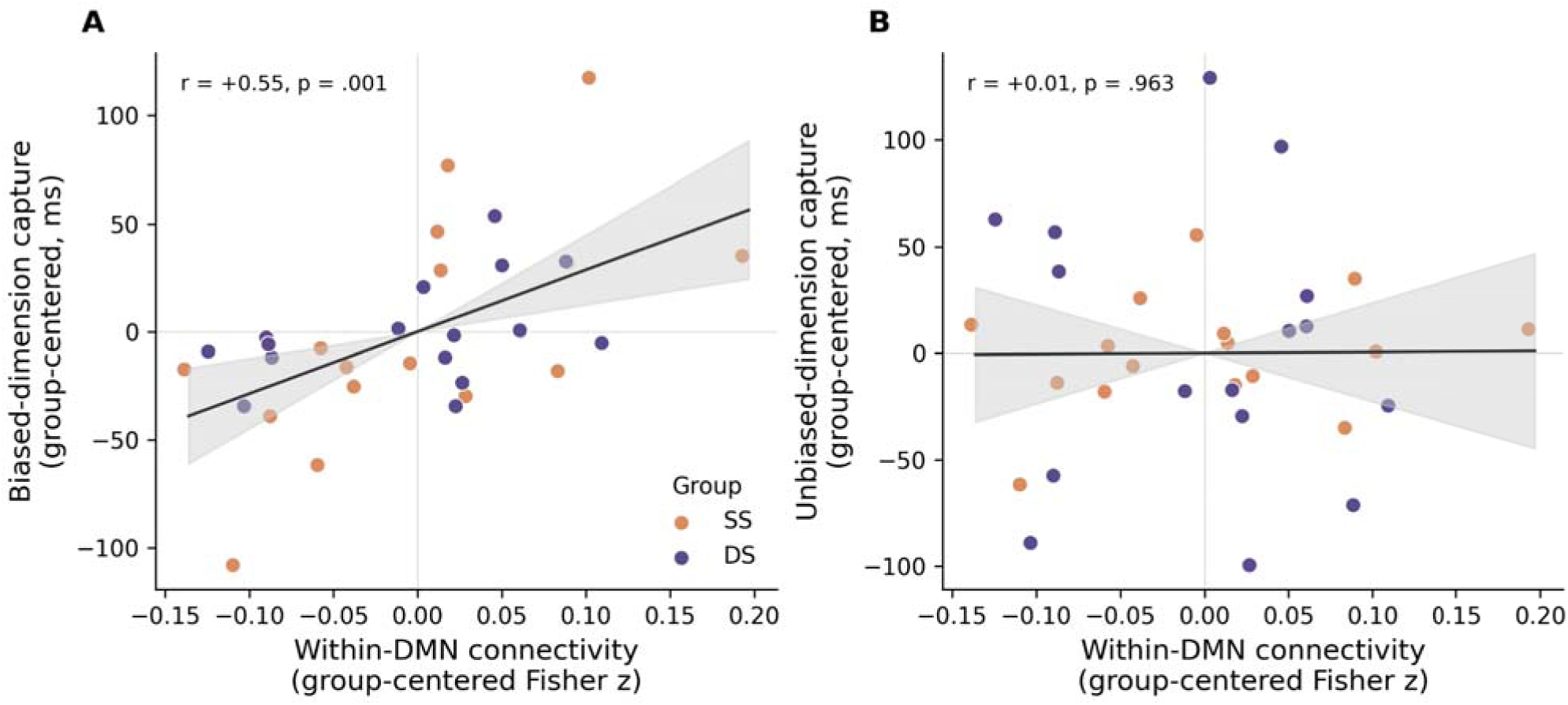
Associations of the anatomically defined 91-parcel Yeo-7 Default network with capture by the biased. (**A**) and unbiased (**B**) distractor dimension. Connectivity and RT contrasts were centered separately within SS and DS before pooling; each point is one participant (SS n=16, DS n=17). Lines show the pooled group-adjusted fits and shading their pointwise 95% confidence bands conditional on centering. Raw two-sided correlation p values are displayed. SS, same-dimension spatial-bias group; DS, different-dimension spatial-bias group.

**Table 3.**
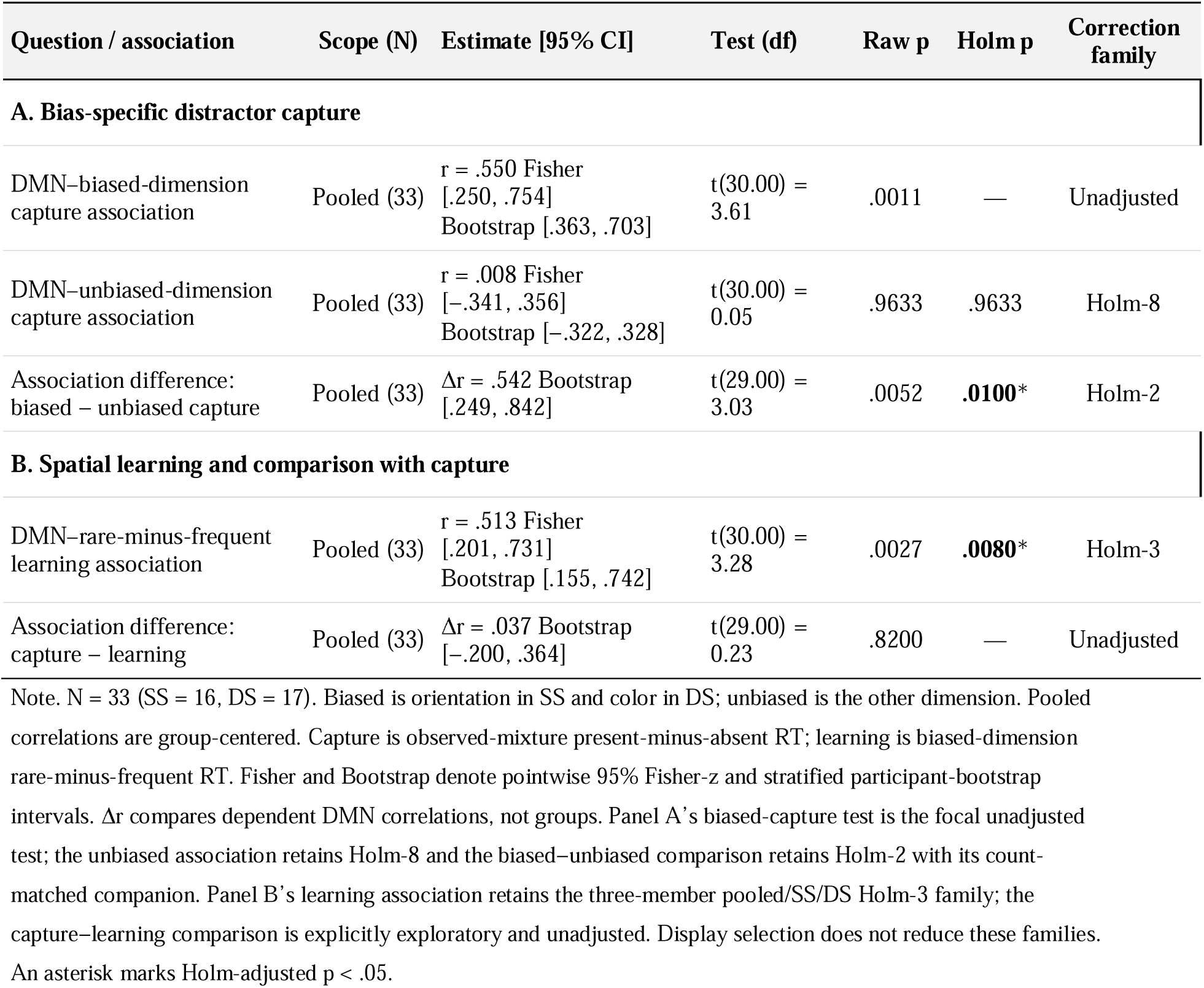
Principal participant-level DMN associations and direct comparisons.

| Question / association | Scope (N) | Estimate [95% CI] | Test (df) | Raw p | Holm p | Correction family |
| --- | --- | --- | --- | --- | --- | --- |
| **A. Bias-specific distractor capture** |  |  |  |  |  |  |
| DMN–biased-dimension capture association | Pooled (33) | $r = .550$ Fisher<br>[.250, .754]<br>Bootstrap [.363, .703] | $t(30.00) = 3.61$ | .0011 | — | Unadjusted |
| DMN–unbiased-dimension capture association | Pooled (33) | $r = .008$ Fisher<br>[-.341, .356]<br>Bootstrap [-.322, .328] | $t(30.00) = 0.05$ | .9633 | .9633 | Holm-8 |
| Association difference: biased – unbiased capture | Pooled (33) | $\Delta r = .542$ Bootstrap<br>[.249, .842] | $t(29.00) = 3.03$ | .0052 | **.0100\*** | Holm-2 |
| **B. Spatial learning and comparison with capture** |  |  |  |  |  |  |
| DMN–rare-minus-frequent learning association | Pooled (33) | $r = .513$ Fisher<br>[.201, .731]<br>Bootstrap [.155, .742] | $t(30.00) = 3.28$ | .0027 | **.0080\*** | Holm-3 |
| Association difference: capture – learning | Pooled (33) | $\Delta r = .037$ Bootstrap<br>[-.200, .364] | $t(29.00) = 0.23$ | .8200 | — | Unadjusted |

Note. $N = 33$ (SS = 16, DS = 17). Biased is orientation in SS and color in DS; unbiased is the other dimension. Pooled correlations are group-centered. Capture is observed-mixture present-minus-absent RT; learning is biased-dimension rare-minus-frequent RT. Fisher and Bootstrap denote pointwise 95% Fisher-z and stratified participant-bootstrap intervals. $\Delta r$ compares dependent DMN correlations, not groups. Panel A’s biased-capture test is the focal unadjusted test; the unbiased association retains Holm-8 and the biased–unbiased comparison retains Holm-2 with its count-matched companion. Panel B’s learning association retains the three-member pooled/SS/DS Holm-3 family; the capture–learning comparison is explicitly exploratory and unadjusted. Display selection does not reduce these families. An asterisk marks Holm-adjusted $p < .05$ .

#### Location-specific capture

In the pooled group-centered analysis, within-DMN connectivity was positively associated with biased-dimension capture at both rare and frequent locations. The rare-location association survived Holm-9 correction (r = .634, Holm-9 p = .0008), whereas the frequent-location association did not (r = .459, raw p = .0083, Holm-9 p = .0579). A direct comparison did not establish a difference between the correlations (rare minus frequent: Δr = .175, raw p = .1672, Holm-9 p = .3344).

#### Partial-correlation

The pooled biased-capture association persisted in separate partial-correlation analyses adjusting for unbiased capture and group (partial r = .577, raw p = .0007, Holm-8 p = .0054) and, respectively, the prior-display priming contrast and group (partial r = .559, raw p = .0011, Holm-8 p = .0076; Table S3A). Neither physical-orientation nor physical-color capture showed a corrected pooled association (Table S3B).

#### Participant-level robustness

This focal association had a positive stratified participant-bootstrap 95% CI [.363, .703] and was supported by within-group permutation (raw p = .0016). All 33 leave-one-participant-out estimates remained positive (rs = .504 – .579, ps = .0006 – .0038), indicating that no single participant accounted for the association.

#### Connectivity-preprocessing sensitivity

Preprocessing sensitivities retained the capture association when nuisance regression was omitted (r = .55, raw p = .0013, Holm-5 p = .0044) or task regressors were added (r = .563, raw p = .0008, Holm-5 p = .0040). Global-signal regression attenuated it, with or without task regression (*r*s = .38–.39, raw *p*s = .0271–.0343, both Holm-5 p = .0543; **Table S4A**).

#### Conditional trial-count matching

After conditional trial-count matching, the median biased-capture association was .50 (95% conditional interval [.36, .62]). The comparison using participant-averaged count-matched scores retained the biased-minus-unbiased association difference (Δr = .544, raw p = .0050, Holm-2 p = .0100; **Table S3A**). Equal-region standardization also retained the pooled biased-capture association (r = .629, 95% participant-bootstrap CI [.430, .769], raw p = .0001, Holm-6 p = .0007) and its difference from unbiased capture (Δr = .623, raw p = .0009, Holm-3 p = .0028; **Table S3C**).

#### Trial-history adjustment

In the separate trial-history capture model, adjustment for participant-specific run effects, trial order and accumulated prior biased/unbiased exposure retained positive pooled biased-capture coupling (20.27 ms/DMN SD, 95% CI [7.98, 32.56], raw p = .0022, Holm-9 p = .0173). Its biased-minus-unbiased slope difference did not survive correction (21.75 ms/DMN SD, raw p = .0253, Holm-9 p = .1770; **Table S5A-B**).

### DMN connectivity and spatial location learning

DMN connectivity was also associated with the scanner rare-minus-frequent RT contrast on biased-dimension trials (pooled r = .513, 95% Fisher-z CI [.201, .731], raw p = .0027, Holm-3 p = .0080; **Table 3B**). This positive association indicates a higher DMN connectivity and a larger RT difference between rare and frequent distractor locations. The spatial-learning association survived its Holm-3 correction in SS, but remained uncertain in DS (**Supplementary Table S3D**).

#### Overlap between spatial learning and overall capture

The rare-minus-frequent RT difference was positively correlated with equal-region biased-dimension capture, defined as the equally weighted mean of frequent-and rare-location capture (r = .691, Holm-9 p = .0001). After adjusting for this equal-region capture score, the pooled DMN–spatial-learning association was attenuated from r = .513 to partial r = .140 (unadjusted p = .4521). Adjustment for the original observed-mixture biased-dimension capture score also yielded no association surviving Holm-9 correction (partial r = .384, raw p = .0330, Holm-9 p = .0991). All analyses adjusted for SS/DS group membership. These analyses did not clearly establish an association between DMN connectivity and spatial modulation beyond its association with overall distractor capture.

#### Participant-level robustness

The pooled learning association had a 95% participant-bootstrap interval of [.155, .742]; corresponding SS and DS intervals were [.119, .867] and [−.233, .706]. Within-group permutation supported the pooled association (raw p = .0040, Holm-3 p = .0120) and the SS association (raw p = .0082, Holm-3 p = .0164), whereas DS remained uncertain (raw p/Holm-3 p = .2499). All 33 leave-one-participant-out estimates remained positive (*r*s = .412–.579).

#### Connectivity-preprocessing sensitivity

The primary association retained support in the new five-variant learning family (r = .513, raw p = .0027, Holm-5 p = .0133), as did added task regression (r = .504, raw p = .0033, Holm-5 p = .0133). Associations were smaller with nuisance regression omitted (r = .358, raw p = .0445, Holm-5 p = .0828), global-signal regression added (r = .389, raw p = .0279, Holm-5 p = .0828), and combined task/global-signal regression (r = .389, raw p = .0276, Holm-5 p = .0828). Thus, corrected support depended on preprocessing; these descriptive differences do not establish a significant difference between correlations or mediation by the removed signals. Numerical results are in **Supplementary Table S4B**.

#### Conditional trial-count matching

Matching frequent and rare trial counts within participants and runs yielded a median correlation of .465 (mean .462; central 95% conditional range [.321, .587]). The full-cell, matched-run-weight reference was r = .513. The smaller average correlation across random subsets is consistent with induced score variability and the nonlinearity of correlation.

#### Trial-history adjustment

In the trial-history learning model, the equal-run-averaged pooled coupling remained positive (26.91 ms/DMN SD, pointwise 95% CI [7.73, 46.09], raw p = .0076, Holm-4 p = .0228). Adjusted coupling was positive in SS (40.63 ms/DMN SD, CI [15.51, 65.75], raw p = .0025, Holm-4 p = .0099), but uncertain in DS (14.00 ms/DMN SD, CI [−14.77, 42.76], raw p = .3276, Holm-4 p = .3288). The direct DS-minus-SS contrast also remained uncertain (−26.64 ms/DMN SD, CI [−64.82, 11.55], raw p = .1644, Holm-4 p = .3288). The fixed design was full rank, and convergence, nonsingularity, and alternate-optimizer checks passed. Model coefficients and coupling contrasts are in **Supplementary Table S5C–D**.

## Hierarchical trial-level models: DMN coupling

### Hierarchical trial-level capture model

Higher DMN connectivity was associated with larger biased-dimension capture costs (pooled coupling = 29.00 ms/DMN SD, 95% CI [14.93, 43.07], raw p = .0002, Holm-9 p = .0019). A 1-SD increase in group-centered connectivity corresponded to an estimated 29-ms increase in distractor-related slowing relative to absent trials. Unbiased-dimension coupling remained uncertain (−0.53 ms/DMN SD, 95% CI [−19.06, 17.99], raw p = .9533, Holm-9 p = 1.0000). The direct biased-minus-unbiased coupling difference was positive (29.54 ms/DMN SD, 95% CI [10.26, 48.82], raw p = .0039, Holm-9 p = .0234; **Table 4A**), supporting stronger coupling with capture in the assigned biased dimension.

**Table 4.**
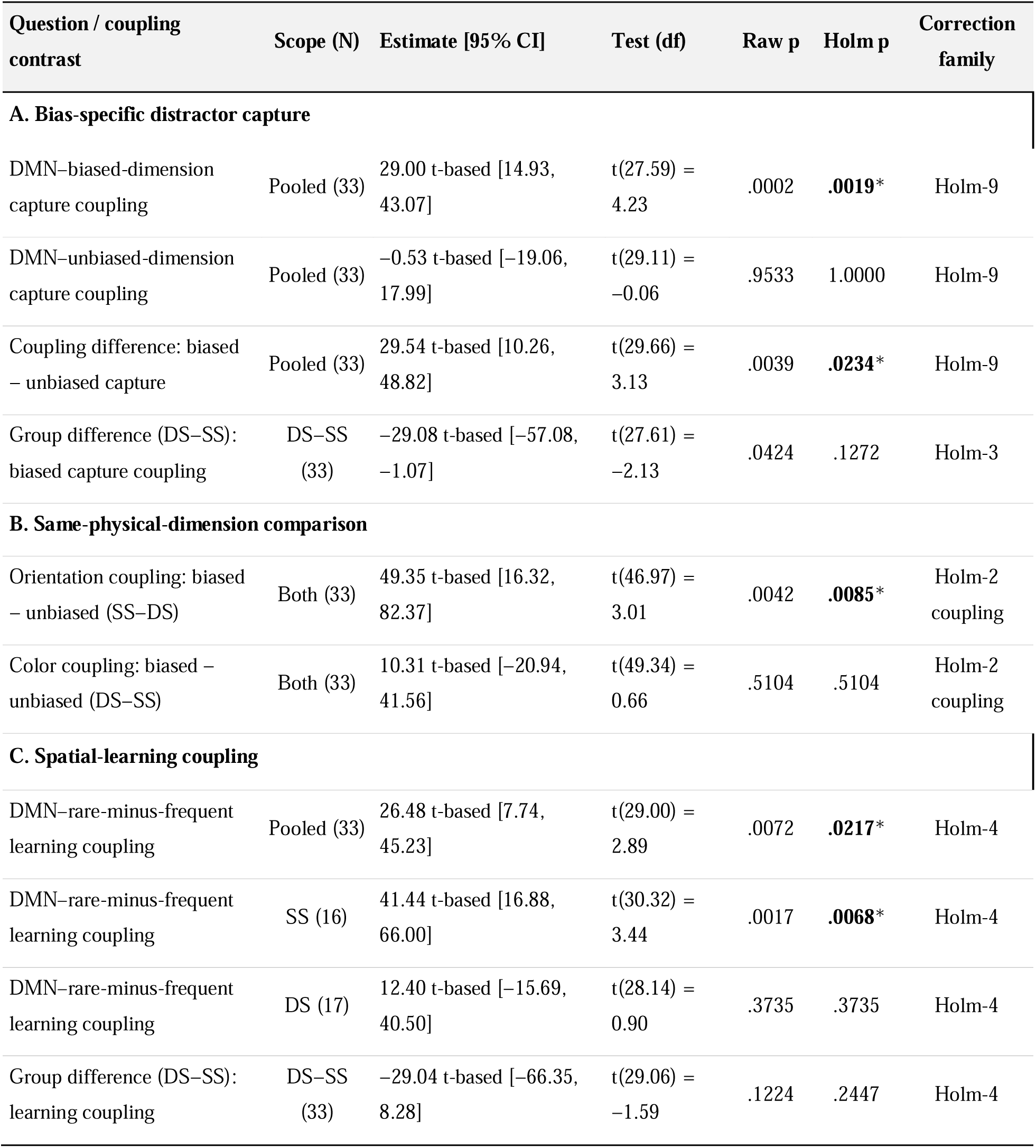
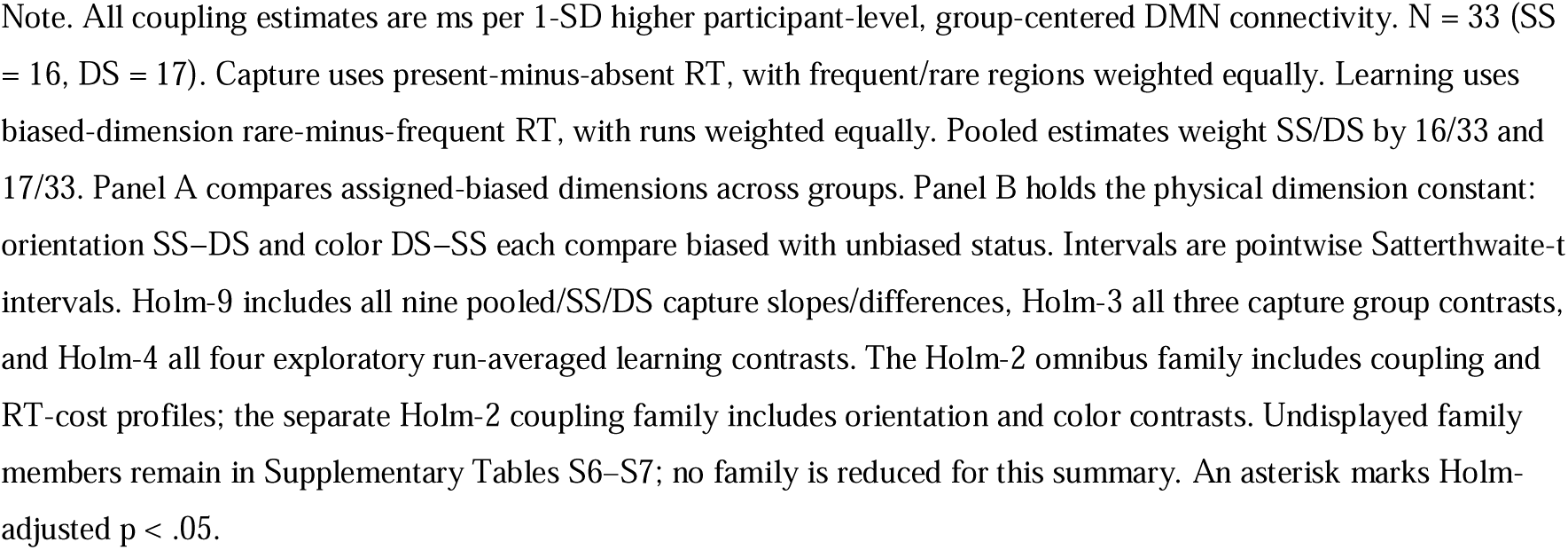
Principal DMN–behavior coupling contrasts from the hierarchical models.

| Question / coupling contrast | Scope (N) | Estimate [95% CI] | Test (df) | Raw p | Holm p | Correction family |
| --- | --- | --- | --- | --- | --- | --- |
| **A. Bias-specific distractor capture** |  |  |  |  |  |  |
| DMN–biased-dimension capture coupling | Pooled (33) | 29.00 t-based [14.93, 43.07] | t(27.59) = 4.23 | .0002 | **.0019\*** | Holm-9 |
| DMN–unbiased-dimension capture coupling | Pooled (33) | −0.53 t-based [−19.06, 17.99] | t(29.11) = −0.06 | .9533 | 1.0000 | Holm-9 |
| Coupling difference: biased – unbiased capture | Pooled (33) | 29.54 t-based [10.26, 48.82] | t(29.66) = 3.13 | .0039 | **.0234\*** | Holm-9 |
| Group difference (DS–SS): biased capture coupling | DS–SS (33) | −29.08 t-based [−57.08, −1.07] | t(27.61) = −2.13 | .0424 | .1272 | Holm-3 |
| **B. Same-physical-dimension comparison** |  |  |  |  |  |  |
| Orientation coupling: biased – unbiased (SS–DS) | Both (33) | 49.35 t-based [16.32, 82.37] | t(46.97) = 3.01 | .0042 | **.0085\*** | Holm-2 coupling |
| Color coupling: biased – unbiased (DS–SS) | Both (33) | 10.31 t-based [−20.94, 41.56] | t(49.34) = 0.66 | .5104 | .5104 | Holm-2 coupling |
| **C. Spatial-learning coupling** |  |  |  |  |  |  |
| DMN–rare-minus-frequent learning coupling | Pooled (33) | 26.48 t-based [7.74, 45.23] | t(29.00) = 2.89 | .0072 | **.0217\*** | Holm-4 |
| DMN–rare-minus-frequent learning coupling | SS (16) | 41.44 t-based [16.88, 66.00] | t(30.32) = 3.44 | .0017 | **.0068\*** | Holm-4 |
| DMN–rare-minus-frequent learning coupling | DS (17) | 12.40 t-based [−15.69, 40.50] | t(28.14) = 0.90 | .3735 | .3735 | Holm-4 |
| Group difference (DS–SS): learning coupling | DS–SS (33) | −29.04 t-based [−66.35, 8.28] | t(29.06) = −1.59 | .1224 | .2447 | Holm-4 |

Note. All coupling estimates are ms per 1-SD higher participant-level, group-centered DMN connectivity. $N = 33$ (SS = 16, DS = 17). Capture uses present-minus-absent RT, with frequent/rare regions weighted equally. Learning uses biased-dimension rare-minus-frequent RT, with runs weighted equally. Pooled estimates weight SS/DS by 16/33 and 17/33. Panel A compares assigned-biased dimensions across groups. Panel B holds the physical dimension constant: orientation SS–DS and color DS–SS each compare biased with unbiased status. Intervals are pointwise Satterthwaite- $t$ intervals. Holm-9 includes all nine pooled/SS/DS capture slopes/differences, Holm-3 all three capture group contrasts, and Holm-4 all four exploratory run-averaged learning contrasts. The Holm-2 omnibus family includes coupling and RT-cost profiles; the separate Holm-2 coupling family includes orientation and color contrasts. Undisplayed family members remain in Supplementary Tables S6–S7; no family is reduced for this summary. An asterisk marks Holm-adjusted $p < .05$ .

The SS biased-capture slope and biased-minus-unbiased slope difference survived Holm-9, whereas the corresponding DS contrasts and both groups’ unbiased slopes remained uncertain (**Supplementary Table S6C**). However, the direct DS-minus-SS biased-capture coupling difference did not survive correction (−29.08 ms/DMN SD, 95% CI [−57.08, −1.07], raw p = .0424, Holm-3 p = .1272; **Table 4A**). Full coefficient results are reported in **Supplementary Table S6**.

When physical distractor dimension was held constant, orientation coupling was stronger in biased SS than in unbiased DS (SS−DS = 49.35 ms/DMN SD, 95% CI [16.32, 82.37], raw p = .0042, Holm-2 coupling p = .0085). The color comparison, biased DS versus unbiased SS, remained uncertain (DS−SS = 10.31 ms/DMN SD, 95% CI [−20.94, 41.56], raw/Holm-2 coupling p = .5104; **Table 4B**).

### Hierarchical trial-level spatial-learning model

Higher DMN connectivity was associated with a larger rare-minus-frequent RT contrast (pooled coupling = 26.48 ms/DMN SD, 95% CI [7.74, 45.23], raw p = .0072, Holm-4 p = .0217; **Table 4C**). A 1-SD increase in connectivity corresponded to an estimated 26.48-ms larger location-dependent RT difference. Equal-run-averaged coupling was positive in SS (41.44 ms/DMN SD, 95% CI [16.88, 66.00], raw p = .0017, Holm-4 p = .0068), but uncertain in DS (12.40 ms/DMN SD, 95% CI [−15.69, 40.50], raw/Holm-4 p = .3735; **Supplementary Table S7B**). The direct DS-minus-SS learning-coupling contrast remained uncertain (−29.04 ms/DMN SD, 95% CI [−66.35, 8.28], raw p = .1224, Holm-4 p = .2447; **Table 4C**). Thus, separate subgroup significance does not establish a group difference. Full conditional coefficients and supporting group slopes are reported in **Supplementary Tables S7**.

### Network associations for distractor capture and spatial learning

#### Network context of distractor capture

Across the 28 network summaries, pooled within-DMN connectivity was positively associated with biased-dimension capture, *r*(30) = .550, 95% Fisher CI [.250, .754], raw *p* = .0011 (**Figure 4A–C**; **Table 5; Supplementary Table S8**). This association also survived permutation maxT-28 correction: unadjusted permutation *p* = .0012, maxT-28 *p* = .0044. Pooled DMN–limbic connectivity additionally survived maxT-28 (*r* = .471, unadjusted permutation *p* = .0030, maxT-28 *p* = .0419). In the exploratory SS subgroup, DMN–limbic connectivity was associated with capture, *r*(14) = .724, 95% Fisher CI [.357, .898], raw *p* = .0015; this association also survived permutation maxT-28 (unadjusted permutation *p* = .0019, maxT-28 *p* = .0263). No DS association reached unadjusted permutation *p* < .05. Direct pooled comparisons supported a stronger DMN association than the dorsal-attention association (Δ*r* = .496, unadjusted Williams *p* = .0041, Holm-6 *p* = .0248), whereas the other five comparisons did not survive their declared correction. Thus, these analyses supported the within-DMN association and one specific network contrast, rather than general DMN superiority.

**Figure 4.**
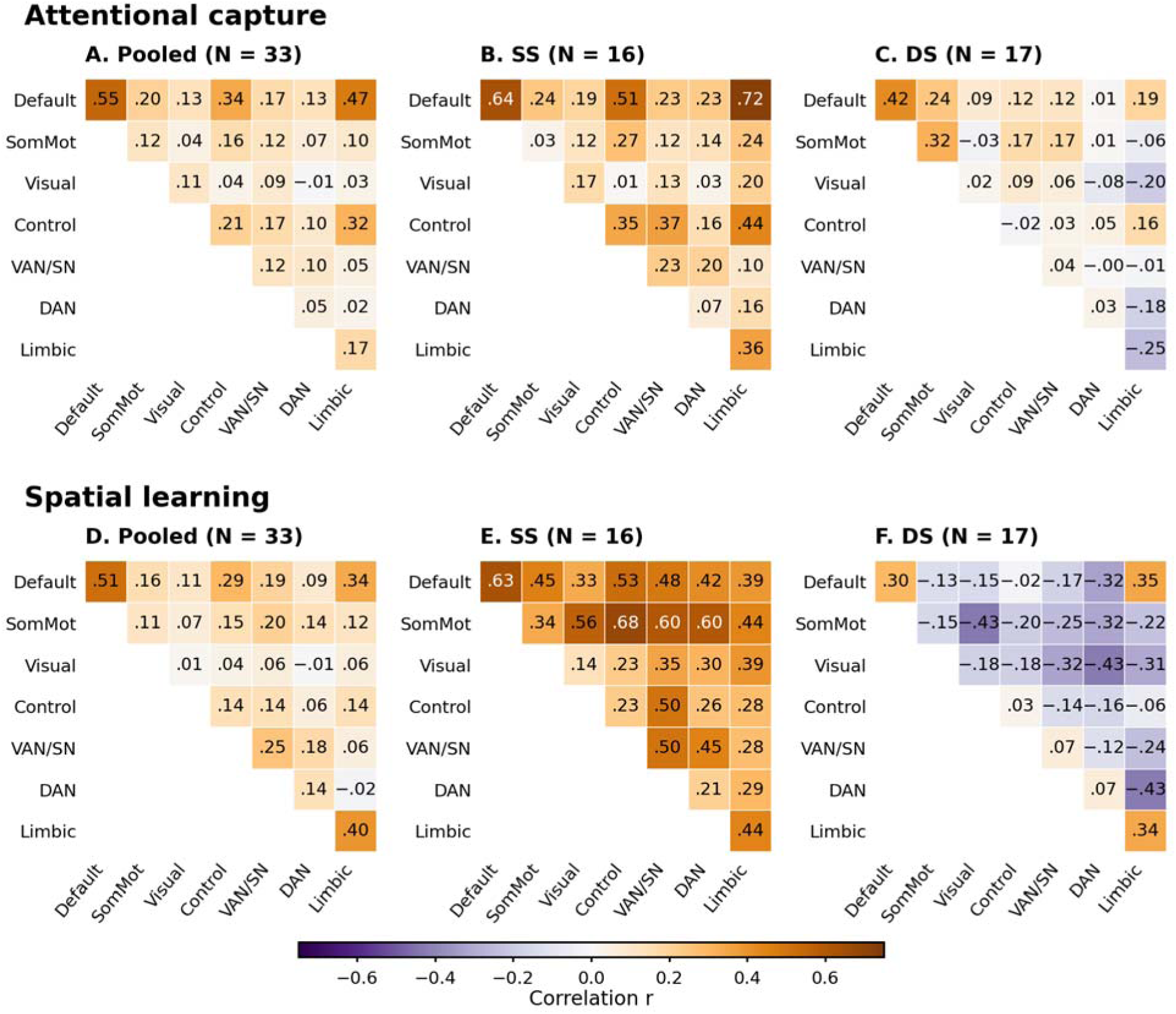
Exploratory network-summary correlations with capture and spatial learning. **A–C** show biased-present minus absent mean RT; **D–F** show biased-dimension rare-minus-frequent RT. Columns: pooled (N = 33), SS (N = 16), DS (N = 17); orientation was biased in SS and color in DS. Cells show in-sample Pearson’s r with 28 within-/between-network Yeo-7 task-run connectivity summaries (mean Fisher z). Pooled variables were centered within the group. Orange and purple denote positive and negative correlations, respectively. Default, DMN; SomMot, somatomotor; VAN/SN, ventral attention/salience; DAN, dorsal attention.

In the equal-region capture sensitivity, pooled within-DMN connectivity was also positively associated with capture, *r*(30) = .629, 95% Fisher CI [.358, .802], raw *p* = .0001; the within-group permutation test gave unadjusted *p* = .0002. Both survived their declared correction. The DMN-minus-dorsal-attention difference was significant (Δ*r* = .532, unadjusted Williams *p* = .0013, Holm-36 *p* = .0453). Group moderation of the DMN-versus-network differences was not supported (approximate bootstrap-Wald *T* = 1.86, unadjusted *p* = .9175).

#### Network context of spatial learning

For the rare-minus-frequent RT contrast, pooled within-DMN connectivity survived permutation maxT-28: *r*(30) = .513, 95% Fisher CI [.201, .731], analytic unadjusted *p* = .0027; unadjusted permutation *p* = .0042, maxT-28 *p* = .0215 (**Figure 4D–F**; **Table 5; Supplementary Tables S10**). The subgroup within-DMN estimates were also positive: SS, *r*(14) = .632, CI [.199, .859], analytic unadjusted *p* = .0086; DS, *r*(15) = .296, CI [−.215, .680], analytic unadjusted *p* = .2483. Neither survived panel-wise correction. None of the six direct pooled DMN-versus-network comparisons survived their declared correction; the smallest unadjusted Williams *p* was .0155 for DMN versus visual connectivity (Δ*r* = .500).

**Table 5.** Network-summary associations.

| Outcome / sample | Network pair | r | Permutation p<br>Unadjusted | Permutation p<br>Holm-28 | Permutation p<br>maxT-28 |
| --- | --- | --- | --- | --- | --- |
| Capture / pooled | DMN–DMN | 0.550 | 0.0012 | **0.0336** | **0.0044** |
| Capture / pooled | DMN–Limbic | 0.471 | 0.0030 | 0.0810 | **0.0419** |
| Capture / SS | DMN–Limbic | 0.724 | 0.0019 | 0.0532 | **0.0263** |
| Learning / pooled | DMN–DMN | 0.513 | 0.0042 | 0.1176 | **0.0215** |

Note. Rows were selected solely because permutation maxT-28 $p < .05$ ; all 168 associations appear in Supplementary Table S8, S10. Every p column is permutation-based. Holm-28 and maxT-28 are separate corrections over all 28 network pairs within each outcome/sample panel. Pooled N = 33 with SS/DS group centering; SS N = 16. Capture uses the observed trial mixture; spatial learning is biased-dimension rare-minus-frequent RT. Bold adjusted p values indicate $p < .05$ .

### Signed-edge enrichment

Positive edge–behavior associations were descriptively concentrated within the DMN for both outcomes. For capture, 226 within-DMN edges passed the exploratory screen among 4,095 possible edges, compared with 41.88 expected from network size (observed/expected = 5.40; unadjusted permutation *p* = .0015, maxZ-56 *p* = .0761). For spatial learning, the corresponding count was 171, compared with 26.72 expected (observed/expected = 6.40; unadjusted permutation *p* = .0029, maxZ-56 *p* = .1558).

The overlapping DMN-touching category, comprising edges with at least one DMN endpoint, also showed nominal enrichment for capture (583 observed versus 328.51 expected; observed/expected = 1.77; unadjusted permutation *p* = .0158, maxZ-14 *p* = .1616) and learning (383 observed versus 209.57 expected; observed/expected = 1.83; unadjusted permutation *p* = .0303, maxZ-14 *p* = .3145). Across all 140 category/sign/outcome tests, ten had unadjusted permutation *p* < .05, but none survived its outcome-specific correction. These results indicate descriptive concentration of positive associations around the DMN, without establishing corrected enrichment or general DMN specificity (**Table 6**; **Figure 5; Supplementary Tables S9–S10**).

**Table 6.** DMN categories in pooled signed-edge enrichment.

| Outcome / category | O | E | O/E | Permutation p | Permutation p | Permutation p |
| --- | --- | --- | --- | --- | --- | --- |
|  |  |  |  | Unadjusted | Holm-56 | maxZ-56 |
| Capture / within-DMN | 226 | 41.88 | 5.40 | 0.0015 | 0.0840 | 0.0761 |
| Capture / DMN-touching | 583 | 328.51 | 1.77 | 0.0158 | 0.2212 | 0.1616 |
| Learning / within-DMN | 171 | 26.72 | 6.40 | 0.0029 | 0.1624 | 0.1558 |
| Learning / DMN-touching | 383 | 209.57 | 1.83 | 0.0303 | 0.4242 | 0.3145 |

Note. All rows describe positive edge–behavior associations and are selected for their relevance to the focal DMN hypothesis, not corrected significance. O = observed count; E = capacity-based expected count; O/E is descriptive. Every p column is permutation-based. Holm and maxZ use 56-test families for network pairs and 14-test families for overlapping network-touching categories, separately for capture and learning. Table S9-S10 reports all tests.

**Figure 5.**
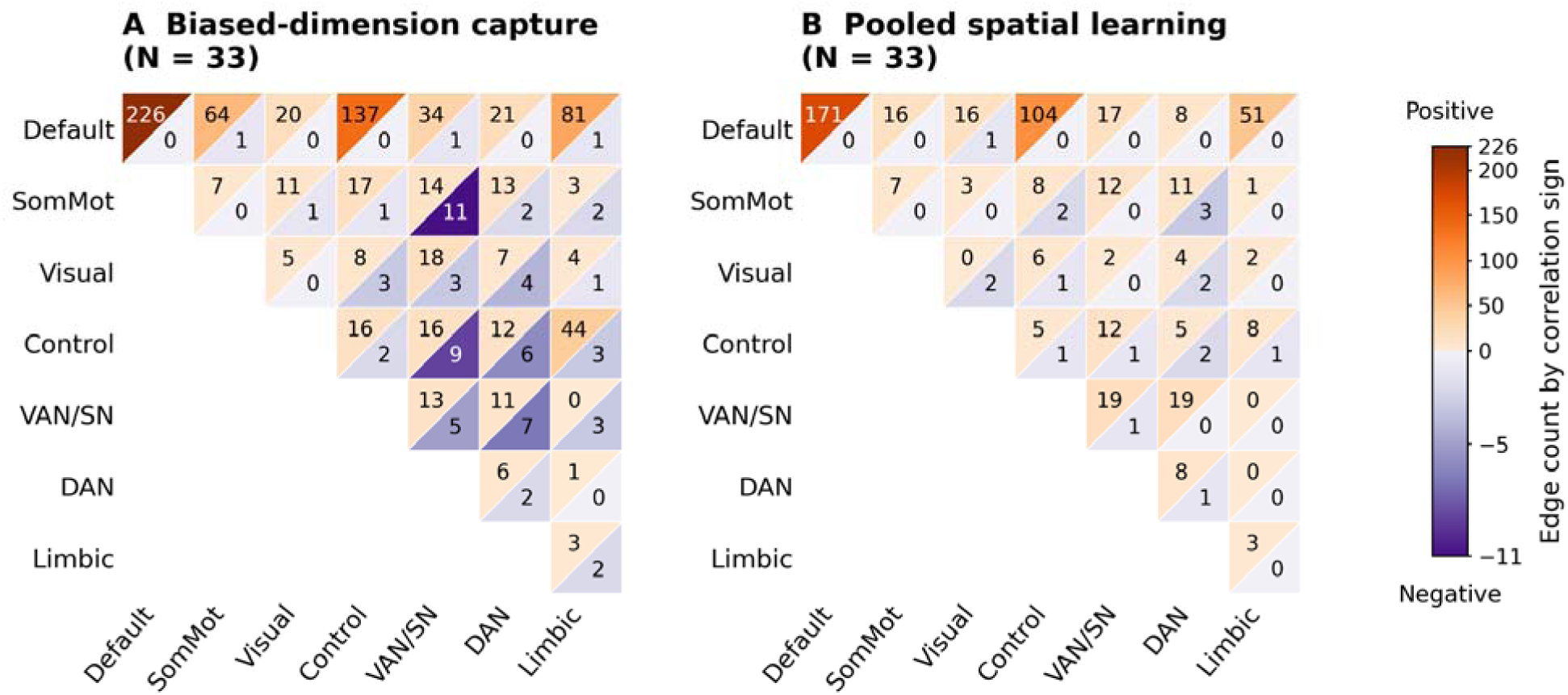
Descriptive signed edge counts for biased-dimension capture and spatial learning. Panels show pooled associations for **(A)** biased-dimension capture and **(B)** biased-dimension spatial learning (N = 33). Capture is biased-present minus absent mean response time (RT); learning is rare-minus-frequent RT on biased-dimension trials. Connectivity and behavioral scores were centered within SS and DS before an uncorrected two-sided p < .01 screen of 79,401 possible edges. Capture retained 812 positive and 70 negative edges; learning retained 518 positive and 18 negative edges. Within each network-pair cell, orange upper-left triangles indicate positive counts and purple lower-right triangles indicate negative counts. Separate color scales are used for each sign and panel. Counts depend on network-pair size and do not represent corrected edge discoveries, enrichment or prediction.

### Exploratory connectome-based prediction

Exploratory connectome-based prediction assessed whether connection patterns within and between DMN subdivisions could estimate attentional capture or spatial learning in participants held out from model fitting. Held-out prediction of distractor capture was strongest for connectivity within the Yeo-17 DefaultB subdivision (held-out r = .619, raw p = .0012, Holm-6 p = .0072; **Supplementary Results “Exploratory connectome-based prediction”, Figure S1**), comprising predominantly lateral temporal, inferior-parietal, and prefrontal DMN territories in the present Schaefer/Yeo parcellation [29, 30]. No corrected predictive effect was obtained for spatial learning.

## Discussion

The present study asked whether within-DMN task-run connectivity was associated with distractor capture, whether this relationship was stronger for the biased distractor dimension, and whether DMN connectivity was also related to location-dependent modulations of capture. Across complementary analyses, higher within-DMN connectivity was associated with greater distractor capture by the biased dimension. In contrast, the corresponding association for the unbiased dimension was weak, and the direct biased-vs.-unbiased comparison supported a stronger association for the biased dimension. This pattern was also evident when the two dimensions were evaluated under the same frequent-/rare-region weighting. Higher within-DMN connectivity was also associated with a larger rare-minus-frequent location RT difference for the biased dimension, although this difference was closely tied to the overall magnitude of capture. Together, these findings indicate that the DMN– distractor-capture association was not expressed uniformly across distractors, but was strongest for the biased dimension, the one embedded in the spatial-probability structure of the task.

The participant-level difference scores and hierarchical trial-level model reached the same conclusion. The trial-level model jointly represented distractor-absent, biased-frequent, biased-rare, unbiased-frequent, and unbiased-rare trials and estimated how distractor-related RT costs changed with DMN connectivity while accounting for repeated observations within participants without requiring equal trial counts. Biased and unbiased coupling could therefore be compared within a common model and under an identical 50/50 weighting of frequent and rare regions. Under this standardized spatial mixture, biased coupling was positive and exceeded unbiased coupling. Thus, the trial-level analysis converged with the participant-level results and showed that the bias-specific association was not simply a consequence of comparing different spatial-region mixtures. The weighting standardizes the estimand, however, rather than the underlying data: biased and unbiased conditions still differed in exposure and in the amount of trial information contributing to their estimates.

The evidence that DMN connectivity was more strongly related to biased-than unbiased-dimension capture was not uniform across analyses. Although the participant-level correlation comparison and the principal trial-level hierarchical model supported this difference, the corresponding contrast in the three-condition trial-history-adjusted model did not survive multiple-comparison correction. These models differed in both their treatment of trial history and their representation of frequent and rare distractor locations, so the difference in statistical outcomes cannot be attributed specifically to adjustment for prior exposure. The positive association with biased-dimension capture itself, by contrast, held across these analyses, including trial-history adjustment, conditional trial-count matching, and equal-region weighting.

The physical-dimension analyses suggest that the connectivity pattern followed, at least in part, assignment to the spatially biased dimension, beyond intrinsic differences between orientation and color distractors. Because orientation carried the spatial bias in SS and color carried it in DS, each physical dimension appeared as biased in one group and unbiased in the other. Orientation showed stronger DMN– distractor-capture coupling in biased SS than in unbiased DS, whereas the corresponding color comparison remained uncertain. Thus, the orientation result supports a contribution of assigned bias status beyond physical dimension and differences in the spatial-region mixture of the analyzed trials, although the asymmetric evidence across dimensions limits generalization.

DMN connectivity was also associated with spatial learning, operationalized as the rare-minus-frequent RT difference for biased-dimension distractors, evident at both the participant level and trial level. Because the same distractor-absent baseline cancels in this contrast, the learning score equals the difference between capture at rare and frequent locations. Capture and spatial learning therefore characterize related aspects of distractor interference: its overall magnitude and its spatial gradient. With higher DMN connectivity, biased distractors produced more interference at both frequent and rare locations, and in the trial-level model the increase was larger at rare locations, yielding a steeper rare-minus-frequent gradient. This gradient, however, was closely tied to the overall magnitude of capture. Participants with larger overall capture also showed larger rare-minus-frequent RT differences, the participant-level associations at rare and frequent locations did not differ reliably, and after accounting for overall capture, the association between DMN connectivity and the spatial difference was much weaker and no longer significant after correction. The steeper gradient at higher connectivity is therefore best understood as part of the larger overall interference.

In the broader network analyses, the within-DMN association with distractor capture was the only pooled network association to survive Holm correction across all 28 within– and between-network summaries. The DMN association was also stronger than the corresponding dorsal-attention-network association, although the remaining direct network comparisons were not supported. The equal-region capture analysis showed the same general pattern. Thus, the broader screen strengthens the evidence that the DMN association was not simply one of many equally strong networks–capture associations, while falling short of demonstrating a unique or general superiority of the DMN over all other networks. For spatial learning, by contrast, the pooled within-DMN spatial-learning association also survived permutation correction (maxT-28), and direct comparisons did not establish a stronger association than for other networks. Network specificity is therefore better supported for distractor capture generally than for spatial learning.

The between-network analyses suggest that the connectivity–capture relationship may extend beyond the DMN. Stronger DMN–limbic connectivity was associated with greater biased-dimension capture in the pooled sample and the SS subgroup, with both associations surviving permutation-based maximum-statistic correction across the 28 network pairs in their respective panels. These findings implicate coupling between the DMN and orbitofrontal and anterior temporal territories, including the temporal poles, that constitute the Yeo limbic network [30, 33]. The pooled within-DMN spatial-learning association also survived this correction. These findings suggest that the capture association extends to coordination between the DMN and orbitofrontal–anterior temporal regions. This interpretation is consistent with evidence that substantial portions of the Yeo limbic network form an extension of the DMN [33]. Given the roles of orbitofrontal cortex in representing inferred task states and anterior temporal cortex in conceptual knowledge [34, 35], stronger coupling could accompany a greater influence of learned representations on attentional selection. In the present task, this might involve expectations concerning the biased distractor dimension and its spatial distribution.

The exploratory CPM analysis pointed in the same direction: held-out prediction of distractor capture was strongest for connectivity within the Yeo-17 DefaultB subdivision, comprising predominantly lateral temporal, inferior-parietal, and prefrontal DMN territories in the present Schaefer/Yeo parcellation [29, 30]. The DefaultB result survived the primary correction but was sensitive to the permutation-tail definition, and no corrected predictive effect was obtained for spatial learning. The CPM findings therefore provide suggestive rather than independent support for the stronger DMN-related pattern observed for distractor capture.

These associations imply that stronger within-DMN coupling was not optimal for the task: participants with higher connectivity lost more time when a biased-dimension distractor appeared, whereas lower connectivity went with less capture. This cost was not offset by faster responding on distractor-absent trials. Two features of the cost constrain its explanation. First, it was confined to the biased dimension: unbiased-dimension capture was unrelated to connectivity, even in DS, where unbiased orientation distractors produced the largest capture of all. A general weakening of attentional control, or more frequent lapses of attention [21], would be expected to increase capture regardless of dimension. Second, the cost arose at both frequent and rare locations, increasing most at rare locations, where the learned spatial expectation was violated. Higher connectivity was therefore not associated with poorer performance in general, but with less effective handling of the one distractor type that the task most required participants to filter out.

These features can be explained without assuming that the DMN represents the spatial context itself. In the present task, the context is given by the task environment: biased-dimension distractors were the most frequent distractors overall and appeared predominantly in the frequent region, so the optimal task set includes filtering out this particular distractor [2, 13]. On this account, high within-DMN connectivity during the task indexes a state that interferes with implementing this task set [21, 22], so that biased-dimension distractors are filtered less effectively. This explains why the connectivity-related cost was confined to the biased dimension: unbiased distractors were less frequent and not the main target of the filtering set, leaving little for connectivity to impair. It also explains why capture was increased at frequent as well as at rare locations. The cost was nevertheless steepest for rare locations, where filtering may depend more strongly on the task set, whereas at frequent locations the learned suppression of that region provides additional shielding [9, 11] – resulting in a steeper rare-minus-frequent gradient. A recent suppression-threshold framework [36] proposes that learned distractor statistics adjust the threshold for triggering suppression: a lower threshold makes suppression routines easier to engage, whereas a higher threshold reduces the risk of inappropriately suppressing targets. Extending this account to the present task, learned spatial regularities would make suppression more readily engaged at frequent locations, providing greater protection against biased-dimension distractors there than at rare locations. Weaker task control, associated with increased within-DMN connectivity, would impair the invoking of suppression routines in both regions while preserving this relative protection. If the strength of task control (modulating readiness to engage suppression) and distractor interference are related nonlinearly, the same weakening could produce a larger increase in interference at rare locations – amplifying the rare-minus-frequent difference without implying stronger spatial learning. Evidence linking particular DMN connections to behavioral variability and task-related reductions in connectivity to faster performance provides a broader context for this interpretation [37, 38]. An alternative view links the DMN to context memory and to the integration of current processing with prior experience [17, 18, 19, 39, 40, 41, 42], suggesting that higher connectivity reflects a greater influence of learned regularities on selections. This view, however, does not explain why capture also increased at frequent locations. These results remain correlational and cannot determine whether high within-DMN connectivity disrupts the implementation of the task set or instead results from a less well implemented task set.

Behavioral reliability varied across measures and groups, which qualifies the individual-difference interpretation. SS orientation/biased capture showed positive consistency and run-to-run stability, whereas estimates for DS color/biased capture were less certain. Spatial-learning scores likewise showed clearer reliability in the pooled sample and in SS than in DS. This issue is important because individual-difference correlations can be attenuated when the underlying behavioral measures are imprecise, even when the experimental effect itself is robust [43]. The hierarchical models reduce reliance on aggregated difference scores by using trial-level RT information and explicitly modeling repeated observations, but they do not eliminate uncertainty in the participant-level connectivity measure or establish that the observed behavioral differences represent stable traits. The findings are therefore best interpreted as associations among measures obtained during the present task, rather than as estimates of enduring individual characteristics.

Several additional features constrain generalization. The imaging sample was modest, particularly for within-group analyses, and all principal and exploratory analyses were based on the same participants. The physical-dimension comparisons therefore provide internal specificity tests rather than independent replications. Likewise, the broader network, edge-level, and CPM analyses provide complementary characterization of the same dataset rather than external validation. Future studies with larger independent samples and repeated sessions will be needed to establish the stability and generalizability of the connectivity–behavior relationships. Most importantly, designs that manipulate spatial regularity independently of dimension exposure would be required to determine whether spatial learning itself, rather than the associated exposure structure, drives the stronger connectivity relationship for the biased dimension.

Overall, the results link within-DMN task-run connectivity to two aspects of behavior under spatially structured distraction. The strongest evidence concerned distractor capture: higher DMN connectivity was associated with greater interference from the biased distractor dimension, and this bias-specific relationship held across participant-level and hierarchical trial-level analyses and when spatial-region weighting was standardized. The broader network and exploratory CPM analyses provided additional support for this pattern. DMN connectivity was also related to spatial learning, but this association was closely tied to overall capture. Together, the findings suggest that stronger within-DMN connectivity during the task marks greater susceptibility to distractors that the task set was tuned to filter out, consistent with less effective implementation or maintenance of that filtering set. Whether high connectivity disrupts the implementation of the task set, or follows from a less well implemented one, remains to be tested.

## Declarations

### Funding

This work was supported by German Research Foundation (DFG) grants CH3093/1-1 and SH 166/10-1, awarded to SC and ZS, respectively, and used the NICUM Siemens Prisma scanner supported by DFG grant INST 86/1739-1 FUGG.

### Competing interests

The authors declare no competing interests.

### Ethical approval

The original data collection was approved by the Ethics Committee of the Department of Psychology, LMU Munich. The present study reanalyzed previously collected data and involved no new recruitment or data collection.

### Consent to participate

Participants provided written informed consent.

### Consent to publish

Not applicable. The manuscript contains no identifiable individual-level information.

### Data and code availability

The task-fMRI source dataset analyzed in this study is publicly available from G-Node Open Data: Shi et al. [25], Dataset: Retinotopic mapping of statistical learning of distractor suppression in visual search, https://doi.org/10.12751/g-node.zfrem9 (CC BY 4.0).

The analysis code is publicly available at https://github.com/msenselab/DMN-attentional-capture-and-spatial-learning.

### Authors’ contributions

SC, HJM and ZS: conceptualization and methodology. SC: formal analysis and writing – original draft. HJM and ZS: writing – review and editing.

## Notes

### Competing Interest Statement

The authors have declared no competing interest.

### Summary of Updates

More details have been clarified, including methods and results.

