## Supplementary for "Default Mode Network Connectivity Is Associated with Capture by a Distractor Dimension Carrying Learned Spatial Regularity"

### Supplementary Methods

#### Scope and reading guide

The supplementary material follows the behavioral and connectivity analyses in the main text. **Supplementary Equations S1–S14** provide definitions and procedures for behavioral scores, participant-level associations, robustness analyses, and hierarchical models. Supplementary Methods S15–S16 describe network-level associations, edge enrichment, and exploratory connectome-based prediction. The Supplementary Results present exploratory connectome-based prediction results (Figure S1). The supplementary tables provide behavioral tests (Table S1), reliability analyses (Table S2), additional participant-level DMN associations (Table S3), and preprocessing sensitivity checks (Table S4). Tables S5–S7 report trial-history sensitivity and hierarchical capture and spatial-learning models, whereas Tables S8–S10 present network-summary associations, signed-edge enrichment, and spatial-learning network summaries and edge counts, respectively. SS denotes the same-dimension spatial-bias group (orientation biased), and DS the different-dimension spatial-bias group (color biased). Biased and unbiased dimensions refer to their assigned spatial-bias status rather than a fixed physical distractor dimension. DMN denotes the default-mode network. Unless otherwise stated, confidence intervals are pointwise 95% intervals.

#### Supplementary Equation S1. Participant-level attentional capture and behavioral spatial learning

Notation: i indexes participants, j trials, g(i) group, and h half or run. β denotes fixed regression coefficients. Superscripts such as cap, hist, OLS, loc, run and mix identify separately fitted models and do not imply that their coefficients are equal. Random effects and residuals retain distinct symbols. Indices and coefficient labels are defined beside each model.

All scores use correct responses with RTs of 200–3,000 ms inclusive. For participant i, Āᵢ is the absent-trial mean RT, and *P̄*_B,i_ and *P̄*_U,i_ are the biased- and unbiased-dimension present-trial means. Means are computed from the corresponding eligible trials, in milliseconds.

##### Participant-level attentional capture:

*C*^B^_i_ = *P̄*_B,i_ − *Ā*_i_

*C*^U^_i_ = *P̄*_U,i_ − *Ā*_i_

C denotes a participant capture score, whereas lower-case b and u below denote trial indicators. Positive capture indicates distractor-related slowing. Each within-group capture score is tested against zero using a two-sided one-sample t-test. Between-group means are compared using Welch tests.

#### Participant-level count-weighted combined capture score:

*C*^all^_i_ = (*n*_B,i_*C*^B^_i_ + *n*_U,i_*C*^U^_i_) / (*n*_B,i_ + *n*_U,i_)

The weights are each participant’s retained biased- (*n*_B,i_) and unbiased-present (*n*_U,i_) trial counts.

Participant-level spatial learning:

*S*_i_ = *RT̄*_Rare,i_ − *RT̄*_Frequent,i_

**Supplementary Equation S2. Training-plus-scanner behavioral model**

*RT*_ij_ = *β*^beh^_0_ + *β*^beh^_K_*K*_ij_ + *β*^beh^_L_*L*_ij_ + *β*^beh^_D_*D*_i_ + *β*^beh^_LD_*L*_ij_*D*_i_ + *a*^beh^_0i_ + *ε*^beh^_ij_

This separate behavioral model uses eligible biased-dimension training and scanner trials from 35 participants. K is the saved continuous block index (1–14), L codes frequent = 0 and rare = 1, and D codes SS = 0 and DS = 1. The participant random intercept is Gaussian; trial residuals are Gaussian. REML and Satterthwaite inference are used. Participant-balanced SS and DS learning tests form a separate Holm-2 family.

#### Supplementary Equation S3. Reliability of capture and spatial-learning scores

For condition *d* and occasion *h* (recorded odd/even half or scanner run), the capture score uses the absent baseline from that same occasion. The condition index is retained throughout.

Capture score: *C*_idh_ = *P̄*_idh_ − *Ā*_ih_

Group centering: *C*^c^_idh_ = *C*_idh_ − *C̄*_g(i),d,h_

Correlation: *r*_d,split_ = cor(*C*^c^_id1_, *C*^c^_id2_)

Spearman–Brown projection: *r*_SB,d_ = (2*r*_d,split_) / (1 + *r*_d,split_)

Group centering is repeated separately within SS and DS for each occasion. Uncentered pooled capture correlations are reported separately because they can also reflect group mean differences. Within-group correlations use the original scores. The pooled group-adjusted test uses df = N − 3; ordinary within-group tests use df = N − 2. The Spearman–Brown projection applies to approximately parallel split halves. Spatial-learning reliability replaces capture with the rare-minus-frequent RT contrast computed within each occasion. Recorded parity is assigned before exclusions, without renumbering eligible trials.

Each splitting scheme has separate Holm-2 pooled-capture and Holm-4 within-group-capture families; all six learning tests share Holm-6. Two-sided Pearson tests use df = N−3 for pooled and N−2 for within-group estimates. Bootstrap intervals and parametric p values can differ in zero-exclusion decisions. Full-behavioral-sample and uncentered capture sensitivities remain in Table S2.

**Supplementary Equation S4. Primary DMN–behavior analyses**

A. **Group-centered associations and connectivity scaling**

F is the equal-run mean Fisher-z connectivity across the 4,095 unique edges among the 91 covered Yeo-7 Default parcels. For participant i in group g(i), define:

*F*^c^_i_ = *F*_i_ − *F̄*_g(i)_

*V*^c^_i_ = *V*_i_ − *V̄*_g(i)_

*r*_FV_ = cor(*F*^c^, *V*^c^)

*X*_i_ = (*F*^c^_i_) / (*s*_Fᶜ_)

V is a specified behavioral score: biased, unbiased or combined capture, or the scanner rare-minus-frequent learning contrast. For the pooled imaging sample (N = 33, SS = 16, DS = 17), removing the one SS–DS contrast gives df = N − 3 = 30 and Fisher-z standard error 1/√(N − 4). Within-group correlations use df = N − 2 and Fisher-z standard error 1/√(N − 3). The SD in X is calculated across the 33 unique participants after group centering, before joining connectivity to trial rows. The focal pooled DMN–biased-capture test is unadjusted. Eight capture associations retain their Holm-8 family; pooled, SS and DS learning correlations form a separate Holm-3 family.

##### B. Physical-dimension mapping and decomposition of the pooled correlation:

This analysis aims to distinguish physical-dimension scores from biased/unbiased assignment, and it shows each group’s contribution to the pooled, group-centered correlation. *C*_ori,i_ and *C*_col,i_ are observed orientation-present and color-present mean RTs minus participant *i*’s absent-trial mean (ms). *C*^B^_i_ and *C*^U^_i_ relabel them as assigned-biased and assigned-unbiased capture using *D*_i_. Orientation is biased in SS and unbiased in DS; color has the reverse assignment. These observed-mixture scores differ from the equal-region scores *C*^B50^_i_ and *C*^U50^_i_ above.

*C*^B^_i_ = (1 − *D*_i_)*C*_ori,i_ + *D*_i_*C*_col,i_

*C*^U^_i_ = (1 − *D*_i_)*C*_col,i_ + *D*_i_*C*_ori,i_

*r*_F,ori_ = cor(*F*^c^, *C*^c^_ori_), *r*_F,col_ = cor(*F*^c^, *C*^c^_col_)

*q*_g_ = (*∑*_i:g(i)=g_*F*^c^_i_*C*^B,c^_i_) / (√(*∑*_i_(*F*^c^_i_)^2^ *∑*_i_(*C*^B,c^_i_)^2^))

*r*_FB_ = *q*_SS_ + *q*_DS_, *share*_g_ = (*q*_g_) / (*r*_FB_)

*F* denotes the DMN connectivity measure defined in S4. The superscript ^c^ denotes centering within SS and DS before pooling: *F*^c^, *C*^c^_ori_ and *C*^c^_col_ are the centered connectivity, orientation-capture and color-capture vectors. *r*_F,ori_ and *r*_F,col_ are their pooled Pearson correlations; the two physical-score tests retain Holm-2 (Table S3B). The superscript ^B,c^ in *C*^B,c^_i_ identifies a centered biased-capture score, and *F*^c^_i_ is participant *i*’s centered connectivity.

Let *g(i)* denote participant *i*’s SS/DS group. *F*^c^_i_ and *C*^B,c^_i_ are values centered within *g(i)*. For *g* ∈ {SS, DS}, *q*_g_ is group *g*’s normalized centered cross-product contribution to the pooled group-centered correlation *r*_FB_. Its numerator includes only that group; both denominator sums extend over all 33 participants. Thus *r*_FB_ = *q*_SS_ + *q*_DS_ and, when *r*_FB_ is nonzero, *share*_g_ is *q*_g_ divided by *r*_FB_: group *g*’s signed share of the pooled centered covariance/cross-product.

**Supplementary Equation S5. Direct dependent-correlation comparisons: Williams test**

For two correlations sharing connectivity, let *a* denote the connectivity–first-outcome correlation, *b* the connectivity–second-outcome correlation, and c the correlation between the outcomes. For biased versus unbiased capture these are rFB, rFU and rBU, respectively. All three pooled correlations use the same group-centered participants.

K = 1 − *a*^2^ − *b*^2^ − *c*^2^ + 2abc

*J*_W_ = (2K(n − 1)) / (n − 3) + ((a + b)^2^) / (4)(1 − c)^3^

*t*_W_ = ((a − b)√((n − 1)(1 + c))) / (√(*J*_W_))

The group-adjusted approximation uses effective n = 32 for all 33 participants, giving df = n − 3 = 29. A positive difference indicates a more positive association with the first outcome. Full-score and mean count-matched biased-versus-unbiased comparisons retain their separate Holm-2 family. The averaged score estimates the original full-score mean in expectation. The capture-versus-learning comparison is a single explicitly exploratory test; equal-region comparisons retain their own family. This compares correlations, not millisecond slopes.

#### Supplementary Equation S6. DMN connectivity and distractor capture: partial associations

Partial correlations assess the linear association between DMN connectivity and biased-dimension capture after adjusting both measures for the same control variables. Each observation is one participant. For participant i, Fᵢ is the Default91 DMN connectivity measure defined in S4, Vᵢ is biased-dimension capture, Uᵢ is unbiased-dimension capture, Pᵢ is inter-trial priming (the prior-display contrast), and Gᵢ is the group indicator (SS = 0; DS = 1).

For the pooled analysis controlling for unbiased capture and group, two ordinary least-squares (OLS) regressions are fitted separately:

*F*_i_ = *a*_F_ + *b*_F_*U*_i_ + *c*_F_*G*_i_ + *e*_Fi_

*V*_i_ = *a*_V_ + *b*_V_*U*_i_ + *c*_V_*G*_i_ + *e*_Vi_

The coefficients a_F_ and a_V_ are the respective intercepts. The coefficients b_F_ and b_V_ describe adjustment for unbiased capture, and c_F_ and c_V_ describe adjustment for group. Each regression has its own estimated coefficients. The residuals e_Fi_ and e_Vi_ are the observed values minus their respective fitted values. Q collects the control variables into a regression design matrix. It has one row per participant, a column of ones for the intercept, and one column for each control variable. In this example, participant i contributes the row [1, Uᵢ, Gᵢ]. N denotes the number of participants, and k counts the control variables without the intercept. The β coefficient vector in the F regression contains [a_F_, b_F_, c_F_], and the separate β vector in the V regression contains [a_V_, b_V_, c_V_]. Each vector includes its own intercept and control-variable slopes. Full rank means that no column of Q is an exact linear combination of the other columns; the control variables need not be uncorrelated. The compact form is:

F = Q*β*^adjF^_Q_ + *e*^F^

V = Q*β*^adjV^_Q_ + *e*^V^

*r*_FV|Q_ = cor(*e*^F^, *e*^V^)

The partial correlation r_FV|Q_ is the Pearson correlation between the residual vectors e_F_ and e_V_. It describes the remaining linear association after the same covariates have been adjusted for in both measures. The common Q specifies the same participants and controls, not shared regression coefficients.

For the pooled priming-adjusted analysis, P replaces U, so the row of Q is [1, Pᵢ, Gᵢ]. Each pooled partial analysis therefore controls two variables (k = 2). Within SS or DS, group is constant and is omitted: the row is [1, Uᵢ], with one control variable (k = 1).

The corresponding two-sided Pearson test uses df = N − k − 2, with the intercept excluded from k. These partial associations remain in the original eight-test exploratory capture-association family (Holm-8; Supplementary Table S3A).

#### Supplementary Equation S7. Participant-bootstrap confidence intervals

Let F_i_ be participant i’s mean within-DMN Fisher-z connectivity and C_i_ the behavioral score being evaluated. For capture, C_i_ is biased-, unbiased-, or count-weighted combined capture (S1); the same participant-resampling principle applies to spatial-learning scores. Within each group g ∈ {SS, DS}, sample n_g_ participant indices I_gk_^(b)^ independently with replacement, retaining connectivity and all behavioral scores together. Group sizes remain SS = 16 and DS = 17 in each of B = 10,000 resamples. Recalculate the connectivity and outcome group means from each resampled group, including repeated participants, before pooling:

I_gk_^(b)^ ∼ Uniform(G_g_), k = 1, …, n_g_, b = 1, …, B

F_gk_^*(b)^ = F_Igk_^(b)^, C_gk_^*(b)^ = C_Igk_^(b)^

F̄_g_^*(b)^ = (1/n_g_)∑_k=1_^n^_g_F_gk_^*(b)^, C̄_g_^*(b)^ = (1/n_g_)∑_k=1_^n^_g_C_gk_^*(b)^

r^*(b)^ = cor({F_gk_^*(b)^ − F̄_g_^*(b)^}, {C_gk_^*(b)^ − C̄_g_^*(b)^})

CI_95%_ = [Q_.025_({r^*(b)^}), Q_.975_({r^*(b)^})]

G_g_ is the observed participant-index set for group g; braces in the correlation collect all resampled SS and DS participants. Q_q_ is the empirical q quantile. The index k labels sampled positions within group g, and b labels resampling iterations. These are paired percentile intervals, not bias-corrected and accelerated intervals. For a difference between correlations sharing connectivity, compute both correlations from the same participant resample and take quantiles of their difference. Within-group intervals use the corresponding group’s resampled participants without pooling.

#### Supplementary Equation S8. Within-group permutation test for biased capture

For the primary capture association, connectivity and group membership were held fixed while biased-capture scores were shuffled independently within SS and DS [40]. Each of B = 10,000 permutations preserved both group sizes and each group’s observed score distribution. The permuted outcome and fixed connectivity were centered within group before calculating the pooled correlation:

C_i_^π(b)^ = C_πg(i)_^(b)^_(i)_

r_b_^π^ = cor({F_i_ − F̄_g(i)_}, {C_i_^π(b)^ − C̄_g(i)_^π(b)^})

p_perm_ = [1 + ∑_b=1_^B^ 𝟙(|r_b_^π^| ≥ |r_obs_|)]/(B + 1)

π_g_^(b)^ denotes the random rearrangement of participant indices within group g in permutation iteration b; its superscript labels the iteration, not a power. C_i_ denotes biased capture in this section. r_obs_ is the observed group-centered correlation (S4), and 𝟙 denotes the indicator function. The two-sided absolute-correlation tail and Monte Carlo add-one correction were used. Because only labels within a group are permuted, the group mean is unchanged; centering is nevertheless applied to each permuted score vector.

#### Supplementary Equation S9. Participant influence: leave-one-participant-out analysis

Each of the 33 participants was omitted in turn. For every omission, connectivity and biased capture were re-centered using the remaining participants in the corresponding SS or DS group, and the pooled correlation was recalculated:

F_j,−i_^c^ = F_j_ − F̄_g(j),−i_, C_j,−i_^c^ = C_j_ − C̄_g(j),−i_

r_−i_ = cor_j≠i_(F_j,−i_^c^, C_j,−i_^c^)

The subscript −i denotes quantities computed after participant i’s removal; r_−i_ is the corresponding pooled correlation. We inspected the range and direction of the 33 omission estimates to assess whether an individual participant drove the association. Where reported, omission-specific two-sided parametric tests use df = (N − 1) − 3, and approximate Fisher-z intervals use SE = 1/√[(N − 1) − 4], with N = 33. These pointwise summaries assess influence on the same-sample association.

#### Supplementary Equation S10. Connectivity-preprocessing sensitivity: nuisance-regression variants

Sensitivity variants used the same 399 common-coverage Schaefer parcels (native label 323 excluded uniformly), Yeo-7 network assignment, retained-volume mask, detrending, 0.01–0.10-Hz filtering, standardization and equal two-run aggregation. Only the nuisance-regression design varied. The primary design contained 24 motion terms (six motion parameters, their derivatives and the squares of both sets), white-matter signal and CSF signal. The four alternatives omitted nuisance regression, added global signal, added three task regressors, or added both task and global-signal regressors:

Z_primary_ = [M_24_, WM, CSF], Z_none_ = ∅

Z_GSR_ = [Z_primary_, GS], Z_task_ = [Z_primary_, T]

Z_task+GSR_ = [Z_primary_, GS, T]

T_c_(t) = (h_Glover_ ∗ d_c_)(t), c ∈ {Absent, Different, Same}

Each d_c_(t) is the recorded display-event time course, using the event onsets and durations for absent, color-distractor (Different) or orientation-distractor (Same) displays; h_Glover_ is the Glover haemodynamic response function, and ∗ denotes convolution. T stacks the three regressors in the order Absent, Different, Same. GS is the global signal. These are display-event nuisance regressors, not regressors isolating a cognitive mechanism.

Ỹ_ir_^(v)^ = Clean(Y_ir_; Z_ir_^(v)^)

z_ir,jk_^(v)^ = arctanh[cor(Ỹ_ir,j_^(v)^, Ỹ_ir,k_^(v)^)]

F_i_^(v)^ = [1/(2|E_DMN_|)]∑_r=1_^2^∑_(j,k)∈EDMN_z_ir,jk_^(v)^

r_capture_^(v)^ = cor({F_i_^(v)^ − F̄_g(i)_^(v)^}, {C_i_ − C̄_g(i)_})

Y_ir_ is the run-r parcel time-series matrix for participant i. The index v identifies the nuisance-regression variant, and j and k identify the parcel pair. Clean denotes the implemented Nilearn signal-cleaning operation, including the common filtering, retained-volume mask, detrending and standardization, with the specified confound design; the confounds are processed consistently with the signal. Z_none_ omits nuisance regression, not the common cleaning steps. E_DMN_ contains all unique, non-self edges among the 91 Yeo-7 Default parcels. Connectivity was Fisher-transformed within run and then averaged equally across the two runs. Each variant’s pooled capture association was centered within SS and DS. The primary and four alternatives retained the existing Holm-5 family. Attenuation under task or global-signal regression is a preprocessing dependence, not evidence for mediation.

#### Supplementary Equation S11. Conditional trial-count matching

***Trial-count matching between biased and unbiased distractor dimensions***

For participant i, let B_i_ be the eligible biased-dimension trial-index set, n_Bi_ its size, and m_i_ the eligible unbiased-dimension trial count (0 < m_i_ ≤ n_Bi_). In each of B = 10,000 draws, a uniformly selected subset S_i_^(b)^ of m_i_ biased trials was sampled without replacement. The participant’s mean RT on all eligible absent trials, Ā_i_, remained fixed and was shared with the unbiased-capture score. C_Bi_ denotes the full eligible biased-capture score, and C_Bi_^(b)^ denotes the subset score for draw b:

S_i_^(b)^ ⊆ B_i_, |S_i_^(b)^| = m_i_

C_Bi_^(b)^ = (1/m_i_)∑_j∈Si_^(b)^RT_ij_ − Ā_i_

C_Bi_^matched^ = (1/B)∑_b=1_^B^C_Bi_^(b)^

E_subset_[C_Bi_^(b)^ | observed trials] = (1/n_Bi_)∑_j∈Bi_RT_ij_ − Ā_i_ = C_Bi_

The averaged matched score C_Bi_^matched^ was computed within each participant and then used in the group-centered connectivity association and the existing direct biased-versus-unbiased comparison. The expectation is over subset selection conditional on the observed participants and eligible trials, not over newly sampled participants. A random-subset mean is unbiased for the full eligible biased-trial mean; averaging many subsets therefore approaches the original full biased-capture score rather than independently equalizing measurement precision. The absent baseline is not resampled. Draw-wise distributions describe conditional trial-resampling stability, not participant-sampling confidence intervals or an independent replication. Existing correction families are unchanged.

##### *Equal-region participant capture scores*

*C*^B50^_i_ and *C*^U50^_i_ are the biased- and unbiased-dimension capture scores in milliseconds. *RT̄*_BF,i_, *RT̄*_BR,i_, *RT̄*_UF,i_ and *RT̄*_UR,i_ are participant *i*’s observed mean RTs in the four present conditions; *Ā*_i_ is the absent-trial mean. The superscripts ^B50^ and ^U50^ identify equal weighting of the frequent and rare regions within each dimension:

*C*^B50^_i_ = (*RT̄*_BF,i_ + *RT̄*_BR,i_) / (2) − *Ā*_i_

*C*^U50^_i_ = (*RT̄*_UF,i_ + *RT̄*_UR,i_) / (2) − *Ā*_i_

#### Supplementary Equation S12. Trial-history sensitivity: hierarchical model and participant-level OLS diagnostic

**A. Three-category coding and fixed-effect components**

This sensitivity is distinct from the principal five-condition capture model in S13. It collapses distractor-present trials into assigned-biased and assigned-unbiased categories and adjusts recorded history. For trial j from participant i, Y = RT/100. Indicators b and u equal 1 on biased and unbiased present trials, respectively, and both equal 0 on absent trials. D codes SS = 0 and DS = 1; X is group-centered DMN connectivity scaled by the pooled participant SD as defined in S4. The superscript hist identifies coefficients estimated jointly in the history-adjusted model. Its fixed absent-trial mean and present-minus-absent components are:

*μ*^hist^_i_ = *β*^hist^_0_ + *β*^hist^_X_*X*_i_ + *β*^hist^_D_*D*_i_ + *β*^hist^_XD_*X*_i_*D*_i_

*λ*^hist^_B,i_ = *β*^hist^_b_ + *β*^hist^_b,X_*X*_i_ + *β*^hist^_b,D_*D*_i_ + *β*^hist^_b,XD_*X*_i_*D*_i_

*λ*^hist^_U,i_ = *β*^hist^_u_ + *β*^hist^_u,X_*X*_i_ + *β*^hist^_u,D_*D*_i_ + *β*^hist^_u,XD_*X*_i_*D*_i_

These expressions expand the 12 core fixed coefficients of one jointly fitted model, not three separate regressions. The absent-trial X coefficient describes baseline RT–connectivity coupling, not a distractor-capture slope.

**B. Recorded history and complete hierarchical model**

R indicates run 2 (run 1 = 0; run 2 = 1). O is original global chronological trial order across runs. H_B and H_U count biased and unbiased exposures strictly before the current trial and accumulate across runs. Order and prior exposures are constructed from the complete recorded sequence before correctness/RT exclusions. For v equal to log(O), H_B or H_U, z_i(v) subtracts the participant’s mean and divides by the sample SD over that participant’s eligible rows:

*z*_i_(v) = (v − *v̄*_i_) / (*s*_v,i_)

*h*^hist^_ij_ = *β*^hist^_R,i_*R*_ij_ + *β*^hist^_order,i_*z*_i_[log(*O*_ij_)]

+ *β*^hist^_priorB,i_*z*_i_(*H*^B^_ij_) + *β*^hist^_priorU,i_*z*_i_(*H*^U^_ij_)

*Y*_ij_ = *μ*^hist^_i_ + *λ*^hist^_B,i_*b*_ij_ + *λ*^hist^_U,i_*u*_ij_ + *h*^hist^_ij_

+ *a*^hist^_0i_ + *a*^hist^_bi_*b*_ij_ + *a*^hist^_ui_*u*_ij_ + *ε*^hist^_ij_

(*a*^hist^_0i_, *a*^hist^_bi_, *a*^hist^_ui_)^T^ ∼ *N*_3_(0, *Σ*^hist^)

*ε*^hist^_ij_ ∼ N(0, *σ*^2^_hist_)

The four history slopes are participant-specific fixed adjustments: run, log order, prior biased exposure and prior unbiased exposure. They add 132 columns for 33 participants to the 12-column core. The three-dimensional Gaussian random vector contains a participant intercept and biased/unbiased condition slopes with covariance matrix Σ_hist; residuals have Gaussian variance σ²_hist. All core coefficients, history slopes, random-effects covariance and residual variance are estimated jointly rather than carrying over the unadjusted model’s coefficients. The logarithm is log(O), not log(1 + O). The fit retains all 10,995 eligible trials from 33 participants (SS = 16; DS = 17), REML estimation, Satterthwaite tests and pointwise t intervals. These adjustments do not identify spatial learning separately from exposure.

**C. History-adjusted DMN–capture slopes and contrasts**

*κ*_B,SS_ = 100*β*^hist^_b,X_

*κ*_B,DS_ = 100*[β*^hist^_b,X_ + *β*^hist^_b,XD_]

*κ*_B,pooled_ = (16) / (33)*κ*_B,SS_ + (17) / (33)*κ*_B,DS_

*κ*_U,SS_ = 100*β*^hist^_u,X_

*κ*_U,DS_ = 100*[β*^hist^_u,X_ + *β*^hist^_u,XD_]

*κ*_U,pooled_ = (16) / (33)*κ*_U,SS_ + (17) / (33)*κ*_U,DS_

*δ*_B−U_ = *κ*_B,pooled_ − *κ*_U,pooled_

κ denotes a present-minus-absent RT–connectivity slope. Multiplication by 100 converts RT/100 coefficients to milliseconds per pooled participant SD of group-centered DMN connectivity. The pooled contrasts weight SS by 16/33 and DS by 17/33. The biased group contrast is κ_B,DS − κ_B,SS, and the unbiased group contrast is κ_U,DS − κ_U,SS; δ_B−U is the pooled biased-minus-unbiased contrast. All are covariance-aware linear contrasts of the same history-adjusted fit, not comparisons of separate significance labels. Supplementary Table S5A retains Holm-6 across the six connectivity-related Type-III terms. Table S5B retains its original Holm-9 family across the pooled, SS and DS biased/unbiased slopes (six tests), the two DS-minus-SS moderation contrasts, and the pooled biased-minus-unbiased contrast. Neither family is reduced by consolidating the presentation.

**D. Separate participant-level OLS diagnostic**

*Y*_ij_ = *β*^OLS^_0,i_ + *β*^OLS^_b,i_*b*_ij_ + *β*^OLS^_u,i_*u*_ij_ + *h*^OLS^_ij_ + *e*^OLS^_ij_

*h*^OLS^_ij_ = *β*^OLS^_R,i_*R*_ij_ + *β*^OLS^_order,i_*z*_i_[log(*O*_ij_)]

+ *β*^OLS^_priorB,i_*z*_i_(*H*^B^_ij_) + *β*^OLS^_priorU,i_*z*_i_(*H*^U^_ij_)

Each participant was fitted with a separate seven-coefficient OLS regression: an intercept, biased and unbiased condition effects, and four history adjustments. The outcome, condition indicators, run, trial-order and prior-exposure variables, and within-participant standardization are defined in subsections A–B. The OLS history term uses the same design with independently estimated coefficients, without shared random effects or participant-level partial pooling. Condition coefficients and standard errors are multiplied by 100 for millisecond reporting.

**Supplementary Equation S13. Hierarchical trial-level capture model**

##### *Capture model: indices, predictors and components*

The model retains all 10,995 correct, RT-valid scanner trials from 33 participants (SS = 16, DS = 17). *i* indexes participants and *j* their trials. *Y*_ij_ is trial RT divided by 100; *X*_i_ is participant-level DMN connectivity centered within the group and divided by the pooled participant SD of the centered values (S4). *D*_i_ = 0 for SS and 1 for DS; *R*_ij_ = 0 in run 1 and 1 in run 2. Connectivity is averaged across the two runs and is constant across a participant’s trials.

The reference condition A is distractor absent. The four present conditions are biased-frequent (BF), biased-rare (BR), unbiased-frequent (UF) and unbiased-rare (UR). *K* = {BF, BR, UF, UR} contains the present conditions, and *C* = {A} ∪ *K* contains all five. *c*_kij_ equals 1 when trial *j* of participant *i* belongs to condition *k*, and 0 otherwise. Frequent/rare on unbiased trials identifies the same assigned regions, not a spatial contingency in that dimension.

The superscript ^mix^ labels this fitted capture model; it is not an exponent. *μ*^mix^_i_ is the fixed-effect absent-trial mean in run 1. *λ*^mix^_ki_ is condition *k*’s fixed-effect present-minus-absent increment in run 1. *ζ*^mix^_ij_ adds the condition-specific run-2 shift, including the absent condition; *β*^mix^_R:k_ is that shift for condition *k*. In *μ*^mix^_i_, *β*^mix^_0_ is the reference intercept, while *β*^mix^_X_, *β*^mix^_D_ and *β*^mix^_XD_ identify the connectivity, group and connectivity-by-group terms. In *λ*^mix^_ki_, *β*^mix^_k_, *β*^mix^_k,X_, *β*^mix^_k,D_ and *β*^mix^_k,XD_ are the corresponding condition-specific increment and interactions. These components algebraically expand one model, not separate fits.

*a*^mix^_0i_ is participant *i*’s random intercept; *a*^mix^_ki_ is the random present-minus-absent slope for condition *k*. Their five-dimensional vector has mean zero and covariance *Σ*^mix^; *N*_5_ denotes its multivariate normal distribution, and the superscript ^T^ denotes transpose. *ε*^mix^_ij_ is trial-level Gaussian residual error with variance *σ*^2^_mix_, distinct from participant-level variation.

*μ*^mix^_i_ = *β*^mix^_0_ + *β*^mix^_X_*X*_i_ + *β*^mix^_D_*D*_i_ + *β*^mix^_XD_*X*_i_*D*_i_

*λ*^mix^_ki_ = *β*^mix^_k_ + *β*^mix^_k,X_*X*_i_ + *β*^mix^_k,D_*D*_i_ + *β*^mix^_k,XD_*X*_i_*D*_i_, k ∈ K

*ζ*^mix^_ij_ = *R*_ij_*∑*_k∈C_*c*_kij_*β*^mix^_R:k_

*Y*_ij_ = *μ*^mix^_i_ + *∑*_k∈K_*c*_kij_*λ*^mix^_ki_ + *ζ*^mix^_ij_

+ *a*^mix^_0i_ + *∑*_k∈K_*c*_kij_*a*^mix^_ki_ + *ε*^mix^_ij_

(*a*^mix^_0i_, *a*^mix^_BF,i_, *a*^mix^_BR,i_, *a*^mix^_UF,i_, *a*^mix^_UR,i_)^T^ ∼ *N*_5_(0, *Σ*^mix^)

*ε*^mix^_ij_ ∼ N(0, *σ*^2^_mix_)

##### *Equal-region model-based connectivity–capture coupling*

*κ*^50^_B,d_ = 50*[β*^mix^_BF,X_ + d*β*^mix^_BF,XD_ + *β*^mix^_BR,X_ + d*β*^mix^_BR,XD_]

*κ*^50^_U,d_ = 50*[β*^mix^_UF,X_ + d*β*^mix^_UF,XD_ + *β*^mix^_UR,X_ + d*β*^mix^_UR,XD_]

*κ*^50^_B,d_ and *κ*^50^_U,d_ are model-based biased- and unbiased-dimension capture slopes: estimated changes in present-minus-absent RT cost per 1-SD higher DMN connectivity, in ms/DMN SD. The subscript *d* is the group value (0 = SS; 1 = DS), and the superscript ^50^ denotes equally weighted regions. The factor 50 combines the 0.5 region weight and multiplication by 100 to return RT/100 to milliseconds. Pooled slopes weight SS and DS by 16/33 and 17/33.

All fixed components and correlated random effects belong to one REML model, with Satterthwaite tests and pointwise t intervals. Biased slopes, unbiased slopes and their direct differences in pooled, SS and DS samples form Holm-9; the three DS-minus-SS slope contrasts form a separate Holm-3 family (Table 4A; Supplementary Table S6). The participant-score companion instead uses six connectivity correlations (Holm-6), three direct dependent-correlation comparisons (Holm-3) and 10,000 paired, group-stratified percentile bootstrap intervals (Table S3C). Equal-region weighting standardizes the evaluated mixture; it does not equalize trial information, exposure, reliability or physical dimension.

##### *Physical dimension versus assigned bias: same-physical-dimension contrasts*

For each present condition *k*, m_k_(d) is the fixed-effect present-minus-absent RT cost, in milliseconds, evaluated at group-mean connectivity (*X*_i_ = 0) in group *d* and averaged equally across runs. The run-offset difference subtracts the absent-condition shift. m_B,d_ and m_U,d_ then average the two region-specific costs within the biased and unbiased dimensions:

m_k_(d) = 100[β^mix^_k_ + dβ^mix^_k,D_ + 0.5(β^mix^_R:k_ − β^mix^_R:A_)]

m_B,d_ = 0.5[m_BF_(d) + m_BR_(d)], m_U,d_ = 0.5[m_UF_(d) + m_UR_(d)]

Δm_ori_ = m_B,0_ − m_U,1_, Δκ_ori_ = κ^50^_B,0_ − κ^50^_U,1_

Δm_col_ = m_B,1_ − m_U,0_, Δκ_col_ = κ^50^_B,1_ − κ^50^_U,0_

Δm_ori_ and Δm_col_ are between-group differences in orientation and color capture costs (ms); Δκ_ori_ and Δκ_col_ are the corresponding differences in connectivity–capture slopes (ms/DMN SD). Each contrast holds the physical dimension constant and places its assigned-biased group first: orientation is SS minus DS, whereas color is DS minus SS.

One omnibus test jointly tests the orientation and color mean-cost contrasts against zero (2 numerator degrees of freedom); a second jointly tests the two coupling contrasts. Separate one-degree-of-freedom tests assess each physical-dimension cost and coupling contrast.

**Supplementary Equation S14. Hierarchical trial-level spatial-learning model**

The hierarchical model uses 6,721 correct, RT-valid biased-dimension scanner trials from the same 33 imaging participants. *i* indexes participants and *j* trials. *Y*_ij_, *X*_i_ and *D*_i_ retain the outcome scaling, participant-level connectivity scaling and group coding defined in S13. *L*_ij_ = 0 for frequent and 1 for rare locations; *R*_ij_ = 0 in run 1 and 1 in run 2. The participant score and mixed-model coupling below are distinct estimands.

##### *Participant-level spatial-learning score*

*S*_i_ is a participant *i*’s rare-minus-frequent RT difference in milliseconds. *RT̄*_Rare,i_ and *RT̄*_Frequent,i_ are observed mean RTs on biased-dimension trials, pooling the two scanner runs:

*S*_i_ = *RT̄*_Rare,i_ − *RT̄*_Frequent,i_

##### *Run-moderated hierarchical model and run-specific coupling*

The superscript ^run^ labels the run-moderated learning model, not an exponent. μ^run^_i_ is the fixed-effect frequent-location mean in run 1; θ^run^_i_ is the fixed-effect rare-minus-frequent increment in run 1. Their displayed expansions include connectivity, group and connectivity-by-group effects. *β*^run^_R_ shifts frequent-location RT in run 2; *β*^run^_LR_ changes the rare-minus-frequent increment. *β*^run^_XR_ changes frequent-location connectivity coupling, whereas *β*^run^_LXR_ changes connectivity coupling of the rare-minus-frequent increment.

*a*^run^_0i_ is participant *i*’s random intercept and *a*^run^_Li_ the random rare-minus-frequent slope. They are jointly Gaussian with mean zero and an estimated covariance matrix; *ε*^run^_ij_ is Gaussian trial-level residual error with a separately estimated variance. These random effects and coefficients belong to the learning fit, not the capture fit.

*κ*^run^_d,r_ is the model-based rare-minus-frequent connectivity slope (ms/DMN SD) for group *d* and run *r*. Here lowercase *d* = 0/1 selects SS/DS, and lowercase *r* = 0/1 selects runs 1/2; *D*_i_ and *R*_ij_ are the corresponding observed predictors. The lowercase *r* in *κ*^run^_d,r_ is the numeric run index, not a correlation coefficient; *r*_F,ori_, *r*_F,col_ and *r*_FB_ denote correlations.

μ^run^_i_ = β^run^_0_ + β^run^_X_X_i_ + β^run^_D_D_i_ + β^run^_XD_X_i_D_i_

θ^run^_i_ = β^run^_L_ + β^run^_L,X_X_i_ + β^run^_L,D_D_i_ + β^run^_L,XD_X_i_D_i_

*Y*_ij_ = *μ*^run^_i_ + *θ*^run^_i_*L*_ij_ + *β*^run^_R_*R*_ij_ + *β*^run^_LR_*L*_ij_*R*_ij_

+ *β*^run^_XR_*X*_i_*R*_ij_ + *β*^run^_LXR_*L*_ij_*X*_i_*R*_ij_ + *a*^run^_0i_ + *a*^run^_Li_*L*_ij_ + *ε*^run^_ij_

*κ*^run^_d,r_ = 100*[β*^run^_L,X_ + d*β*^run^_L,XD_ + r*β*^run^_LXR_]

##### *Equal-run-averaged learning coupling*

κ̄^run^_d_ is the equally weighted average of the two run-specific slopes for group *d*; each run receives weight 0.5 (Table 4C):

κ̄^run^_d_ = 100[β^run^_L,X_ + dβ^run^_L,XD_ + 0.5β^run^_LXR_]

κ̄^run^_0_ is the SS slope and κ̄^run^_1_ the DS slope. The pooled slope is (16/33)κ̄^run^_0_ + (17/33)κ̄^run^_1_; the direct DS-minus-SS contrast is κ̄^run^_1_ − κ̄^run^_0_. These are contrasts of the same fitted model. The three pooled/SS/DS averaged slopes plus the direct group contrast form the four-member Holm-4 family (Table 4C; Supplementary Table S7).

##### *Run-related change in coupling*

*Δκ*_run2−run1_ = 100*β*^run^_LXR_

*Δκ*_run2−run1_ is the run-2-minus-run-1 change in learning coupling, in ms/DMN SD. It is determined by *β*^run^_LXR_ and is common to SS and DS: the model includes no group-by-location-by-connectivity-by-run interaction. *β*^run^_XR_ retains the lower-order connectivity-by-run term but does not itself represent a change in rare-minus-frequent coupling. Connectivity remains participant-level and averaged across runs; this is not a test of run-specific changes in connectivity.

The learning model uses REML estimation, Satterthwaite tests and pointwise t intervals, conditional on the retained correlated random-intercept/location-slope structure.

#### Supplementary Methods S15. Network-level associations and edge enrichment

##### *Network summaries and behavioral outcomes*

Analyses used the 33-participant imaging sample (SS = 16, DS = 17). Seven within-network and 21 between-network Yeo-7 summaries yielded 28 network-pair measures. Each participant’s summary was the mean Fisher-z connectivity across the constituent unique edges, with the two scanner runs weighted equally. Biased-dimension capture was distractor-present RT minus distractor-absent RT under the observed trial mixture; spatial learning was rare-minus-frequent RT on biased-dimension distractor trials. The biased dimension was orientation in SS and color in DS.

##### *Network-summary associations and direct comparisons*

Pooled analyses centered connectivity and behavior within SS and DS (N = 33, association df = 30); subgroup analyses used ordinary Pearson correlations (SS: N = 16, df = 14; DS: N = 17, df = 15). For each outcome, six approximate Williams tests compared the pooled within-DMN association directly with the other within-network associations, using effective N = 32, df = 29 and Holm-6. These comparisons tested differences in association strength, not unique conditional variance or causation.

*Permutation network-summary sensitivity.* All 168 associations were additionally evaluated with 10,000 synchronized behavioral-label permutations restricted within SS and DS. The same participant rearrangement was used across network summaries and applied consistently across the sample panels and outcomes, retaining their dependence. Connectivity values and group membership were held fixed; pooled connectivity and behavioral variables were centered separately within SS and DS. For each association, the unadjusted permutation p value compared the magnitude of the observed statistic with its own two-sided permutation null distribution. Holm-28 adjusted these individual permutation p values within each 28-pair outcome/sample panel: sorted p values were multiplied by the successively decreasing numbers of remaining hypotheses, with cumulative maxima and an upper bound of 1. This sequential procedure does not use the empirical dependence among the test statistics. In contrast, maxT-28 compared each observed statistic with the maximum absolute association statistic across all 28 pairs in that panel on each synchronized permutation. Thus, maxT uses the joint permutation distribution and incorporates dependence between tests; it is a separate family-wise adjustment.

*Matched seven-network extension.* An exploratory extension evaluated equal-region biased capture and spatial learning against the seven within-network summaries in pooled, SS and DS samples. Equal-region capture was the equally weighted mean of frequent- and rare-region biased RTs minus absent RT; it was distinct from the original capture score. Associations across two outcomes, three samples and seven networks formed Holm-42; direct DMN-versus-network comparisons across two outcomes, three samples and six comparators formed Holm-36. Analytic association and approximate Williams tests were primary. Shared 20,000 paired participant-bootstrap draws, stratified by group, provided percentile intervals and approximate centered-bootstrap comparison tests; 10,000 within-group outcome permutations assessed association sensitivity. Pooled variables were recentered within groups in each bootstrap draw.

Group moderation was the SS-minus-DS difference in each DMN-versus-network association contrast. Approximate bootstrap-Wald tests jointly assessed the six contrasts for each outcome (Holm-2 across outcomes), using their shared bootstrap covariance; individual moderation contrasts retained Holm-12.

***Descriptive signed-edge screens***

Both distractor capture and spatial learning outcomes used the same 399 common-coverage parcels and 79,401 unique undirected non-self edges. Connectivity and behavior were centered within SS and DS before pooled edgewise association (N = 33, df = 30). Only pooled edge-count screens were reported. Edges meeting exploratory raw two-sided *p* < .01 were separated by association sign and counted across 28 mutually exclusive within/between-network categories. Seven additional endpoint categories counted edges touching each network; these categories overlapped and their counts must not be summed. Screened edges were descriptive selections, not multiplicity-corrected discoveries.

##### *Capacity-adjusted enrichment and multiplicity*

*Whole-connectome base rates.* The reference universe comprised all unique, undirected connections between the 399 common-coverage parcels, excluding self-connections and duplicate edges: M = 399 × 398 / 2 = 79,401 possible edges. The seven Yeo networks contained 91 Default, 77 Somatomotor, 61 Visual, 52 Control, 47 Salience/Ventral Attention, 46 Dorsal Attention, and 25 Limbic parcels. For a network a with nₐ parcels, the within-network capacity was mₐₐ = nₐ(nₐ − 1)/2; between two different networks a and b, it was mₐᵦ = nₐnᵦ. The 28 within- and between-network categories partitioned the complete edge universe. For the additional endpoint category of edges touching network a (at least one endpoint in that network), the capacity was mₐ,touch = nₐ(nₐ − 1)/2 + nₐ(399 − nₐ). A connection was counted once within a touching category, although between-network edges appeared in both endpoint networks’ touching categories. For any category c, the whole-connectome base rate was f꜀ = m꜀/M. For example, the 4,095 possible within-Default edges represented 5.157% of the universe, whereas the 32,123 edges touching Default represented 40.457%. These base rates reflect category capacity, not the screening threshold or the distribution of selected edges.

*Expected counts and descriptive ratios.* Distractor capture and spatial learning were screened separately using pooled, group-centered edgewise associations and an exploratory raw two-sided p < .01 threshold. Selected edges were separated into positive and negative associations. For each sign s, let Kₛ be the total number of selected edges across the full connectome and O꜀,ₛ the number selected in category c. The capacity-based expected count was E꜀,ₛ = Kₛ f꜀, and the observed/expected ratio was R꜀,ₛ = O꜀,ₛ/E꜀,ₛ. Thus, a ratio above one indicates greater descriptive concentration than expected from category size alone; it does not establish statistically significant enrichment.

*Permutation inference and multiplicity.* Formal enrichment was assessed using the observed-minus-expected surplus D꜀,ₛ = O꜀,ₛ − Kₛf꜀. For each behavioral outcome, 10,000 permutations shuffled participant behavioral labels separately within SS and DS and repeated the complete signed edgewise screen. Both the total selected count Kₛ and category count O꜀,ₛ were recalculated in every permutation, while the category base rate f꜀ remained fixed. The upper-tail permutation p value was [1 + number of permuted surpluses ≥ the observed surplus]/10,001. Holm correction was applied separately to the 28 mutually exclusive network-pair categories across two signs (56 tests) and the seven overlapping network-endpoint categories across two signs (14 tests), with separate families for capture and spatial learning. Permuting participant labels preserved the dependence structure among connectivity edges. These enrichment tests were distinct from descriptive counts and ratios, network-summary associations, and full-universe Holm-79,401 edgewise tests (Tables S9 and S10).

*Exploratory maximum-statistic sensitivity.* In addition to the retained marginal upper-surplus permutation p values and their Holm adjustments, maxZ standardized each category/sign surplus using the mean and standard deviation across the observed row plus the 10,000 null rows. Each observed standardized surplus was compared with the maximum standardized surplus across its declared family in each permutation, using the same add-one p-value rule. Holm sequentially adjusts individual permutation p values; maxZ instead uses their joint standardized-statistic distribution, retaining dependence among categories and signs. Families remain outcome-specific: 56 pair/sign tests and 14 overlapping touching/sign tests per outcome. The 140 reported tests do not form a single corrected family; capture and learning used different permutation seeds, and no combined cross-outcome maximum was calculated.

#### Supplementary Methods S16. Exploratory connectome-based prediction

##### *Targets, sample and predefined edge sets*

Exploratory connectome-based prediction assessed whether connection patterns within and between DMN subdivisions could estimate attentional capture or spatial learning in participants held out from model fitting. These analyses assessed same-sample held-out performance in the imaging sample (N = 33; SS = 16, DS = 17), not independent external validation. Capture and spatial learning were modeled separately. The original capture target was biased-dimension distractor-present RT minus absent RT under the observed trial mixture; the learning target was rare-minus-frequent RT on biased-dimension trials.

The physical Yeo-17 DefaultA/B/C set contained 79 parcels (DefaultA = 34, DefaultB = 32, DefaultC = 13), including 70 parcels overlapping the focal Yeo-7 Default91 set. It complemented, rather than partitioned or replaced, Default91. Six predefined edge sets were evaluated: A–A, A–B, A–C, B–B, B–C and C–C, with 561, 1,088, 442, 496, 416 and 78 candidate edges, respectively.

##### *Training-only feature selection and prediction*

Each leave-one-participant-out fold held out participant i and used the remaining 32 participants for training. Here, j indexes a training participant, c indexes a predefined edge set, and g(j) denotes group. V is the behavioral target; the superscript c on V denotes a centered target, whereas the subscript c on feature sets and coefficients identifies the edge set. Z is raw Fisher-z edge connectivity, not a group-centered edge score. The training outcome was centered using its training-group mean. Positive and negative feature sets E⁺ and E⁻ were selected by ordinary two-sided Pearson tests within training data only (*p* < .01; training df = 30). Signed network strength was the sum of positive-edge connectivity minus the sum of negative-edge connectivity. Ordinary least squares related this strength to the centered training outcome:

*V*^c^_j,−i_ = *V*_j_ − *V̄*_g(j),train(−i)_

*S*_jic_ = *∑*_e∈E⁺ᵢc_*Z*_je_ − *∑*_e∈E⁻ᵢc_*Z*_je_

*V*^c^_j,−i_ = *β*^CPM^_0,ic_ + *β*^CPM^_S,ic_*S*_jic_ + *η*_jic_

*V̂*^c^_i,−i_ = *β̂*^CPM^_0,ic_ + *β̂*^CPM^_S,ic_*S*_iic_

The fitted intercept and strength coefficient were applied to the held-out participant’s training-selected edges. The observed held-out target was centered using the corresponding training-group mean, never a mean recomputed using the held-out participant. Performance was the Pearson correlation between the fold-specific centered targets and their held-out predictions:

V^c^_i,−i_ = V_i_ − V̄_g(i),train(−i)_, r_CV,c_ = cor({V^c^_i,−i_}, {V̂^c^_i,−i_})

All centering, feature selection, strength construction, regression fitting and prediction were repeated within every held-out fold. Zero variance in training strength yielded a centered prediction of zero. Null or negative models and zero-strength folds were retained. A constant prediction vector does not define a Pearson correlation; a zero fallback must not be interpreted as an estimable correlation. Capture and learning remained separate targets.

##### *Permutation inference and correction families*

Inference used 10,000 outcome-label permutations within SS and DS, each repeating the complete training-only prediction pipeline. The primary two-sided equal-tail p value was twice the smaller add-one upper- or lower-tail probability, capped at one. The absolute-tail sensitivity counted null correlations whose absolute magnitude equaled or exceeded the observed magnitude, also using the add-one rule. Separate outcome-specific Holm-6 families covered the six DefaultA/B/C edge sets under each tail rule.

### Supplementary Results

#### Exploratory connectome-based prediction

Exploratory connectome-based prediction assessed whether connection patterns within and between DMN subdivisions could estimate attentional capture or spatial learning in participants held out from model fitting. All prediction analyses used the same participants and nested feature selection, centering and regression within training folds and 10,000 within-group permutations. Equal-tail and absolute-tail tests retain separate correction families. The physical DefaultA/B/C set is complementary to, not a partition of, the focal Default91set.

*DefaultA/B/C prediction of attentional capture*

Only within-DefaultB survived the primary equal-tail Holm-6 family, not the absolute-tail Holm-6 sensitivity (**Figure S1**).

DefaultA (A–A): *r* = .459; equal-tail *p* = .0408 / .1690; absolute-tail *p* = .0821 / .3855.

A–B: *r* = .458; equal-tail *p* = .0338 / .1690; absolute-tail *p* = .0771 / .3855.

A–C: *r* = .098; equal-tail *p* = .6651 / 1.0000; absolute-tail *p* = .7740 / 1.0000.

DefaultB (B–B): *r* = .619; equal-tail *p* = .0012 / .0072; absolute-tail *p* = .0130 / .0780.

B–C: *r* = .194; equal-tail *p* = .4448 / 1.0000; absolute-tail *p* = .5919 / 1.0000.

DefaultC (C–C): *r* = .044; equal-tail *p* = .3972 / 1.0000; absolute-tail *p* = .6583 / 1.0000.

All observed selected edges were positive. Median positive selections and ranges across the 33 folds were: DefaultA 38 (25–48); A–B 81 (57–100); A–C 12 (6–22); DefaultB 49 (33–61); B–C 10 (3–16); DefaultC 2 (0–8).

*DefaultA/B/C prediction of spatial learning*

The largest positive held-out correlations were A–B and within-DefaultB; neither supported corrected prediction.

DefaultA (A–A): *r* = .047; equal-tail *p* = .8583 / 1.0000; absolute-tail *p* = .9027 / 1.0000.

A–B: *r* = .364; equal-tail *p* = .1686 / 1.0000; absolute-tail *p* = .2280 / 1.0000.

A–C: *r* = −.182; equal-tail *p* = .7715 / 1.0000; absolute-tail *p* = .6460 / 1.0000.

DefaultB (B–B): *r* = .358; equal-tail *p* = .1812 / 1.0000; absolute-tail *p* = .2855 / 1.0000.

B–C: *r* = −.017; equal-tail *p* = .8895 / 1.0000; absolute-tail *p* = .9523 / 1.0000.

DefaultC (C–C): constant zero predictions in all 33 folds; equal-tail *p* = .9881 / 1.0000; absolute-tail *p* = 1.0000 / 1.0000.

All observed selected edges were positive. Median positive selections and fold ranges were: DefaultA 16 (7–30); A–B 49 (20–68); A–C 3 (1–12); DefaultB 23 (11–33); B–C 8 (3–19); DefaultC 0 (0–0).

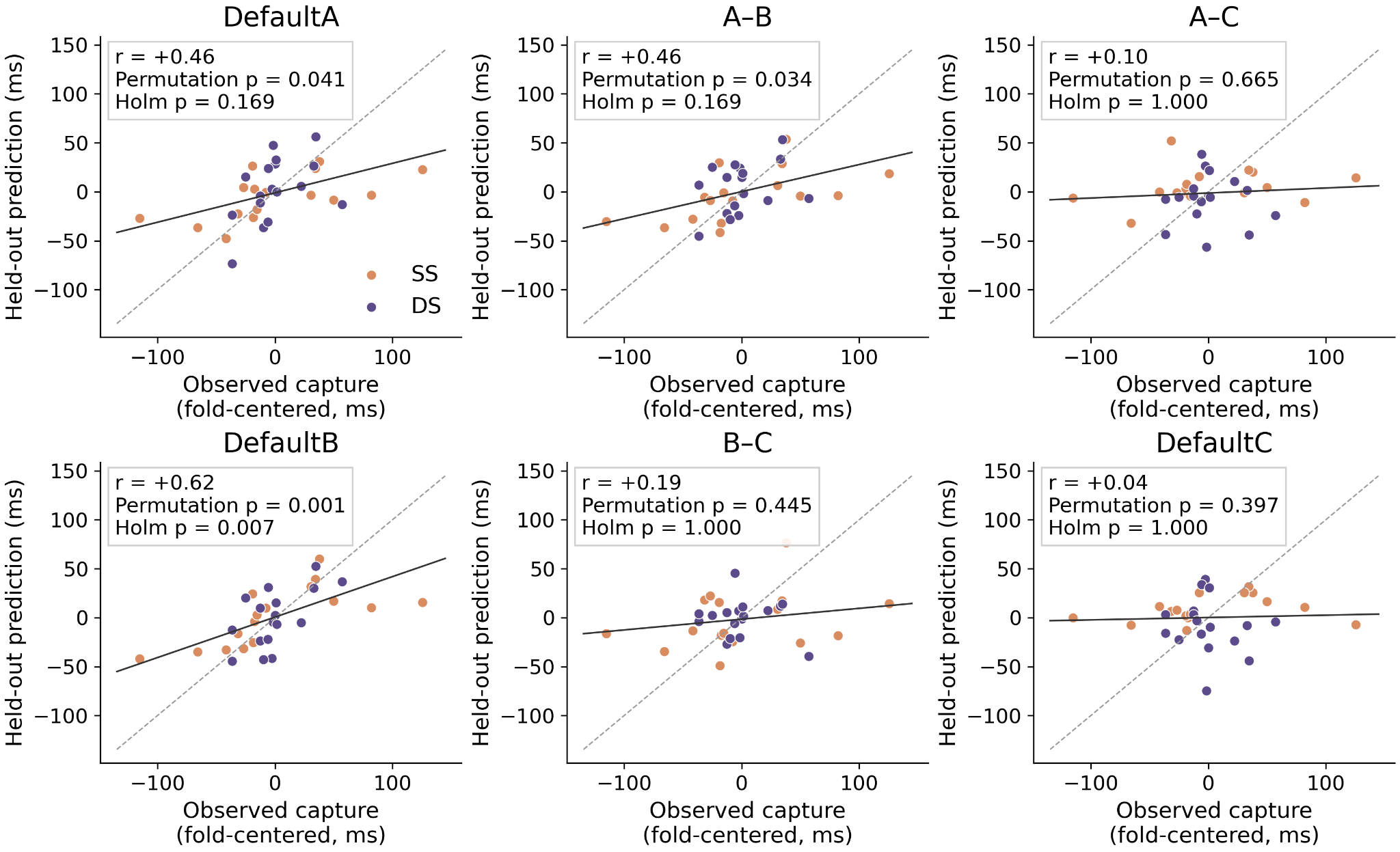

**Figure S1.** Exploratory connectome-based prediction for all six physical Yeo-17 DefaultA/B/C edge families. Each dot pairs one participant’s observed target centered using that fold’s training-group mean with its held-out prediction. Edge screening, signed-strength calculation, centering, and regression were repeated inside every training fold and each of 10,000 within-group permutations. Panel r values are held-out prediction correlations; displayed p_Holm values use the declared equal-tail permutation rule and Holm-6. The absolute-correlation-tail sensitivity is reported in the accompanying capture-prediction results and gives a different conclusion for DefaultB. The 79 physical parcels comprise DefaultA=34, DefaultB=32, and DefaultC=13; this analysis is complementary to, not a partition of, the focal Default91 set. N=33.

**Supplementary tables**

**Supplementary Table S1. Behavioral learning and capture tests**

| **Contrast** | **N** | **Mean / model estimate (ms) [95% CI]** | **t (df)** | **Raw p** | **Holm p** | **Standardized effect [95% CI]** |
| --- | --- | --- | --- | --- | --- | --- |
| LMM: SS rare minus frequent | 35 | 44.16 [29.13, 59.19] | 5.76 (11402.11) | <.001 | — | Raw ms contrast |
| LMM: DS rare minus frequent | 35 | 8.53 [−5.84, 22.89] | 1.16 (11402.04) | 0.2447 | — | Raw ms contrast |
| LMM: DS − SS rare-minus-frequent RT difference | 35 | −35.63 [−56.42, −14.84] | −3.36 (11402.08) | <.001 | — | Raw ms contrast |
| DS learning | 18 | 10.27 [−6.56, 27.11] | 1.29 (17.00) | 0.2153 | 0.2153 | $d_{z}$ 0.30 [−0.16, 0.79] |
| SS learning | 17 | 44.93 [14.47, 75.38] | 3.13 (16.00) | 0.0065 | 0.0130 | $d_{z}$ 0.76 [0.38, 1.35] |
| DS: All present weighted | 18 | 51.35 [36.54, 66.16] | 7.32 (17.00) | <.001 | <.001 | $d_{z}$ 1.72 [1.31, 2.72] |
| DS: Orientation | 18 | 191.23 [158.00, 224.46] | 12.14 (17.00) | <.001 | <.001 | $d_{z}$ 2.86 [2.30, 4.18] |
| DS: Color | 18 | 6.91 [−4.41, 18.24] | 1.29 (17.00) | 0.2152 | 0.4304 | $d_{z}$ 0.30 [−0.16, 0.74] |
| SS: All present weighted | 17 | 98.70 [76.82, 120.58] | 9.56 (16.00) | <.001 | <.001 | $d_{z}$ 2.32 [1.57, 4.16] |
| SS: Orientation | 17 | 132.25 [104.91, 159.59] | 10.26 (16.00) | <.001 | <.001 | $d_{z}$ 2.49 [1.81, 4.18] |
| SS: Color | 17 | 8.73 [−5.41, 22.86] | 1.31 (16.00) | 0.2092 | 0.4184 | $d_{z}$ 0.32 [−0.15, 0.91] |
| DS−SS: All present weighted | 18/17 | −47.35 [−72.90, −21.80] | −3.79 (28.49) | <.001 | 0.0021 | g −1.27 [−2.30, −0.59] |
| DS−SS: Orientation | 18/17 | 58.98 [17.52, 100.44] | 2.90 (32.10) | 0.0067 | 0.0134 | g 0.95 [0.35, 1.74] |
| DS−SS: Color | 18/17 | −1.81 [−19.27, 15.64] | −0.21 (31.15) | 0.8335 | 0.8335 | g −0.07 [−0.80, 0.57] |
| DS−SS: Biased dimension | 18/17 | −125.34 [−154.35, −96.32] | −8.97 (21.42) | <.001 | <.001 | g −3.03 [−4.68, −2.23] |
| DS−SS: Unbiased dimension | 18/17 | 182.51 [147.11, 217.90] | 10.67 (22.86) | <.001 | <.001 | g 3.45 [2.79, 4.86] |

*Note.* RT: ms; CIs: pointwise 95%. Paired $d_{z}$ is for within-participant effects; LMM contrasts have no $d_{z}$. Between-group effects use Hedges $g$. Effect-size CIs: 10,000 participant-percentile bootstraps. Holm families: two learning effects, five capture definitions/group, five between-group capture comparisons; duplicate scores retain family membership. LMM contrasts are unadjusted.

**Supplementary Table S2. Reliability complements: sample, pooling and connectivity diagnostics**

**A. Run-to-run physical capture: full behavioral sample**

| **sample/group** | **Dimension / bias** | **N** | **r [95% bootstrap CI]** | **t (df)** | **Raw p** | **Holm-4 p** |
| --- | --- | --- | --- | --- | --- | --- |
| Full sample / SS | Orientation (biased) | 17 | 0.626 [0.224, 0.849] | 3.11 (15) | 0.0072 | 0.0287 |
| Full sample / SS | Color (unbiased) | 17 | 0.054 [−0.418, 0.449] | 0.21 (15) | 0.8379 | 1.0000 |
| Full sample / DS | Orientation (unbiased) | 18 | 0.499 [0.177, 0.774] | 2.30 (16) | 0.0351 | 0.1053 |
| Full sample / DS | Color (biased) | 18 | 0.078 [−0.491, 0.720] | 0.31 (16) | 0.7570 | 1.0000 |

**B. Run-to-run pooled capture: uncentered and full-sample complements**

| **sample / score** | **Pooling** | **N** | **r [95% bootstrap CI]** | **t (df)** | **Raw p** | **Holm-2 p** | **Spearman–Brown projection [95% bootstrap CI]** |
| --- | --- | --- | --- | --- | --- | --- | --- |
| Imaging sample / biased dimension | Uncentered | 33 | 0.807 [0.701, 0.904] | 7.62 (31) | <.001 | <.001 | 0.893 [0.824, 0.950] |
| Imaging sample / unbiased dimension | Uncentered | 33 | 0.819 [0.709, 0.907] | 7.96 (31) | <.001 | <.001 | 0.901 [0.830, 0.951] |
| Full sample / biased dimension | Uncentered | 35 | 0.818 [0.718, 0.909] | 8.16 (33) | <.001 | <.001 | 0.900 [0.836, 0.952] |
| Full sample / biased dimension | Group centered | 35 | 0.507 [0.138, 0.754] | 3.33 (32) | 0.0022 | 0.0044 | 0.673 [0.242, 0.859] |
| Full sample / unbiased dimension | Uncentered | 35 | 0.825 [0.711, 0.910] | 8.38 (33) | <.001 | <.001 | 0.904 [0.831, 0.953] |
| Full sample / unbiased dimension | Group centered | 35 | 0.413 [0.111, 0.663] | 2.57 (32) | 0.0152 | 0.0152 | 0.585 [0.200, 0.797] |

**C. Run-to-run DMN connectivity**

| **Pooling** | **N** | **r [95% bootstrap CI]** | **t (df)** | **Raw p** | **Spearman–Brown projection [95% bootstrap CI]** |
| --- | --- | --- | --- | --- | --- |
| Uncentered | 33 | 0.718 [0.474, 0.881] | 5.74 (31) | <.001 | 0.836 [0.644, 0.937] |
| Group centered | 33 | 0.733 [0.510, 0.880] | 5.91 (30) | <.001 | 0.846 [0.675, 0.936] |

**D. Odd–even capture: uncentered and pooled physical-color diagnostics**

| **Score** | **Pooling** | **N** | **r [95% bootstrap CI]** | **t (df)** | **Raw p** | **Holm-2 p** | **Spearman–Brown projection [95% bootstrap CI]** |
| --- | --- | --- | --- | --- | --- | --- | --- |
| Biased-dimension capture | Uncentered | 33 | 0.699 [0.488, 0.861] | 5.45 (31) | <.001 | <.001 | 0.823 [0.656, 0.925] |
| Unbiased-dimension capture | Uncentered | 33 | 0.776 [0.614, 0.895] | 6.84 (31) | <.001 | <.001 | 0.874 [0.761, 0.944] |
| Color-dimension capture | Uncentered | 33 | −0.268 [−0.557, 0.096] | -1.55 (31) | 0.1310 | Descriptive | −0.734 [−2.516, 0.175] |
| Color-dimension capture | Group centered | 33 | −0.229 [−0.537, 0.153] | -1.29 (30) | 0.2081 | Descriptive | −0.593 [−2.320, 0.266] |

**E. Odd–even physical capture: full behavioral sample**

| **sample/group** | **Dimension / bias** | **N** | **r [95% bootstrap CI]** | **t (df)** | **Raw p** | **Holm-4 p** | **Spearman–Brown projection [95% bootstrap CI]** |
| --- | --- | --- | --- | --- | --- | --- | --- |
| Full sample / SS | Orientation (biased) | 17 | 0.564 [0.127, 0.813] | 2.65 (15) | 0.0183 | 0.0733 | 0.721 [0.225, 0.897] |
| Full sample / SS | Color (unbiased) | 17 | 0.044 [−0.437, 0.411] | 0.17 (15) | 0.8666 | 0.8666 | 0.084 [−1.556, 0.583] |
| Full sample / DS | Orientation (unbiased) | 18 | 0.355 [−0.152, 0.736] | 1.52 (16) | 0.1487 | 0.2975 | 0.524 [−0.357, 0.848] |
| Full sample / DS | Color (biased) | 18 | −0.403 [−0.725, 0.105] | -1.76 (16) | 0.0972 | 0.2916 | −1.351 [−5.270, 0.190] |

*Note.* N = 35 overlaps the N = 33 imaging sample. Each occasion uses its own absent baseline; parity precedes exclusions. CIs are pointwise 95% paired participant-percentile intervals (10,000 draws; pooled draws stratified by group). Run and odd–even families are separate: Holm-4 for physical-dimension scores and Holm-2 for biased/unbiased scores within each sample/pooling definition. DMN and pooled physical-color rows are descriptive unadjusted diagnostics. df = N−2 within-group/uncentered and N−3 group-adjusted. Spearman–Brown is a comparable-two-part projection, not absolute agreement or validated trait reliability; negative projections do not imply negative reliability.

**Supplementary Table S3. Additional participant-level DMN associations and sensitivities**

| **Analysis / contrast** | **Group (N)** | **Estimate [95% CI]** | **Test (df)** | **Raw p** | **Holm p** | **Correction** |
| --- | --- | --- | --- | --- | --- | --- |
| **A. Complementary capture associations and robustness** | | | | | | |
| Biased-dimension capture | SS (16) | r = .635 Fisher [.203, .860] | t(14.00) = 3.08 | .0082 | .0493* | Holm-8 |
| Biased-dimension capture | DS (17) | r = .417 Fisher [−.080, .748] | t(15.00) = 1.78 | .0959 | .1918 | Holm-8 |
| All-present capture | Pooled (33) | r = .445 Fisher [.114, .687]  Bootstrap [.197, .641] | t(30.00) = 2.72 | .0106 | .0532 | Holm-8 |
| Biased-dimension capture: adjust unbiased + group | Pooled (33) | partial r = .577 Fisher [.280, .773] | t(29.00) = 3.81 | .0007 | .0054* | Holm-8 |
| Biased-dimension capture: adjust prior-display contrast + group | Pooled (33) | partial r = .559 Fisher [.255, .762] | t(29.00) = 3.63 | .0011 | .0076* | Holm-8 |
| Biased-dimension capture: adjust unbiased | SS (16) | partial r = .610 Fisher [.142, .855] | t(13.00) = 2.77 | .0158 | .0633 | Holm-8 |
| Biased-dimension capture: adjust unbiased | DS (17) | partial r = .531 Fisher [.048, .813] | t(14.00) = 2.35 | .0342 | .1026 | Holm-8 |
| Count-matched biased − unbiased difference | Pooled (33) | Δr = .544 | t(29.00) = 3.04 | .0050 | .0100* | Holm-2 |
| Biased-dimension capture: permutation | Pooled (33) | r = .550 | — | .0016 | — | Unadjusted |
| **B. DMN associations with orientation and color capture** | | | | | | |
| Orientation capture | Pooled (33) | r = .260 Fisher [−.098, .558]  Bootstrap [−.060, .542] | t(30.00) = 1.47 | .1515 | .1732 | Holm-2 |
| Color capture | Pooled (33) | r = .308 Fisher [−.046, .593]  Bootstrap [.009, .558] | t(30.00) = 1.77 | .0866 | .1732 | Holm-2 |
| **C. Location-standardized connectivity–capture associations** | | | | | | |
| Biased-dimension equal-region capture | Pooled (33) | r = .629 Fisher [.358, .802]  Bootstrap [.430, .769] | t(30.00) = 4.43 | .0001 | .0007* | Holm-6 |
| Unbiased-dimension equal-region capture | Pooled (33) | r = .006 Fisher [−.343, .354]  Bootstrap [−.327, .323] | t(30.00) = 0.03 | .9744 | 1.0000 | Holm-6 |
| Biased − unbiased equal-region r | Pooled (33) | Δr = .623 Bootstrap [.343, .911] | t(29.00) = 3.68 | .0009 | .0028* | Holm-3 |
| Biased-dimension equal-region capture | SS (16) | r = .731 Fisher [.368, .900]  Bootstrap [.492, .900] | t(14.00) = 4.00 | .0013 | .0065* | Holm-6 |
| Unbiased-dimension equal-region capture | SS (16) | r = .230 Fisher [−.300, .651]  Bootstrap [−.275, .618] | t(14.00) = 0.88 | .3919 | 1.0000 | Holm-6 |
| Biased-dimension equal-region capture | DS (17) | r = .463 Fisher [−.022, .772]  Bootstrap [.111, .758] | t(15.00) = 2.03 | .0610 | .2440 | Holm-6 |
| Unbiased-dimension equal-region capture | DS (17) | r = −.102 Fisher [−.556, .398]  Bootstrap [−.565, .376] | t(15.00) = −0.40 | .6956 | 1.0000 | Holm-6 |
| Biased − unbiased equal-region r | SS (16) | Δr = .501 Bootstrap [.206, .892] | t(13.00) = 2.33 | .0368 | .0604 | Holm-3 |
| Biased − unbiased equal-region r | DS (17) | Δr = .566 Bootstrap [.121, 1.061] | t(14.00) = 2.41 | .0302 | .0604 | Holm-3 |
| **D. Additional within-group spatial-learning associations** | | | | | | |
| Rare − frequent learning | SS (16) | r = .632 Fisher [.199, .859]  Bootstrap [.119, .867] | t(14.00) = 3.05 | .0086 | .0172* | Holm-3 |
| Rare − frequent learning | DS (17) | r = .296 Fisher [−.215, .680]  Bootstrap [−.233, .706] | t(15.00) = 1.20 | .2483 | .2483 | Holm-3 |

**E. Within-group permutation of the spatial-learning association**

| **Scope (N)** | **Observed r** | **Extreme permutations** | **Raw p** | **Holm-3 p** |
| --- | --- | --- | --- | --- |
| Pooled (33) | .513 | 39 | .0040 | .0120* |
| SS (16) | .632 | 81 | .0082 | .0164* |
| DS (17) | .296 | 2498 | .2499 | .2499 |

*Note (Panels A–D).* Biased = orientation in SS and color in DS; unbiased is the complementary dimension. Pooled Pearson correlations are group centered; partial correlations adjust the covariates named in each row. CIs are pointwise 95% Fisher-z or paired participant-percentile intervals (10,000 draws; pooled resampling stratified by group with recentering). Williams tests compare dependent correlations. All tests are two-sided. Separate families: A, eight secondary capture/partial associations (Holm-8) and two biased–unbiased comparisons (Holm-2); B, two physical-dimension correlations (Holm-2); C, six equal-region correlations (Holm-6) and three correlation differences (Holm-3); D, the pooled/SS/DS learning family (Holm-3; the pooled member is in main Table 3). Complete correction families are retained when members are displayed elsewhere. Equal-region scores weight frequent/rare regions 50/50; neither this weighting nor averaging count-matched scores equalizes measurement precision. The unadjusted capture permutation tests the same primary association. *Holm-adjusted p < .05; pointwise intervals need not agree with adjusted tests. Separate subgroup significance does not establish moderation. *(Panel E)* Learning is biased-dimension rare-minus-frequent RT. Scores are permuted within SS/DS over 10,000 draws; pooled correlations are group centered. Two-sided add-one p = (extreme count + 1)/10,001, using absolute correlations. Holm-3 covers pooled, SS and DS permutation tests, separately from the association and network families. *Holm-adjusted p < .05. This is same-data sensitivity, not independent replication or a group-interaction test.

**Supplementary Table S4. Preprocessing sensitivity checks**

**A. Capture associations**

| **Variant** | **r** | **95% CI** | **Raw p** | **Holm p** |
| --- | --- | --- | --- | --- |
| Primary nuisance regression | .55 | [.25, .75] | .0011 | .0044 |
| No nuisance regression | .55 | [.24, .75] | .0013 | .0044 |
| Primary + global signal regression | .38 | [.03, .64] | .0343 | .0543 |
| Primary + task regression | .56 | [.27, .76] | < .001 | .0040 |
| Primary + task + global signal regression | .39 | [.05, .65] | .0271 | .0543 |

*Note (Panel A).* N = 33, df = 30; pooled group-centered DMN–biased-capture correlations use observed-mixture present-minus-absent RT scores. CIs are approximate pointwise 95% Fisher-z intervals. Primary nuisance regression includes 24 motion regressors plus white-matter and cerebrospinal-fluid signals. Alternatives omit nuisance regression or add global signal, three Glover-HRF-convolved display regressors, or both. Detrending/filtering are unchanged. The five capture variants form one Holm-5 family.

**B. Spatial-learning associations**

| **Variant** | **r [95% Fisher CI]** | **t (df)** | **Raw p** | **Holm-5 p** |
| --- | --- | --- | --- | --- |
| Primary nuisance regression | .513 [.201, .731] | 3.28 (30.00) | .0027 | .0133* |
| No nuisance regression | .358 [.010, .628] | 2.10 (30.00) | .0445 | .0828 |
| Primary + global signal regression | .389 [.046, .649] | 2.31 (30.00) | .0279 | .0828 |
| Primary + task regression | .504 [.189, .725] | 3.20 (30.00) | .0033 | .0133* |
| Primary + task + global signal regression | .389 [.047, .650] | 2.32 (30.00) | .0276 | .0828 |

*Note (Panel B).* N = 33 (SS = 16, DS = 17), df = 30; pooled correlations are group centered. Learning is biased-dimension rare-minus-frequent RT. CIs are approximate pointwise 95% Fisher-z intervals. Holm-5 covers all five learning variants, including primary, separately from the capture family. *Holm-adjusted p < .05. Other processing is fixed; no volume scrubbing, mean-FD adjustment or FD-based exclusion is used. Differences across variants are not direct correlation-difference tests or evidence of mediation.

**C. Explained variance in filtered parcel time series**

| **Model or regressor** | **DMN: 91 parcels  Mean (range), %** | **Whole connectome: 399 parcels  Mean (range), %** |
| --- | --- | --- |
| Primary model: total R² | 45.48 (23.79–70.58) | 54.56 (31.65–74.73) |
| Full model: total R² | 56.94 (32.83–79.26) | 64.31 (42.97–82.40) |
| Motion: ΔR² | 13.71 (6.24–28.33) | 12.94 (8.01–25.84) |
| WM/CSF: ΔR² | 3.26 (1.41–7.23) | 2.29 (1.01–4.31) |
| Global-signal: ΔR² | 10.03 (2.52–25.41) | 8.01 (2.66–17.37) |
| Glover-HRF display: ΔR² | 1.62 (0.30–6.00) | 1.52 (0.49–3.86) |

Note *(Panel C)*. For each participant, run and parcel, we regressed the detrended, band-pass-filtered parcel time series on identically processed nuisance regressors. Explained variance was calculated as R² = 1 − SSE/SST, where SSE is the residual sum of squares and SST is the total sum of squares around the parcel-series mean. The primary model included 26 regressors and the full model included 30. We separately evaluated four regressor sets: motion, WM/CSF, global signal and the three display-event regressors. Each set’s unique contribution was calculated as ΔR² = R²(full model) − R²(model excluding that set), refitting the reduced model while retaining all other regressors. R² values express the percentage of filtered time-series variance explained; ΔR² values express increments in percentage points. Values were averaged across parcels within each domain and run, then equally across all 66 runs, with both runs included for every participant. Ranges represent the minimum and maximum run-level parcel averages, not confidence intervals. Because the regressor sets share explanatory information, their unique contributions do not sum to the full model’s R².

**Supplementary Table S5. Trial-history sensitivity of DMN–capture and DMN–learning coupling**

**A. History-adjusted capture model: fixed effects**

| **Term** | **F** | **Denominator df** | **Raw p** | **Holm-6 p** |
| --- | --- | --- | --- | --- |
| DMN connectivity | .14 | 28.97 | .7139 | 1.0000 |
| Biased dimension × DMN connectivity | 15.89 | 28.33 | < .001 | .0026 |
| Group × DMN connectivity | 1.84 | 29.07 | .1858 | .7434 |
| Unbiased dimension × DMN connectivity | .27 | 27.86 | .6054 | 1.0000 |
| Biased dimension × Group × DMN connectivity | 3.09 | 28.20 | .0895 | .4477 |
| Group × Unbiased dimension × DMN connectivity | .66 | 28.95 | .4227 | 1.0000 |

**B. History-adjusted DMN–capture slopes and contrasts**

| **Contrast** | **Estimate (ms/DMN SD)** | **95% CI** | **t** | **df** | **Raw p** | **Holm-9 p** |
| --- | --- | --- | --- | --- | --- | --- |
| Pooled participant-weighted biased DMN connectivity slope | 20.3 | [8.0, 32.6] | 3.38 | 28.2 | .0022 | .0173 |
| Pooled participant-weighted unbiased DMN connectivity slope | -1.5 | [-20.1, 17.1] | −.16 | 29.0 | .8719 | 1.0000 |
| SS biased DMN connectivity slope | 31.1 | [15.1, 47.1] | 3.99 | 28.3 | < .001 | .0039 |
| DS biased DMN connectivity slope | 10.1 | [-8.5, 28.6] | 1.11 | 28.1 | .2746 | 1.0000 |
| SS unbiased DMN connectivity slope | 6.1 | [-17.8, 30.0] | .52 | 27.9 | .6054 | 1.0000 |
| DS unbiased DMN connectivity slope | -8.6 | [-36.9, 19.6] | −.62 | 29.8 | .5379 | 1.0000 |
| Group moderation biased DS-SS | -21.0 | [-45.5, 3.5] | -1.76 | 28.2 | .0895 | .5373 |
| Group moderation unbiased DS-SS | -14.7 | [-51.7, 22.3] | −.81 | 28.9 | .4227 | 1.0000 |
| Pooled biased-minus-unbiased DMN connectivity slope | 21.7 | [2.9, 40.6] | 2.36 | 29.6 | .0253 | .1770 |

*Note (Panels A–B).* History-adjusted three-category model (Supplementary Equation S12A–C): 10,995 eligible trials, 33 participants; REML/Satterthwaite inference. This sensitivity differs from the principal five-condition model (Supplementary Table S6). Participant-specific run, standardized log recorded order and strictly prior biased/unbiased exposure are controlled. CIs are pointwise 95% t intervals. Connectivity omnibus terms and derived capture slopes/contrasts use separate Holm-6 and Holm-9 families. These adjustments do not separate learning from exposure.

**C. History-adjusted learning model: fixed effects**

| **Term** | **Estimate (ms)** | **SE (ms)** | **95% CI (ms)** | **t (df)** | **Raw p** |
| --- | --- | --- | --- | --- | --- |
| Intercept (SS, frequent, Run 1) | 928.87 | 36.74 | [853.87, 1003.87] | 25.28 (30.31) | < .0001 |
| Rare location | 49.05 | 14.60 | [19.65, 78.45] | 3.36 (45.47) | .0016 |
| DMN connectivity | 30.22 | 34.33 | [−39.83, 100.27] | 0.88 (30.62) | .3856 |
| Group (DS − SS) | −53.27 | 50.63 | [−156.81, 50.28] | −1.05 (29.00) | .3014 |
| Run 2 − Run 1 | −25.11 | 10.64 | [−45.97, −4.24] | −2.36 (6557.55) | .0184 |
| Rare × DMN | 32.51 | 13.96 | [4.48, 60.55] | 2.33 (50.21) | .0239 |
| Rare × Group | −50.52 | 18.16 | [−87.66, −13.38] | −2.78 (28.99) | .0094 |
| DMN × Group | −87.00 | 51.99 | [−193.32, 19.32] | −1.67 (29.01) | .1050 |
| Rare × Run 2 | −1.68 | 12.53 | [−26.25, 22.88] | −0.13 (6561.93) | .8932 |
| DMN × Run 2 | 21.85 | 10.96 | [0.36, 43.34] | 1.99 (6558.53) | .0463 |
| Rare × DMN × Group | −26.64 | 18.68 | [−64.82, 11.55] | −1.43 (29.15) | .1644 |
| Rare × DMN × Run 2 | 16.23 | 12.83 | [−8.92, 41.39] | 1.26 (6557.63) | .2059 |

*Note (Panel C).* The learning model adds participant-specific slopes for three within-participant-standardized history covariates: log recorded order and strictly prior biased/unbiased exposure, constructed before exclusions across runs. The fit uses 6,721 eligible trials from 33 participants (SS = 16, DS = 17), with REML/Satterthwaite inference. Rare, Group and Run2 are coded 0/1 (frequent/rare, SS/DS, runs 1/2); DMN is group centered and scaled by the pooled participant SD. Estimates/SEs are in ms (DMN terms: ms/SD); CIs are pointwise 95% t intervals. Core coefficient tests are unadjusted; participant-specific history slopes are included but not displayed.

**D. History-adjusted learning coupling and direct group contrast**

| **Contrast** | **Estimate** | **SE** | **95% CI** | **t (df)** | **Raw p** | **Holm-4 p** |
| --- | --- | --- | --- | --- | --- | --- |
| Pooled learning coupling | 26.91 | 9.38 | [7.73, 46.09] | 2.87 (29.10) | .0076 | .0228* |
| SS learning coupling | 40.63 | 12.31 | [15.51, 65.75] | 3.30 (30.36) | .0025 | .0099* |
| DS learning coupling | 14.00 | 14.05 | [−14.77, 42.76] | 1.00 (28.28) | .3276 | .3288 |
| Group contrast (DS − SS) | −26.64 | 18.68 | [−64.82, 11.55] | −1.43 (29.15) | .1644 | .3288 |

*Note (Panel D).* Coupling is the modeled rare-minus-frequent RT slope per DMN SD. Runs are averaged equally; pooled SS/DS weights are 16/33 and 17/33. CIs are pointwise 95% Satterthwaite-t intervals using the full coefficient covariance. Holm-4 covers pooled, SS, DS and DS−SS contrasts, separately from the principal-learning and capture families. *Holm-adjusted p < .05. The direct DS−SS contrast tests moderation; history adjustment does not identify causal learning.

**E. Participant OLS learning-score companion (descriptive)**

| **Descriptive association** | **r [95% Fisher CI]** | **t (df)** | **Raw p** |
| --- | --- | --- | --- |
| DMN vs history-adjusted learning | .512 [.199, .730] | 3.26 (30.00) | .0028 |
| Raw vs history-adjusted learning | .996 [.992, .998] | 61.03 (30.00) | < .0001 |

*Note (Panel E).* Participant OLS fits adjust the rare/frequent contrast for run and three history covariates. Correlations are group centered (N = 33, df = 30). Pointwise 95% Fisher intervals and unadjusted Pearson tests are descriptive and do not propagate first-stage score uncertainty. Model-based inference is in Panels C–D.

**Supplementary Table S6. Hierarchical capture model: full coefficients and supporting contrasts**

| **Analysis / contrast** | **Group (N)** | **Estimate [95% CI]** | **Test (df)** | **Raw p** | **Holm p** | **Correction** |
| --- | --- | --- | --- | --- | --- | --- |
| **A. Full fixed-effect coefficients** | | | | | | |
| Intercept | Joint fit (33) | β = 806.02  t-based [734.37, 877.67] | t(29.94)  = 22.98 | <.0001 | — | Unadjusted |
| Biased dimension, frequent region | Joint fit (33) | β = 125.84  t-based [105.83, 145.86] | t(54.54)  = 12.60 | <.0001 | — | Unadjusted |
| Biased dimension, rare region | Joint fit (33) | β = 172.94  t-based [141.27, 204.62] | t(46.68)  = 10.99 | <.0001 | — | Unadjusted |
| Unbiased dimension, frequent region | Joint fit (33) | β = 12.92  t-based [−17.93, 43.76] | t(44.82)  = 0.84 | .4035 | — | Unadjusted |
| Unbiased dimension, rare region | Joint fit (33) | β = 22.38  t-based [−11.91, 56.68] | t(42.80)  = 1.32 | .1951 | — | Unadjusted |
| DMN connectivity | Joint fit (33) | β = 17.72  t-based [−48.61, 84.05] | t(28.96)  = 0.55 | .5889 | — | Unadjusted |
| Group | Joint fit (33) | β = 62.49  t-based [−36.65, 161.63] | t(28.95)  = 1.29 | .2075 | — | Unadjusted |
| Biased dimension, frequent region × DMN connectivity | Joint fit (33) | β = 23.08  t-based [6.87, 39.30] | t(28.57)  = 2.91 | .0069 | — | Unadjusted |
| Biased dimension, rare region × DMN connectivity | Joint fit (33) | β = 64.88  t-based [38.25, 91.51] | t(28.90)  = 4.98 | <.0001 | — | Unadjusted |
| Unbiased dimension, frequent region × DMN connectivity | Joint fit (33) | β = 8.15  t-based [−17.54, 33.84] | t(26.44)  = 0.65 | .5202 | — | Unadjusted |
| Unbiased dimension, rare region × DMN connectivity | Joint fit (33) | β = 1.04  t-based [−28.11, 30.19] | t(27.75)  = 0.07 | .9423 | — | Unadjusted |
| Biased dimension, frequent region × Group | Joint fit (33) | β = −116.72  t-based [−140.88, −92.55] | t(28.40)  = −9.89 | <.0001 | — | Unadjusted |
| Biased dimension, rare region × Group | Joint fit (33) | β = −163.29  t-based [−202.74, −123.84] | t(27.87)  = −8.48 | <.0001 | — | Unadjusted |
| Unbiased dimension, frequent region × Group | Joint fit (33) | β = 200.91  t-based [162.20, 239.62] | t(27.42)  = 10.64 | <.0001 | — | Unadjusted |
| Unbiased dimension, rare region × Group | Joint fit (33) | β = 171.24  t-based [127.28, 215.20] | t(29.02)  = 7.97 | <.0001 | — | Unadjusted |
| DMN connectivity × Group | Joint fit (33) | β = −71.57  t-based [−173.37, 30.24] | t(28.96)  = −1.44 | .1612 | — | Unadjusted |
| Distractor absent × Run 2 | Joint fit (33) | β = −26.54  t-based [−43.98, −9.09] | t(10835.44)  = −2.98 | .0029 | — | Unadjusted |
| Biased dimension, frequent region × Run 2 | Joint fit (33) | β = −29.58  t-based [−39.98, −19.17] | t(10825.84)  = −5.57 | <.0001 | — | Unadjusted |
| Biased dimension, rare region × Run 2 | Joint fit (33) | β = −30.56  t-based [−52.97, −8.15] | t(10838.67)  = −2.67 | .0075 | — | Unadjusted |
| Unbiased dimension, frequent region × Run 2 | Joint fit (33) | β = −40.99  t-based [−63.69, −18.28] | t(10844.12)  = −3.54 | .0004 | — | Unadjusted |
| Unbiased dimension, rare region × Run 2 | Joint fit (33) | β = −36.61  t-based [−59.49, −13.74] | t(10845.07)  = −3.14 | .0017 | — | Unadjusted |
| Biased dimension, frequent region × DMN connectivity × Group | Joint fit (33) | β = −14.55  t-based [−39.39, 10.30] | t(28.47)  = −1.20 | .2406 | — | Unadjusted |
| Biased dimension, rare region × DMN connectivity × Group | Joint fit (33) | β = −43.61  t-based [−84.16, −3.05] | t(27.98)  = −2.20 | .0360 | — | Unadjusted |
| Unbiased dimension, frequent region × DMN connectivity × Group | Joint fit (33) | β = −16.47  t-based [−56.37, 23.43] | t(27.78)  = −0.85 | .4049 | — | Unadjusted |
| Unbiased dimension, rare region × DMN connectivity × Group | Joint fit (33) | β = −3.45  t-based [−48.69, 41.80] | t(29.33)  = −0.16 | .8773 | — | Unadjusted |
| **B. Same-physical-dimension capture costs and group comparisons** | | | | | | |
| Orientation cost (ms) | SS (16) | 147.63 t-based [128.02, 167.24] | t(27.84) = 15.42 | <.0001 | — | Pointwise |
| Orientation cost (ms) | DS (17) | 197.60 t-based [172.50, 222.69] | t(29.71) = 16.09 | <.0001 | — | Pointwise |
| Color cost (ms) | SS (16) | 11.52 t-based [−14.03, 37.07] | t(27.97) = 0.92 | .3635 | — | Pointwise |
| Color cost (ms) | DS (17) | 7.63 t-based [−11.28, 26.53] | t(27.19) = 0.83 | .4153 | — | Pointwise |
| Joint physical RT-cost profile | Pooled (33) | — | F = 6.01  (2, 28.78) | .0066 | .0132* | Holm-2 |
| Orientation: SS−DS cost (ms) | Both (33) | −49.97 t-based [−81.24, −18.69] | t(50.10) = −3.21 | .0023 | .0046* | Holm-2 |
| Color: DS−SS cost (ms) | Both (33) | −3.90 t-based [−35.09, 27.30] | t(46.77) = −0.25 | .8027 | .8027 | Holm-2 |
| **C. Additional subgroup coupling and group contrasts** | | | | | | |
| DMN–biased-dimension capture coupling | SS (16) | 43.98 t-based [25.65, 62.31] | t(28.08) = 4.91 | <.0001 | .0003* | Holm-9 |
| DMN–unbiased-dimension capture coupling | SS (16) | 4.60 t-based [−19.23, 28.42] | t(28.01) = 0.40 | .6958 | 1.0000 | Holm-9 |
| biased − unbiased capture coupling | SS (16) | 39.39 t-based [14.51, 64.26] | t(28.95) = 3.24 | .0030 | .0211* | Holm-9 |
| DMN–biased-dimension capture coupling | DS (17) | 14.91 t-based [−6.27, 36.08] | t(27.27) = 1.44 | .1602 | .8008 | Holm-9 |
| DMN–unbiased-dimension capture coupling | DS (17) | −5.36 t-based [−33.48, 22.75] | t(29.84) = −0.39 | .6995 | 1.0000 | Holm-9 |
| biased − unbiased capture coupling | DS (17) | 20.27 t-based [−8.93, 49.47] | t(30.12) = 1.42 | .1667 | .8008 | Holm-9 |
| Group difference (DS−SS): DMN–unbiased-dimension capture coupling | DS−SS (33) | −9.96 t-based [−46.81, 26.89] | t(29.06) = −0.55 | .5847 | .6336 | Holm-3 |
| Group difference (DS−SS): biased − unbiased capture coupling | DS−SS (33) | −19.12 t-based [−57.48, 19.24] | t(29.62) = −1.02 | .3168 | .6336 | Holm-3 |

*Note.* Principal hierarchical trial-level capture model (Supplementary Equation S13): 10,995 correct trials with finite RTs of 200–3000 ms inclusive, 33 participants (SS = 16, DS = 17); REML/Satterthwaite inference. Estimates are in ms (DMN terms: ms/SD); CIs are pointwise 95% t intervals. Group = 0/1 for SS/DS; Run2 = 0/1 for runs 1/2. DMN is group centered and scaled by the pooled participant SD. Reference values are SS, run 1 and DMN = 0, with the distractor-absent condition as the reference condition. Condition main effects are present-minus-absent costs; condition × DMN terms are coupling increments. Condition × Run2 terms are condition-specific, not absent-referenced, run changes. Raw coefficient tests are unadjusted.

Derived capture contrasts weight frequent/rare regions equally; Panel B also averages runs. Pooled SS/DS weights are 16/33 and 17/33. Orientation is SS biased/DS unbiased; color is complementary. Complete families, including Table 4 members: nine capture slopes/differences (Holm-9), three group contrasts (Holm-3), and three separate Holm-2 families (joint physical profiles, cost comparisons, coupling comparisons). *Holm-adjusted p < .05; pointwise intervals are not corrected decisions. The canonical fit was nonsingular/stable.

**Supplementary Table S7. Hierarchical spatial-learning model: coefficients and additional group-specific slopes**

| **Analysis / contrast** | **Scope (N)** | **Estimate [95% CI]** | **Test (df)** | **Raw p** | **Holm p** | **Correction** |
| --- | --- | --- | --- | --- | --- | --- |
| **A. Full fixed-effect coefficients** | | | | | | |
| Intercept | Joint fit (33) | β = 931.49  t-based [857.02, 1005.97] | t(29.34)  = 25.57 | <.0001 | — | Unadjusted |
| Rare location | Joint fit (33) | β = 47.34  t-based [18.39, 76.29] | t(46.91)  = 3.29 | .0019 | — | Unadjusted |
| DMN connectivity | Joint fit (33) | β = 52.79  t-based [−16.66, 122.24] | t(29.42)  = 1.55 | .1309 | — | Unadjusted |
| Group | Joint fit (33) | β = −53.74  t-based [−157.25, 49.77] | t(29.00)  = −1.06 | .2971 | — | Unadjusted |
| Run 2 | Joint fit (33) | β = −29.55  t-based [−40.08, −19.02] | t(6652.01)  = −5.50 | <.0001 | — | Unadjusted |
| Rare location × DMN connectivity | Joint fit (33) | β = 34.54  t-based [6.91, 62.18] | t(51.96)  = 2.51 | .0153 | — | Unadjusted |
| Rare location × Group | Joint fit (33) | β = −46.77  t-based [−83.06, −10.47] | t(28.93)  = −2.64 | .0134 | — | Unadjusted |
| DMN connectivity × Group | Joint fit (33) | β = −86.38  t-based [−192.67, 19.91] | t(29.01)  = −1.66 | .1073 | — | Unadjusted |
| Rare location × Run 2 | Joint fit (33) | β = −1.22  t-based [−26.23, 23.80] | t(6660.79)  = −0.10 | .9241 | — | Unadjusted |
| DMN connectivity × Run 2 | Joint fit (33) | β = −22.86  t-based [−33.61, −12.12] | t(6651.83)  = −4.17 | <.0001 | — | Unadjusted |
| Rare location × DMN connectivity × Group | Joint fit (33) | β = −29.04  t-based [−66.35, 8.28] | t(29.06)  = −1.59 | .1224 | — | Unadjusted |
| Rare location × DMN connectivity × Run 2 | Joint fit (33) | β = 13.79  t-based [−11.82, 39.41] | t(6656.41)  = 1.06 | .2913 | — | Unadjusted |
| **B. Additional equal-run DMN–learning coupling by group** | | | | | | |
| DMN–rare-minus-frequent learning coupling | SS (16) | 41.44 t-based [16.88, 66.00] | t(30.32) = 3.44 | .0017 | .0068* | Holm-4 |
| DMN–rare-minus-frequent learning coupling | DS (17) | 12.40 t-based [−15.69, 40.50] | t(28.14) = 0.90 | .3735 | .3735 | Holm-4 |
| **C. Run-moderated fit: six run-specific companion slopes** | | | | | | |
| Run 1 learning slope | Pooled (33) | 19.59 t-based [−3.02, 42.19] | t(67.54) = 1.73 | .0883 | .2650 | Holm-6 |
| Run 1 learning slope | SS (16) | 34.54 t-based [6.91, 62.18] | t(51.96) = 2.51 | .0153 | .0611 | Holm-6 |
| Run 1 learning slope | DS (17) | 5.51 t-based [−25.21, 36.23] | t(42.75) = 0.36 | .7194 | .7194 | Holm-6 |
| Run 2 learning slope | Pooled (33) | 33.38 t-based [11.04, 55.72] | t(64.24) = 2.98 | .0040 | .0201* | Holm-6 |
| Run 2 learning slope | SS (16) | 48.34 t-based [20.99, 75.69] | t(49.66) = 3.55 | .0009 | .0051* | Holm-6 |
| Run 2 learning slope | DS (17) | 19.30 t-based [−11.29, 49.89] | t(41.95) = 1.27 | .2099 | .4198 | Holm-6 |

*Note.* Run-moderated learning model (Supplementary Equation S14): 6,721 correct biased-dimension trials with finite RTs of 200–3000 ms inclusive, 33 participants (SS = 16, DS = 17); REML/Satterthwaite inference. Learning uses orientation in SS and color in DS. Rare, Group and Run2 are coded 0/1 (frequent/rare, SS/DS, runs 1/2); the intercept and DMN coefficient refer to frequent SS trials in run 1. DMN is group centered and scaled by the pooled participant SD. Estimates are in ms (DMN terms: ms/SD); CIs are pointwise 95% t intervals. Raw coefficient tests are unadjusted.

Learning coupling is the modeled rare-minus-frequent RT slope per DMN SD. Equal-run estimates average both runs; pooled SS/DS weights are 16/33 and 17/33. Pooled, SS, DS and DS−SS contrasts retain Holm-4; the six run-specific slopes retain separate Holm-6. Rare × DMN × Run2 tests the common run change; no group-specific run interaction is fitted. DMN is averaged across task runs, not a within-person change measure. *Holm-adjusted p < .05; separate subgroup/run significance does not establish an interaction.

**Supplementary Table S8. Capture network-summary associations and direct comparisons**

**A. All 28 pooled network summaries**

| **Network pair** | **Edge class** | **Possible edges** | **r** | **95% CI** | **Raw p** | **Holm-28 p** | **Perm. p**  **Raw** | **Perm. p**  **maxT-28** |
| --- | --- | --- | --- | --- | --- | --- | --- | --- |
| Default-mode – Default-mode | Within network | 4095 | .55 | [.25, .75] | .0011 | .0308 | 0.0012 | 0.0044 |
| Default-mode – Somatomotor | Between networks | 7007 | .20 | [−.16, .51] | .2703 | 1.0000 | 0.1865 | 0.9267 |
| Default-mode – Visual | Between networks | 5551 | .13 | [−.23, .46] | .4732 | 1.0000 | 0.4156 | 0.9982 |
| Default-mode – Frontoparietal control | Between networks | 4732 | .34 | [−.01, .62] | .0553 | 1.0000 | 0.0414 | 0.3206 |
| Default-mode – Ventral attention/salience | Between networks | 4277 | .17 | [−.19, .49] | .3589 | 1.0000 | 0.3166 | 0.9801 |
| Default-mode – Dorsal attention | Between networks | 4186 | .13 | [−.23, .46] | .4770 | 1.0000 | 0.4270 | 0.9982 |
| Default-mode – Limbic | Between networks | 2275 | .47 | [.15, .70] | .0065 | .1760 | 0.0030 | 0.0419 |
| Somatomotor – Somatomotor | Within network | 2926 | .12 | [−.24, .45] | .5178 | 1.0000 | 0.4757 | 0.9990 |
| Somatomotor – Visual | Between networks | 4697 | .04 | [−.31, .39] | .8077 | 1.0000 | 0.7687 | 1.0000 |
| Somatomotor – Frontoparietal control | Between networks | 4004 | .16 | [−.20, .48] | .3795 | 1.0000 | 0.2230 | 0.9863 |
| Somatomotor – Ventral attention/salience | Between networks | 3619 | .12 | [−.24, .45] | .5134 | 1.0000 | 0.4515 | 0.9989 |
| Somatomotor – Dorsal attention | Between networks | 3542 | .07 | [−.29, .41] | .7144 | 1.0000 | 0.6582 | 1.0000 |
| Somatomotor – Limbic | Between networks | 1925 | .10 | [−.26, .43] | .5903 | 1.0000 | 0.5333 | 0.9999 |
| Visual – Visual | Within network | 1830 | .11 | [−.25, .44] | .5536 | 1.0000 | 0.5404 | 0.9994 |
| Visual – Frontoparietal control | Between networks | 3172 | .04 | [−.32, .38] | .8485 | 1.0000 | 0.8218 | 1.0000 |
| Visual – Ventral attention/salience | Between networks | 2867 | .09 | [−.27, .43] | .6166 | 1.0000 | 0.5747 | 0.9999 |
| Visual – Dorsal attention | Between networks | 2806 | −.01 | [−.35, .34] | .9709 | 1.0000 | 0.9673 | 1.0000 |
| Visual – Limbic | Between networks | 1525 | .03 | [−.32, .38] | .8528 | 1.0000 | 0.8289 | 1.0000 |
| Frontoparietal control – Frontoparietal control | Within network | 1326 | .21 | [−.15, .52] | .2517 | 1.0000 | 0.2374 | 0.9059 |
| Frontoparietal control – Ventral attention/salience | Between networks | 2444 | .17 | [−.19, .49] | .3618 | 1.0000 | 0.2373 | 0.9811 |
| Frontoparietal control – Dorsal attention | Between networks | 2392 | .10 | [−.26, .43] | .6041 | 1.0000 | 0.5438 | 0.9999 |
| Frontoparietal control – Limbic | Between networks | 1300 | .32 | [−.04, .60] | .0783 | 1.0000 | 0.0641 | 0.4358 |
| Ventral attention/salience – Ventral attention/salience | Within network | 1081 | .12 | [−.24, .45] | .5125 | 1.0000 | 0.4312 | 0.9989 |
| Ventral attention/salience – Dorsal attention | Between networks | 2162 | .10 | [−.25, .44] | .5700 | 1.0000 | 0.5194 | 0.9996 |
| Ventral attention/salience – Limbic | Between networks | 1175 | .05 | [−.30, .39] | .7773 | 1.0000 | 0.7551 | 1.0000 |
| Dorsal attention – Dorsal attention | Within network | 1035 | .05 | [−.30, .40] | .7679 | 1.0000 | 0.7631 | 1.0000 |
| Dorsal attention – Limbic | Between networks | 1150 | .02 | [−.33, .37] | .8978 | 1.0000 | 0.8827 | 1.0000 |
| Limbic – Limbic | Within network | 300 | .17 | [−.19, .49] | .3411 | 1.0000 | 0.3598 | 0.9736 |

**B. DMN versus six other within-network summaries (DMN r = .55)**

| **Comparator** | **Comparator r** | **Δr** | **Williams t** | **df** | **Raw p** | **Holm-6 p** |
| --- | --- | --- | --- | --- | --- | --- |
| Somatomotor | .12 | .43 | 2.19 | 29 | .0367 | .0948 |
| Visual | .11 | .44 | 2.30 | 29 | .0288 | .0948 |
| Frontoparietal control | .21 | .34 | 2.39 | 29 | .0237 | .0948 |
| Ventral attention/salience | .12 | .43 | 2.74 | 29 | .0104 | .0519 |
| Dorsal attention | .05 | .50 | 3.11 | 29 | .0041 | .0248 |
| Limbic | .17 | .38 | 2.30 | 29 | .0289 | .0948 |

*Note. N = 33 (SS = 16, DS = 17); pooled correlations are group adjusted. Panel A retains all 28 summaries under Holm-28; its 95% Fisher intervals are approximate and pointwise. Panel B uses six dependent-correlation Williams comparisons under separate Holm-6, with a pooled group-adjusted approximation. Δr = DMN r minus comparator r.*

**Supplementary Table S9. Signed-edge enrichment for capture**

**A. Positive-edge concentration**

| **Scope** | **Category** | **Possible edges** | **Observed edges** | **Expected edges** | **Observed /**  **expected** | **Raw p** | **Holm p** | **Perm. p**  **Raw** | **Perm. p**  **maxZ**  **56/14** |
| --- | --- | --- | --- | --- | --- | --- | --- | --- | --- |
| Network pair | Default-mode – Default-mode | 4095 | 226 | 41.9 | 5.40 | .0015 | .0840 | 0.0015 | 0.0761 |
| Network pair | Default-mode – Somatomotor | 7007 | 64 | 71.7 | .89 | .4771 | 1.0000 | 0.4771 | 1.0000 |
| Network pair | Default-mode – Visual | 5551 | 20 | 56.8 | .35 | .9764 | 1.0000 | 0.9764 | 1.0000 |
| Network pair | Default-mode – Frontoparietal control | 4732 | 137 | 48.4 | 2.83 | .0147 | .7790 | 0.0147 | 0.4075 |
| Network pair | Default-mode – Ventral attention/salience | 4277 | 34 | 43.7 | .78 | .8542 | 1.0000 | 0.8542 | 1.0000 |
| Network pair | Default-mode – Dorsal attention | 4186 | 21 | 42.8 | .49 | .9627 | 1.0000 | 0.9627 | 1.0000 |
| Network pair | Default-mode – Limbic | 2275 | 81 | 23.3 | 3.48 | .0044 | .2420 | 0.0044 | 0.2540 |
| Network pair | Somatomotor – Somatomotor | 2926 | 7 | 29.9 | .23 | .9442 | 1.0000 | 0.9442 | 1.0000 |
| Network pair | Somatomotor – Visual | 4697 | 11 | 48.0 | .23 | .9657 | 1.0000 | 0.9657 | 1.0000 |
| Network pair | Somatomotor – Frontoparietal control | 4004 | 17 | 40.9 | .42 | .9341 | 1.0000 | 0.9341 | 1.0000 |
| Network pair | Somatomotor – Ventral attention/salience | 3619 | 14 | 37.0 | .38 | .9434 | 1.0000 | 0.9434 | 1.0000 |
| Network pair | Somatomotor – Dorsal attention | 3542 | 13 | 36.2 | .36 | .9582 | 1.0000 | 0.9582 | 1.0000 |
| Network pair | Somatomotor – Limbic | 1925 | 3 | 19.7 | .15 | .9711 | 1.0000 | 0.9711 | 1.0000 |
| Network pair | Visual – Visual | 1830 | 5 | 18.7 | .27 | .9520 | 1.0000 | 0.9520 | 1.0000 |
| Network pair | Visual – Frontoparietal control | 3172 | 8 | 32.4 | .25 | .9675 | 1.0000 | 0.9675 | 1.0000 |
| Network pair | Visual – Ventral attention/salience | 2867 | 18 | 29.3 | .61 | .9210 | 1.0000 | 0.9210 | 1.0000 |
| Network pair | Visual – Dorsal attention | 2806 | 7 | 28.7 | .24 | .9678 | 1.0000 | 0.9678 | 1.0000 |
| Network pair | Visual – Limbic | 1525 | 4 | 15.6 | .26 | .9633 | 1.0000 | 0.9633 | 1.0000 |
| Network pair | Frontoparietal control – Frontoparietal control | 1326 | 16 | 13.6 | 1.18 | .2648 | 1.0000 | 0.2648 | 1.0000 |
| Network pair | Frontoparietal control – Ventral attention/salience | 2444 | 16 | 25.0 | .64 | .9108 | 1.0000 | 0.9108 | 1.0000 |
| Network pair | Frontoparietal control – Dorsal attention | 2392 | 12 | 24.5 | .49 | .9465 | 1.0000 | 0.9465 | 1.0000 |
| Network pair | Frontoparietal control – Limbic | 1300 | 44 | 13.3 | 3.31 | .0074 | .3996 | 0.0074 | 0.2905 |
| Network pair | Ventral attention/salience – Ventral attention/salience | 1081 | 13 | 11.1 | 1.18 | .2125 | 1.0000 | 0.2125 | 1.0000 |
| Network pair | Ventral attention/salience – Dorsal attention | 2162 | 11 | 22.1 | .50 | .9300 | 1.0000 | 0.9300 | 1.0000 |
| Network pair | Ventral attention/salience – Limbic | 1175 | 0 | 12.0 | .00 | .9757 | 1.0000 | 0.9757 | 1.0000 |
| Network pair | Dorsal attention – Dorsal attention | 1035 | 6 | 10.6 | .57 | .9108 | 1.0000 | 0.9108 | 1.0000 |
| Network pair | Dorsal attention – Limbic | 1150 | 1 | 11.8 | .09 | .9764 | 1.0000 | 0.9764 | 1.0000 |
| Network pair | Limbic – Limbic | 300 | 3 | 3.1 | .98 | .3809 | 1.0000 | 0.3809 | 1.0000 |
| Network endpoint | Default-mode | 32123 | 583 | 328.5 | 1.77 | .0158 | .2212 | 0.0158 | 0.1616 |
| Network endpoint | Somatomotor | 27720 | 129 | 283.5 | .46 | .9710 | 1.0000 | 0.9710 | 1.0000 |
| Network endpoint | Visual | 22448 | 73 | 229.6 | .32 | .9911 | 1.0000 | 0.9911 | 1.0000 |
| Network endpoint | Frontoparietal control | 19370 | 250 | 198.1 | 1.26 | .0836 | 1.0000 | 0.0836 | 0.6956 |
| Network endpoint | Ventral attention/salience | 17625 | 106 | 180.2 | .59 | .9711 | 1.0000 | 0.9711 | 1.0000 |
| Network endpoint | Dorsal attention | 17273 | 71 | 176.6 | .40 | .9911 | 1.0000 | 0.9911 | 1.0000 |
| Network endpoint | Limbic | 9650 | 136 | 98.7 | 1.38 | .0791 | 1.0000 | 0.0791 | 0.6451 |

**B. Negative-edge concentration**

| **Scope** | **Category** | **Possible edges** | **Observed edges** | **Expected edges** | **Observed /**  **expected** | **Raw p** | **Holm p** | **Perm. p**  **Raw** | **Perm. p**  **maxZ**  **56/14** |
| --- | --- | --- | --- | --- | --- | --- | --- | --- | --- |
| Network pair | Default-mode – Default-mode | 4095 | 0 | 3.6 | .00 | .7375 | 1.0000 | 0.7375 | 1.0000 |
| Network pair | Default-mode – Somatomotor | 7007 | 1 | 6.2 | .16 | .3687 | 1.0000 | 0.3687 | 1.0000 |
| Network pair | Default-mode – Visual | 5551 | 0 | 4.9 | .00 | .5965 | 1.0000 | 0.5965 | 1.0000 |
| Network pair | Default-mode – Frontoparietal control | 4732 | 0 | 4.2 | .00 | .7522 | 1.0000 | 0.7522 | 1.0000 |
| Network pair | Default-mode – Ventral attention/salience | 4277 | 1 | 3.8 | .27 | .5946 | 1.0000 | 0.5946 | 1.0000 |
| Network pair | Default-mode – Dorsal attention | 4186 | 0 | 3.7 | .00 | .5743 | 1.0000 | 0.5743 | 1.0000 |
| Network pair | Default-mode – Limbic | 2275 | 1 | 2.0 | .50 | .5800 | 1.0000 | 0.5800 | 1.0000 |
| Network pair | Somatomotor – Somatomotor | 2926 | 0 | 2.6 | .00 | .5398 | 1.0000 | 0.5398 | 1.0000 |
| Network pair | Somatomotor – Visual | 4697 | 1 | 4.1 | .24 | .4340 | 1.0000 | 0.4340 | 1.0000 |
| Network pair | Somatomotor – Frontoparietal control | 4004 | 1 | 3.5 | .28 | .3608 | 1.0000 | 0.3608 | 1.0000 |
| Network pair | Somatomotor – Ventral attention/salience | 3619 | 11 | 3.2 | 3.45 | .1429 | 1.0000 | 0.1429 | 0.9992 |
| Network pair | Somatomotor – Dorsal attention | 3542 | 2 | 3.1 | .64 | .4145 | 1.0000 | 0.4145 | 1.0000 |
| Network pair | Somatomotor – Limbic | 1925 | 2 | 1.7 | 1.18 | .2594 | 1.0000 | 0.2594 | 1.0000 |
| Network pair | Visual – Visual | 1830 | 0 | 1.6 | .00 | .6975 | 1.0000 | 0.6975 | 1.0000 |
| Network pair | Visual – Frontoparietal control | 3172 | 3 | 2.8 | 1.07 | .2504 | 1.0000 | 0.2504 | 1.0000 |
| Network pair | Visual – Ventral attention/salience | 2867 | 3 | 2.5 | 1.19 | .3125 | 1.0000 | 0.3125 | 1.0000 |
| Network pair | Visual – Dorsal attention | 2806 | 4 | 2.5 | 1.62 | .2891 | 1.0000 | 0.2891 | 1.0000 |
| Network pair | Visual – Limbic | 1525 | 1 | 1.3 | .74 | .3117 | 1.0000 | 0.3117 | 1.0000 |
| Network pair | Frontoparietal control – Frontoparietal control | 1326 | 2 | 1.2 | 1.71 | .3775 | 1.0000 | 0.3775 | 1.0000 |
| Network pair | Frontoparietal control – Ventral attention/salience | 2444 | 9 | 2.2 | 4.18 | .0889 | 1.0000 | 0.0889 | 0.9727 |
| Network pair | Frontoparietal control – Dorsal attention | 2392 | 6 | 2.1 | 2.85 | .2366 | 1.0000 | 0.2366 | 1.0000 |
| Network pair | Frontoparietal control – Limbic | 1300 | 3 | 1.1 | 2.62 | .2249 | 1.0000 | 0.2249 | 1.0000 |
| Network pair | Ventral attention/salience – Ventral attention/salience | 1081 | 5 | 1.0 | 5.25 | .1483 | 1.0000 | 0.1483 | 0.9988 |
| Network pair | Ventral attention/salience – Dorsal attention | 2162 | 7 | 1.9 | 3.67 | .1818 | 1.0000 | 0.1818 | 1.0000 |
| Network pair | Ventral attention/salience – Limbic | 1175 | 3 | 1.0 | 2.90 | .1764 | 1.0000 | 0.1764 | 1.0000 |
| Network pair | Dorsal attention – Dorsal attention | 1035 | 2 | .9 | 2.19 | .3912 | 1.0000 | 0.3912 | 1.0000 |
| Network pair | Dorsal attention – Limbic | 1150 | 0 | 1.0 | .00 | .5298 | 1.0000 | 0.5298 | 1.0000 |
| Network pair | Limbic – Limbic | 300 | 2 | .3 | 7.56 | .1622 | 1.0000 | 0.1622 | 0.9992 |
| Network endpoint | Default-mode | 32123 | 3 | 28.3 | .11 | .7864 | 1.0000 | 0.7864 | 1.0000 |
| Network endpoint | Somatomotor | 27720 | 18 | 24.4 | .74 | .3420 | 1.0000 | 0.3420 | 0.9967 |
| Network endpoint | Visual | 22448 | 12 | 19.8 | .61 | .6100 | 1.0000 | 0.6100 | 1.0000 |
| Network endpoint | Frontoparietal control | 19370 | 24 | 17.1 | 1.41 | .3342 | 1.0000 | 0.3342 | 0.9999 |
| Network endpoint | Ventral attention/salience | 17625 | 39 | 15.5 | 2.51 | .1543 | 1.0000 | 0.1543 | 0.8840 |
| Network endpoint | Dorsal attention | 17273 | 21 | 15.2 | 1.38 | .3400 | 1.0000 | 0.3400 | 1.0000 |
| Network endpoint | Limbic | 9650 | 12 | 8.5 | 1.41 | .3534 | 1.0000 | 0.3534 | 1.0000 |

*Note.* The raw two-sided p < .01 screen covers 79,401 edges, retaining 812 positive and 70 negative edges. Pair categories are exclusive; endpoint categories overlap and must not be summed. Expected counts reflect possible-edge capacity; observed/expected ratios are descriptive. Each of 10,000 within-group permutations repeats the signed screen. Add-one upper-tail tests use separate Holm-56 (28 pairs × two signs). Unadjusted permutation p and exploratory maxZ-adjusted p: 56 pair/sign tests for network pairs and 14 endpoint/sign tests for network endpoints, within capture. Both signs share each family.

**Supplementary Table S10. Spatial-learning network summaries and pooled signed-edge counts**

**A. Pooled network-summary associations (N33, df30)**

| **Network pair** | **r [95% CI]** | **t** | **Raw p** | **Holm-28 p** | **Perm. p**  **Raw** | **Perm. p**  **maxT-28** |
| --- | --- | --- | --- | --- | --- | --- |
| Default-mode – Default-mode | .513 [.201, .731] | 3.276 | .0027 | .0745 | 0.0042 | 0.0215 |
| Default-mode – Somatomotor | .158 [−.202, .480] | .877 | .3873 | 1.0000 | 0.3269 | 0.9904 |
| Default-mode – Visual | .110 [−.248, .442] | .606 | .5491 | 1.0000 | 0.5104 | 0.9999 |
| Default-mode – Frontoparietal control | .295 [−.060, .583] | 1.689 | .1017 | 1.0000 | 0.0891 | 0.5598 |
| Default-mode – Ventral attention/salience | .195 [−.165, .509] | 1.089 | .2849 | 1.0000 | 0.2552 | 0.9449 |
| Default-mode – Dorsal attention | .092 [−.265, .427] | .505 | .6173 | 1.0000 | 0.5896 | 1.0000 |
| Default-mode – Limbic | .343 [−.006, .618] | 2.003 | .0543 | 1.0000 | 0.0381 | 0.3476 |
| Somatomotor – Somatomotor | .114 [−.245, .445] | .627 | .5352 | 1.0000 | 0.5051 | 0.9998 |
| Somatomotor – Visual | .069 [−.287, .408] | .379 | .7072 | 1.0000 | 0.6650 | 1.0000 |
| Somatomotor – Frontoparietal control | .151 [−.209, .475] | .836 | .4096 | 1.0000 | 0.3050 | 0.9943 |
| Somatomotor – Ventral attention/salience | .200 [−.160, .513] | 1.119 | .2720 | 1.0000 | 0.2276 | 0.9337 |
| Somatomotor – Dorsal attention | .144 [−.216, .469] | .796 | .4325 | 1.0000 | 0.3729 | 0.9966 |
| Somatomotor – Limbic | .119 [−.239, .449] | .659 | .5152 | 1.0000 | 0.4567 | 0.9998 |
| Visual – Visual | .014 [−.337, .361] | .074 | .9412 | 1.0000 | 0.9417 | 1.0000 |
| Visual – Frontoparietal control | .037 [−.316, .381] | .203 | .8401 | 1.0000 | 0.8176 | 1.0000 |
| Visual – Ventral attention/salience | .061 [−.294, .401] | .335 | .7396 | 1.0000 | 0.7241 | 1.0000 |
| Visual – Dorsal attention | −.006 [−.354, .344] | −.032 | .9746 | 1.0000 | 0.9687 | 1.0000 |
| Visual – Limbic | .064 [−.291, .404] | .352 | .7276 | 1.0000 | 0.6916 | 1.0000 |
| Frontoparietal control – Frontoparietal control | .143 [−.216, .469] | .794 | .4334 | 1.0000 | 0.4340 | 0.9967 |
| Frontoparietal control – Ventral attention/salience | .143 [−.217, .468] | .789 | .4362 | 1.0000 | 0.3551 | 0.9968 |
| Frontoparietal control – Dorsal attention | .062 [−.293, .402] | .341 | .7358 | 1.0000 | 0.7088 | 1.0000 |
| Frontoparietal control – Limbic | .141 [−.219, .467] | .779 | .4422 | 1.0000 | 0.4324 | 0.9973 |
| Ventral attention/salience – Ventral attention/salience | .252 [−.106, .552] | 1.427 | .1640 | 1.0000 | 0.1135 | 0.7585 |
| Ventral attention/salience – Dorsal attention | .175 [−.185, .494] | .976 | .3368 | 1.0000 | 0.2994 | 0.9748 |
| Ventral attention/salience – Limbic | .056 [−.299, .397] | .306 | .7617 | 1.0000 | 0.7464 | 1.0000 |
| Dorsal attention – Dorsal attention | .143 [−.216, .469] | .794 | .4335 | 1.0000 | 0.4212 | 0.9967 |
| Dorsal attention – Limbic | −.023 [−.369, .328] | −.127 | .8999 | 1.0000 | 0.8868 | 1.0000 |
| Limbic – Limbic | .395 [.054, .654] | 2.357 | .0251 | .6789 | 0.0318 | 0.1836 |

**B. SS network-summary associations (N16, df14)**

| **Network pair** | **r [95% CI]** | **t** | **Raw p** | **Holm-28 p** | **Perm. p**  **Raw** | **Perm. p**  **maxT-28** |
| --- | --- | --- | --- | --- | --- | --- |
| Default-mode – Default-mode | .632 [.199, .859] | 3.054 | .0086 | .2316 | 0.0101 | 0.1220 |
| Default-mode – Somatomotor | .452 [−.057, .774] | 1.894 | .0791 | 1.0000 | 0.0771 | 0.5805 |
| Default-mode – Visual | .334 [−.194, .712] | 1.327 | .2059 | 1.0000 | 0.2017 | 0.9100 |
| Default-mode – Frontoparietal control | .525 [.040, .810] | 2.310 | .0367 | .8430 | 0.0363 | 0.3491 |
| Default-mode – Ventral attention/salience | .481 [−.019, .789] | 2.052 | .0593 | 1.0000 | 0.0562 | 0.4802 |
| Default-mode – Dorsal attention | .417 [−.099, .756] | 1.718 | .1078 | 1.0000 | 0.1052 | 0.6934 |
| Default-mode – Limbic | .389 [−.132, .742] | 1.580 | .1365 | 1.0000 | 0.1314 | 0.7785 |
| Somatomotor – Somatomotor | .340 [−.188, .715] | 1.351 | .1983 | 1.0000 | 0.1965 | 0.9016 |
| Somatomotor – Visual | .563 [.093, .828] | 2.549 | .0232 | .5563 | 0.0241 | 0.2507 |
| Somatomotor – Frontoparietal control | .677 [.273, .878] | 3.445 | .0039 | .1104 | 0.0032 | 0.0649 |
| Somatomotor – Ventral attention/salience | .600 [.148, .844] | 2.806 | .0140 | .3646 | 0.0163 | 0.1736 |
| Somatomotor – Dorsal attention | .599 [.147, .844] | 2.801 | .0141 | .3646 | 0.0156 | 0.1744 |
| Somatomotor – Limbic | .444 [−.067, .770] | 1.852 | .0853 | 1.0000 | 0.0856 | 0.6080 |
| Visual – Visual | .135 [−.386, .591] | .512 | .6169 | 1.0000 | 0.6163 | 1.0000 |
| Visual – Frontoparietal control | .228 [−.302, .650] | .876 | .3960 | 1.0000 | 0.3953 | 0.9971 |
| Visual – Ventral attention/salience | .354 [−.172, .723] | 1.416 | .1786 | 1.0000 | 0.1798 | 0.8709 |
| Visual – Dorsal attention | .305 [−.225, .695] | 1.197 | .2511 | 1.0000 | 0.2521 | 0.9546 |
| Visual – Limbic | .394 [−.126, .744] | 1.604 | .1311 | 1.0000 | 0.1313 | 0.7629 |
| Frontoparietal control – Frontoparietal control | .228 [−.302, .650] | .877 | .3955 | 1.0000 | 0.3905 | 0.9970 |
| Frontoparietal control – Ventral attention/salience | .498 [.002, .797] | 2.146 | .0499 | 1.0000 | 0.0511 | 0.4307 |
| Frontoparietal control – Dorsal attention | .257 [−.273, .668] | .997 | .3356 | 1.0000 | 0.3347 | 0.9910 |
| Frontoparietal control – Limbic | .279 [−.251, .681] | 1.088 | .2949 | 1.0000 | 0.2950 | 0.9776 |
| Ventral attention/salience – Ventral attention/salience | .503 [.010, .799] | 2.177 | .0471 | 1.0000 | 0.0505 | 0.4141 |
| Ventral attention/salience – Dorsal attention | .449 [−.060, .773] | 1.880 | .0810 | 1.0000 | 0.0824 | 0.5902 |
| Ventral attention/salience – Limbic | .279 [−.252, .681] | 1.087 | .2954 | 1.0000 | 0.2957 | 0.9780 |
| Dorsal attention – Dorsal attention | .207 [−.321, .638] | .793 | .4410 | 1.0000 | 0.4379 | 0.9995 |
| Dorsal attention – Limbic | .294 [−.236, .689] | 1.152 | .2688 | 1.0000 | 0.2744 | 0.9654 |
| Limbic – Limbic | .440 [−.071, .768] | 1.835 | .0879 | 1.0000 | 0.0885 | 0.6180 |

**C. DS network-summary associations (N17, df15)**

| **Network pair** | **r [95% CI]** | **t** | **Raw p** | **Holm-28 p** | **Perm. p**  **Raw** | **Perm. p**  **maxT-28** |
| --- | --- | --- | --- | --- | --- | --- |
| Default-mode – Default-mode | .296 [−.215, .680] | 1.201 | .2483 | 1.0000 | 0.2489 | 0.8999 |
| Default-mode – Somatomotor | −.129 [−.574, .375] | −.504 | .6214 | 1.0000 | 0.6252 | 0.9999 |
| Default-mode – Visual | −.149 [−.587, .358] | −.582 | .5692 | 1.0000 | 0.5628 | 0.9998 |
| Default-mode – Frontoparietal control | −.021 [−.496, .465] | −.080 | .9375 | 1.0000 | 0.9375 | 1.0000 |
| Default-mode – Ventral attention/salience | −.171 [−.602, .337] | −.673 | .5110 | 1.0000 | 0.5140 | 0.9992 |
| Default-mode – Dorsal attention | −.324 [−.696, .186] | −1.324 | .2052 | 1.0000 | 0.2070 | 0.8382 |
| Default-mode – Limbic | .349 [−.158, .711] | 1.444 | .1692 | 1.0000 | 0.1719 | 0.7723 |
| Somatomotor – Somatomotor | −.153 [−.590, .354] | −.600 | .5573 | 1.0000 | 0.5586 | 0.9997 |
| Somatomotor – Visual | −.432 [−.756, .061] | −1.855 | .0834 | 1.0000 | 0.0829 | 0.5202 |
| Somatomotor – Frontoparietal control | −.196 [−.618, .315] | −.773 | .4517 | 1.0000 | 0.4578 | 0.9963 |
| Somatomotor – Ventral attention/salience | −.247 [−.650, .265] | −.987 | .3393 | 1.0000 | 0.3458 | 0.9718 |
| Somatomotor – Dorsal attention | −.315 [−.691, .195] | −1.287 | .2176 | 1.0000 | 0.2160 | 0.8565 |
| Somatomotor – Limbic | −.225 [−.637, .287] | −.893 | .3857 | 1.0000 | 0.3890 | 0.9864 |
| Visual – Visual | −.179 [−.607, .330] | −.705 | .4917 | 1.0000 | 0.4928 | 0.9986 |
| Visual – Frontoparietal control | −.183 [−.610, .326] | −.720 | .4824 | 1.0000 | 0.4793 | 0.9982 |
| Visual – Ventral attention/salience | −.319 [−.694, .191] | −1.305 | .2116 | 1.0000 | 0.2123 | 0.8489 |
| Visual – Dorsal attention | −.425 [−.752, .070] | −1.819 | .0889 | 1.0000 | 0.0916 | 0.5448 |
| Visual – Limbic | −.307 [−.686, .204] | −1.247 | .2314 | 1.0000 | 0.2322 | 0.8771 |
| Frontoparietal control – Frontoparietal control | .027 [−.460, .501] | .105 | .9181 | 1.0000 | 0.9226 | 1.0000 |
| Frontoparietal control – Ventral attention/salience | −.136 [−.579, .369] | −.530 | .6040 | 1.0000 | 0.6104 | 0.9999 |
| Frontoparietal control – Dorsal attention | −.159 [−.594, .349] | −.623 | .5429 | 1.0000 | 0.5491 | 0.9997 |
| Frontoparietal control – Limbic | −.061 [−.526, .432] | −.238 | .8150 | 1.0000 | 0.8193 | 1.0000 |
| Ventral attention/salience – Ventral attention/salience | .066 [−.428, .530] | .256 | .8017 | 1.0000 | 0.8066 | 1.0000 |
| Ventral attention/salience – Dorsal attention | −.121 [−.568, .382] | −.470 | .6450 | 1.0000 | 0.6464 | 1.0000 |
| Ventral attention/salience – Limbic | −.237 [−.644, .275] | −.945 | .3597 | 1.0000 | 0.3551 | 0.9800 |
| Dorsal attention – Dorsal attention | .066 [−.428, .530] | .256 | .8012 | 1.0000 | 0.8038 | 1.0000 |
| Dorsal attention – Limbic | −.426 [−.753, .069] | −1.824 | .0882 | 1.0000 | 0.0879 | 0.5418 |
| Limbic – Limbic | .341 [−.167, .706] | 1.407 | .1799 | 1.0000 | 0.1810 | 0.7935 |

**D. Pooled descriptive raw-screen edge counts**

| **Network pair** | **Possible edges** | **Positive count** | **Negative count** | **Positive share (%)** | **Negative share (%)** | **Perm. p**  **Raw**  **(+ / −)** | **Perm. p**  **maxZ-56**  **(+ / −)** |
| --- | --- | --- | --- | --- | --- | --- | --- |
| Default-mode – Default-mode | 4095 | 171 | 0 | 4.18 | 0.00 | 0.0029 / 0.5635 | 0.1558 / 1.0000 |
| Default-mode – Somatomotor | 7007 | 16 | 0 | 0.23 | 0.00 | 0.8915 / 0.1730 | 1.0000 / 1.0000 |
| Default-mode – Visual | 5551 | 16 | 1 | 0.29 | 0.02 | 0.9186 / 0.2616 | 1.0000 / 1.0000 |
| Default-mode – Frontoparietal control | 4732 | 104 | 0 | 2.20 | 0.00 | 0.0240 / 0.5715 | 0.5420 / 1.0000 |
| Default-mode – Ventral attention/salience | 4277 | 17 | 0 | 0.40 | 0.00 | 0.8633 / 0.4197 | 1.0000 / 1.0000 |
| Default-mode – Dorsal attention | 4186 | 8 | 0 | 0.19 | 0.00 | 0.9502 / 0.3030 | 1.0000 / 1.0000 |
| Default-mode – Limbic | 2275 | 51 | 0 | 2.24 | 0.00 | 0.0184 / 0.5265 | 0.5877 / 1.0000 |
| Somatomotor – Somatomotor | 2926 | 7 | 0 | 0.24 | 0.00 | 0.8536 / 0.3347 | 1.0000 / 1.0000 |
| Somatomotor – Visual | 4697 | 3 | 0 | 0.06 | 0.00 | 0.9351 / 0.2719 | 1.0000 / 1.0000 |
| Somatomotor – Frontoparietal control | 4004 | 8 | 2 | 0.20 | 0.05 | 0.8886 / 0.1244 | 1.0000 / 1.0000 |
| Somatomotor – Ventral attention/salience | 3619 | 12 | 0 | 0.33 | 0.00 | 0.8588 / 0.4057 | 1.0000 / 1.0000 |
| Somatomotor – Dorsal attention | 3542 | 11 | 3 | 0.31 | 0.08 | 0.8764 / 0.2253 | 1.0000 / 1.0000 |
| Somatomotor – Limbic | 1925 | 1 | 0 | 0.05 | 0.00 | 0.9310 / 0.3415 | 1.0000 / 1.0000 |
| Visual – Visual | 1830 | 0 | 2 | 0.00 | 0.11 | 0.9365 / 0.3913 | 1.0000 / 1.0000 |
| Visual – Frontoparietal control | 3172 | 6 | 1 | 0.19 | 0.03 | 0.9202 / 0.2375 | 1.0000 / 1.0000 |
| Visual – Ventral attention/salience | 2867 | 2 | 0 | 0.07 | 0.00 | 0.9509 / 0.4049 | 1.0000 / 1.0000 |
| Visual – Dorsal attention | 2806 | 4 | 2 | 0.14 | 0.07 | 0.9355 / 0.2904 | 1.0000 / 1.0000 |
| Visual – Limbic | 1525 | 2 | 0 | 0.13 | 0.00 | 0.9179 / 0.3213 | 1.0000 / 1.0000 |
| Frontoparietal control – Frontoparietal control | 1326 | 5 | 1 | 0.38 | 0.08 | 0.8642 / 0.3932 | 1.0000 / 1.0000 |
| Frontoparietal control – Ventral attention/salience | 2444 | 12 | 1 | 0.49 | 0.04 | 0.7717 / 0.3148 | 1.0000 / 1.0000 |
| Frontoparietal control – Dorsal attention | 2392 | 5 | 2 | 0.21 | 0.08 | 0.9318 / 0.3377 | 1.0000 / 1.0000 |
| Frontoparietal control – Limbic | 1300 | 8 | 1 | 0.62 | 0.08 | 0.4990 / 0.3129 | 1.0000 / 1.0000 |
| Ventral attention/salience – Ventral attention/salience | 1081 | 19 | 1 | 1.76 | 0.09 | 0.0458 / 0.3109 | 0.8331 / 1.0000 |
| Ventral attention/salience – Dorsal attention | 2162 | 19 | 0 | 0.88 | 0.00 | 0.1805 / 0.4712 | 1.0000 / 1.0000 |
| Ventral attention/salience – Limbic | 1175 | 0 | 0 | 0.00 | 0.00 | 0.9359 / 0.3700 | 1.0000 / 1.0000 |
| Dorsal attention – Dorsal attention | 1035 | 8 | 1 | 0.77 | 0.10 | 0.3577 / 0.4185 | 1.0000 / 1.0000 |
| Dorsal attention – Limbic | 1150 | 0 | 0 | 0.00 | 0.00 | 0.9438 / 0.3204 | 1.0000 / 1.0000 |
| Limbic – Limbic | 300 | 3 | 0 | 1.00 | 0.00 | 0.2143 / 0.3853 | 1.0000 / 1.0000 |

*Note. Learning is biased-dimension rare-minus-frequent RT; pooled correlations are group centered. CIs are approximate pointwise 95% Fisher-z intervals. All 28 pairs are retained within each pooled, SS and DS Holm-28 family. Panel D counts pooled raw two-sided p < .01 edges among 79,401 possibilities; percentages use each pair’s possible-edge capacity. maxZ-56 uses all 28 pairs × both signs within learning.*
